# Multifaceted quality assessment of gene repertoire annotation with OMArk

**DOI:** 10.1101/2022.11.25.517970

**Authors:** Yannis Nevers, Victor Rossier, Clément Marie Train, Adrian Altenhoff, Christophe Dessimoz, Natasha Glover

## Abstract

Assessing the quality of protein-coding gene repertoires is critical in an era of increasingly abundant genome sequences for a diversity of species. State-of-the-art genome annotation assessment tools measure the completeness of a gene repertoire, but are blind to other types of errors, such as gene over-prediction or contamination.

We developed OMArk, a software relying on fast, alignment-free sequence comparisons between a query proteome and precomputed gene families across the tree of life. OMArk assesses not only the completeness, but also the consistency of the gene repertoire as a whole relative to closely related species. It also reports likely contamination events.

We validated OMArk with simulated data, then performed an analysis of the 1805 UniProt Eukaryotic Reference Proteomes, illustrating its usefulness for comparing and prioritizing proteomes based on their quality measures. In particular, we found strong evidence of contamination in 59 proteomes, and identified error propagation in avian gene annotation resulting from the use of a fragmented zebra finch proteome as reference.

OMArk is available on GitHub (https://github.com/DessimozLab/OMArk), as a Python package on PyPi, and as an interactive online tool at https://omark.omabrowser.org/.

## Introduction

Ambitious sequencing initiatives around the world have already begun to release chromosome-level genome assemblies of an unprecedented number of species, covering a wider range of earth biodiversity than ever before^1–3^. These additional species from diverse taxa will drastically improve the power of comparative genomics methods and more genome sequences will undoubtedly lead to a more accurate inference of when and how genes and species evolved^4^. However, such a goal can only be reached if the available data are a good reflection of biological reality, which requires extensive quality control.

Robust quality standards for genome assembly have been defined as part of sequencing initiatives, but there is a need for better standards and metrics of genomic features, in particular protein-coding genes^5^. These standards include: 1) whether the repertoire is complete; 2) the proportion of genes that have been correctly annotated; 3) whether there is erroneous annotation of non-coding sequences as protein-coding genes; and 4) whether exogenous genomic sequences, resulting from contamination, are present in the data. A few methods, based on conserved gene markers, can be used to measure the completeness of a gene repertoire (e.g. BUSCO^6,7^, EukCC^8^, DOGMA^9^) and to some extent contamination from other species (e.g. EukCC). Other indicators of quality, such as the UniProt Complete Proteome Detector, flag annotations with an unexpected number of protein-coding genes^10^. However, key aspects of proteome quality are not covered by existing methods, especially estimates of artifactual annotation, which may be more common than expected in publicly available genomes^11^.

Here, we present OMArk, a novel method for proteome quality assessment based on fast placement of protein sequences within known gene families. OMArk compares the query species’ inferred gene families to the expected families of its lineage and outputs multiple complementary quality statistics (Figure 1). First, OMArk gives an estimate of the query proteome’s completeness, based on the proportion of expected ancestrally conserved genes, similar to BUSCO but considering conserved multi-copy genes as well. Second, it gives an estimated proportion of protein sequences that are placed in known gene families from the same lineage (taxonomic consistency). Those that are not may be contaminant or erroneous sequences—with contamination being recognisable by a concentration of placement into particular secondary lineages. Thus, OMArk assesses proteome quality not only by measuring what is expected to be there, but also by evaluating what is *not* expected to be there. This feature is, to our knowledge, not provided by any other existing methods. We demonstrate OMArk’s ability to accurately estimate multiple quality metrics on proteomes with artificially introduced errors as well as on real use cases.

**Figure 1.**
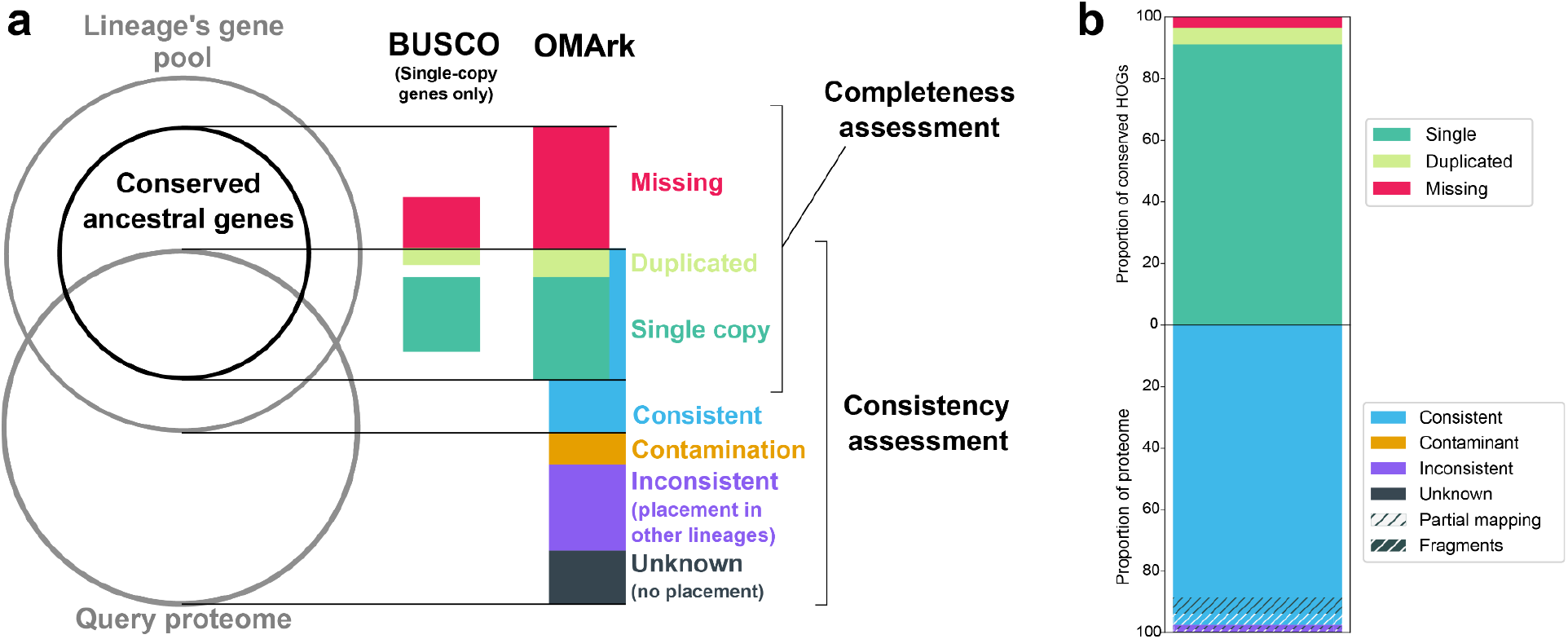
Summary of OMArk proteome quality statistics. **a) Schematic overview of the OMArk concept and output.** OMArk provides two main quality assessment categories: **1) Completeness assessment**, based on the overlap of the query proteome with a conserved ancestral gene set of the species’ lineage. OMArk classifies genes in the query proteome which are found in *Single copy*, multiple copies (*Duplicated*), or *Missing*. This is similar to methods like BUSCO, but also considers conserved genes which are in multiple copies. **2) Consistency assessment**, based on the proportion of query proteins placed in gene families of the correct lineage (Consistent), gene families of an incorrect lineage (either randomly (*Inconsistent*) or to specific species (*Contamination*)), and query proteins placed in no gene families at all (*Unknown*). **b) An example of OMArk’s graphical output for the model organism zebrafish** (*Danio rerio)*. The top of the stacked bar plot represents the completeness assessment: genes found in Single-copy (dark green; here: 90.89%), Duplicated (light green, 5.55%) and Missing genes (red, 3.56%). The lower part of the bar plot represents the consistency assessment: taxonomically Consistent genes (blue, 96.96%), taxonomically Inconsistent genes (violet, 1.92%), Contaminants (orange; none in this example), or with no detected homology (Unknown; black, 1.12%). All categories annotated with hashes correspond to the proportion of *Partial mappings* (black hashes) and *Fragmented* genes (white hashes).

## Results

### Software overview

OMArk is available as an open source command-line tool and as a web server. The command-line tool is distributed as a python package on PyPI and on GitHub (https://github.com/DessimozLab/OMArk). The OMArk pipeline is easy to install and running it locally requires few dependencies. It needs only a precomputed OMAmer database, which is available for download on Zenodo and on the OMA Browser^12^. The web server (https://omark.omabrowser.org) lets users upload a FASTA file of their proteome of interest and visualize or download the results once the computation is done, on average in 35 minutes for a proteome of 20,000 sequences.

#### Query protein placement

OMArk takes as input a protein sequence file where each gene is represented by at least one protein sequence it codes for. The OMArk pipeline starts by running OMAmer^13^, an ultra-fast *k-mer*-based method that assigns proteins to gene families and subfamilies (Figure 2a), represented as Hierarchical Orthologous Groups (HOGs)^14^. It exploits predefined gene families for 2,500 species available from the OMA database^12^, but could in principle be used on any other database using the HOG concept to represent them.

**Figure 2.**
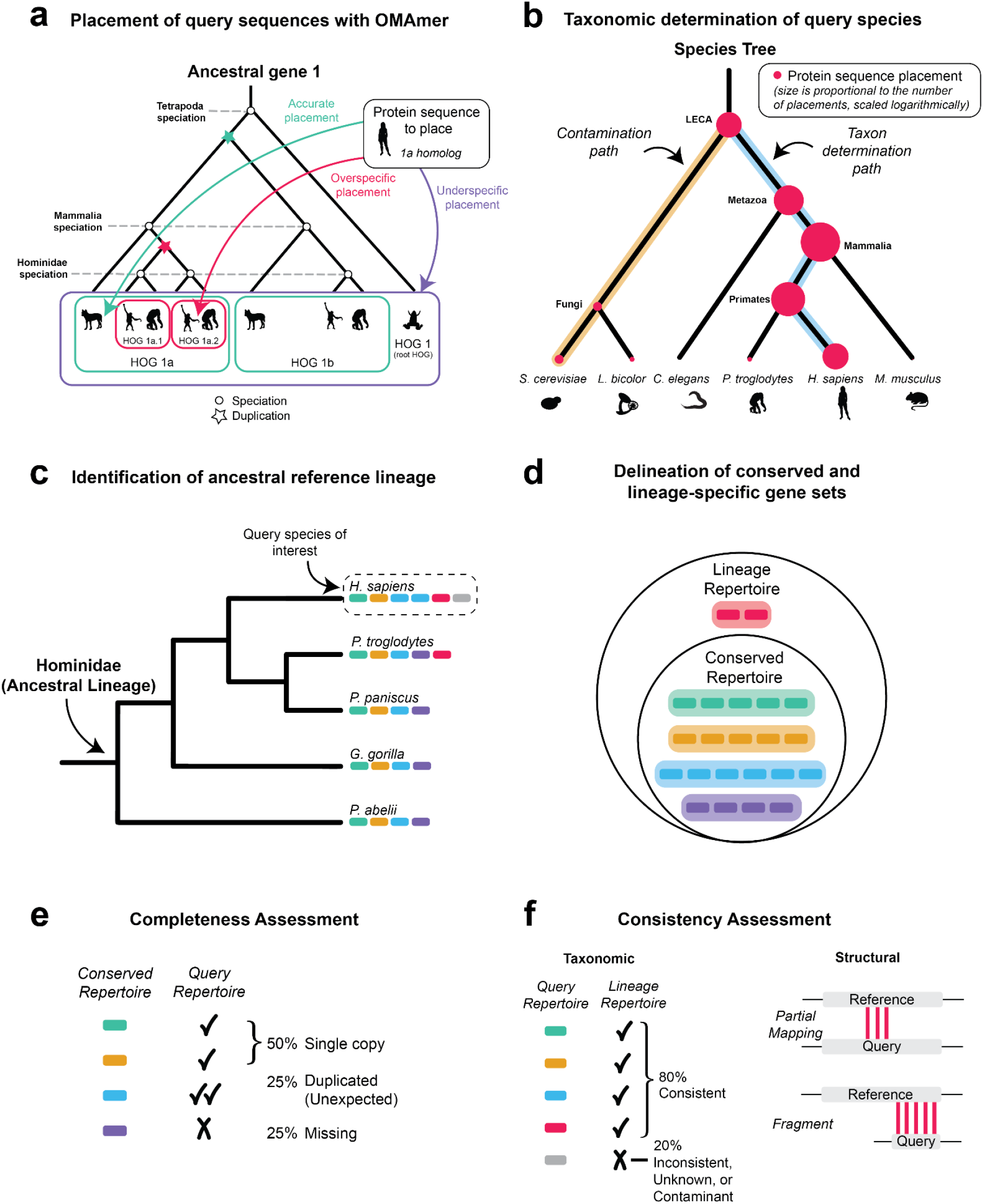
Overview of the OMArk methodology. **a)** Sequences from the query proteome are placed into known Hierarchical Orthologous Groups (HOGs) using the *k-mer* based fast mapping method OMAmer. Shown is a gene tree with the nested gene families (HOGs), delineated by speciation and duplication events. OMAmer provides accurate placement of protein sequences in their correct subfamily. **b)** The specific taxon of the query species is automatically determined by OMArk. Here, the species tree is shown, with protein placements represented by red dots. The size of the dots are logarithmically proportional to the number of placements in a typical scenario, but simplified for this schema. The path to the query taxon is inferred based on the maximal number of placements (blue) and the path(s) to contaminant taxa are determined as those with more placements than expected by chance. **c)** OMArk defines the ancestral reference lineage for a given query species as the most recent taxonomic level including the species and represented by at least five species in the OMA database. Here, a species tree is shown with colored bars representing individual genes. **d)** The conserved and lineage-specific gene sets are defined. The conserved repertoire contains all the HOGs defined at the reference ancestral level which cover at least 80% of the species in the clade. These are gene families inferred to be present since the common ancestor. The lineage repertoire is a superset of the conserved repertoire, with the addition of genes which originated later in the lineage and are still present in at least one species in the OMA database. In the repertoires, genes from the different species are grouped into their HOGs. **e)** OMArk assesses completeness by comparing the conserved ancestral repertoire to the query protein sequences and classifying them as Single copy, Duplicated, or Missing. **f)** OMArk assesses consistency by comparing the query protein sequences to the lineage repertoire and classifying them as taxonomically Consistent, Inconsistent, Unknown or Contaminant. It also does a gene model structure assessment by classifying them as Partial or Fragment.

#### Species identification

OMArk exploits the query protein’s placement into gene families and the knowledge of the taxon in which the gene families emerged to infer the species composition of the proteomes (Figure 2b). In a perfect case, a proteome from a given species will have placements in gene families that emerged at every taxon the species belongs to and no other. For example, human genes will have originated at the common ancestor of primates, mammals, vertebrates, but *not* rodents. OMArk starts from this assumption and identifies paths in the taxonomic tree where placements are overrepresented. It then selects the most recently emerged clade as the inferred taxon. If more than one path is overrepresented, OMArk will report the one with the highest number of placements as the main taxon, and all others as possible contaminants.

#### Ancestral reference lineage identification

Based on the main taxon placement, or a taxonomic identifier proposed by the user, OMArk then selects an **ancestral lineage**: the taxon closest to the leaves that contains the species of interest and at least five species in the OMA database (Figure 2c).

#### Completeness assessment

From the OMAmer database, OMArk selects all gene families that were present in the common ancestor of the ancestral lineage and still are represented in at least 80% of its extant species (Conserved Repertoire; Figure 2d). Because most of these genes are expected to be present in the query proteome, completeness of this gene family set is used as a proxy of proteome completeness. OMArk reports the proportion of the conserved gene families that are found in the query proteome in Single copy, multiple copies (Duplicated) or are Missing (Figure 2e). An incomplete proteome would have a high proportion of Missing gene families. Contrary to BUSCO, the selected conserved genes are not necessarily expected to be in single copy in extant genomes although they were likely in a single copy in the lineage’s ancestor. Thus, the Duplicated category is divided into subcategories to indicate if it corresponds to a known duplication that occurred after the ancestral speciation of the relevant lineage (Expected) or not (Unexpected). An example of the OMArk output for the *Danio rerio* proteome shows it has a high level of completeness, as expected from a model species (Figure 1).

#### Consistency assessment

The main novelty of OMArk is that it provides an assessment of the consistency of all the genes in the query proteome compared to what is known for its lineage, both taxonomically and structurally.

#### Taxonomic consistency

classifies proteins of the query proteome based on their taxonomic origin by comparing it to the set of all gene families that are known to exist in its lineage (Lineage Repertoire; Figure 2d). The proportion of the query proteome that is placed into this set is classified as Consistent, whereas the proportion of genes associated to other gene families is labeled as either Inconsistent or Contaminant (Figure 2f). The Contaminant categories will contain all Inconsistent mappings that are closer to a contaminant species than to the main species, as determined in the protein placement step. The proportion of proteins with no gene family assignment are simply labeled as Unknown.

#### Structural consistency

classifies proteins of the query proteome based on sequence feature comparisons with their gene family. The proteins that only share *k-mers* with their gene families over part of its sequence are classified as Partial mappings, and the proteins whose lengths are less than half the median length of their gene families are categorized as Fragments (Figure 2f).

These two complementary parts of the consistency assessment, performed over the whole coding-gene repertoire, are invaluable in identifying annotation artifacts and are missing in most existing quality assessment methods. A high proportion of Consistent proteins for a given proteome gives a high confidence on the correctness of the annotation, whereas the proportion of Partial mappings and Fragments can help identify cases where the gene models may be incorrect or incomplete. A high proportion of Inconsistent proteins, especially if they are mainly Partial or Fragments, however, could point to entirely erroneous genome annotation. As shown in Figure 1, the *Danio rerio* proteome is mostly composed of taxonomically and structurally consistent genes.

### Validation on simulated proteomes

In order to evaluate the ability of OMArk to provide accurate statistics for quality assessment, we simulated cases of genome incompleteness, presence of erroneous sequences, and cross-species contamination. We used two datasets of eukaryotic proteomes for our simulations. First, a set of nine model species whose quality is expected to be high (‘model dataset’). Second, a set of 16 species representing the diversity of Eukaryotes and absent from the original OMA database, which mirrors the expected use case (‘representative dataset’). The datasets are listed in Supplementary Table 1.

The completeness simulation was performed for each proteome in the datasets by randomly removing different percentages of genes from the proteome and running OMArk on the reduced repertoire. We show that OMArk retrieves close estimates of the simulated completeness in most cases, with a tendency to overestimate completeness (Figure 3a). The error margin is lower in the model dataset (+2.6% on average) than in the representative dataset (+8.8% on average). Its performance is similar to BUSCO, which overestimates completeness by a slightly lower margin (+2.1% and +6.1% on average for model and representative sets, respectively). Both methods overestimate completeness in species with higher numbers of duplicated genes, which explains the higher error rate in the representative set. OMArk is likely more sensitive to duplication because it considers all conserved gene families, not only those in single-copy, like BUSCO.

**Figure 3.**
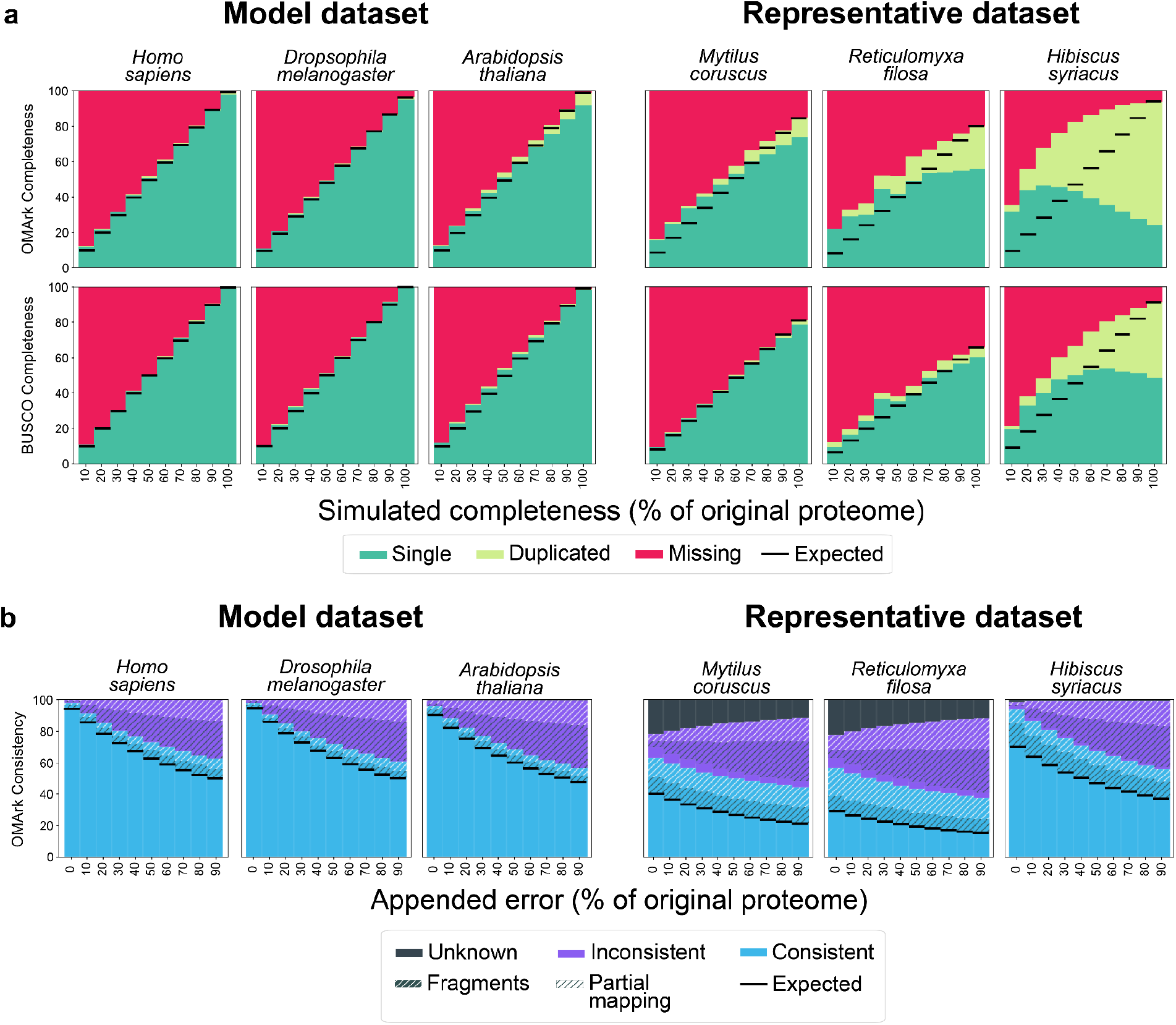
OMArk results for simulated proteomes. **a) Simulated completeness.** OMArk (top) and BUSCO (bottom) results for three example species of the model dataset (left) and the representative dataset (right). Colors represent the part of the conserved gene set found in Single-copy (green), Duplicated (light green) or Missing (red). The simulated completeness corresponds to the percentage of the genome that has been randomly selected in each simulation. Dotted horizontal lines show the expected completeness given the measured completeness for the source proteome. **b) Erroneous sequence simulation**. Colors represent proteins which map to the correct lineage (Consistent, blue), to another lineage (Inconsistent, violet) or have no homologs (Unknown, black). Hashes indicate structural inconsistency relative to the gene family; either Partial mapping (black hashes) or Fragmented genes (white hashes). The appended error (x-axis) corresponds to the quantity of erroneous sequences that was added to the proteome, as a percentage of its original protein number. Dotted horizontal lines indicate the expected number of structural and taxonomically Consistent genes, considering the proportion in the source proteome and the known introduced error.

The erroneous sequence simulation was done by adding various amounts of randomly generated sequences to each proteome. The addition of erroneous sequences led to a higher proportion of Inconsistent genes, and to a lesser extent, higher proportions of genes in the Partial mapping and Fragment categories of the Consistent genes (Figure 3b). Interestingly, the proportion of Unknown genes, which is higher in the representative dataset (likely because of low clade sampling in OMA) also decreases with introduced error. This is because randomly generated sequences still get placed into gene families but with low sequence similarity and coverage. Notably, the proportion of taxonomically and structurally Consistent genes detected by OMArk is equal to the expected value in all simulations, indicating that this category gives an appropriate estimate of the proportion of high-confidence coding sequences.

The contamination simulation was done by introducing sequences from other microbial species (bacteria, fungi, microbial eukaryotes) or humans to the model and representative datasets. OMArk was accurate in detecting the taxonomic origin of the contaminant with variable degrees of sensitivity. For bacterial and fungal species, contamination could be detected from only ten contaminant proteins in some cases, and reliably with 50 contaminant proteins. It was less accurately detected when it came from other eukaryotic species (detected in most cases starting from 100-200 proteins). OMArk was unable to detect contamination when the contaminant species had no close relatives in our reference database (i.e. *Stentor coeruleus* and *Planoprotostelium fungivorum*), or when the contaminant proteome was closely related to the contaminated proteome (Supplementary Table 3; detailed analysis for all simulations in Supplementary Data).

### OMArk results for 1805 Eukaryotic reference proteomes

Comparing protein-coding gene annotations between closely related species, including one gold standard, is essential to assess annotation quality^5^. For this reason, we ran OMArk on a set of 1805 Eukaryotic UniProt proteomes, which serves as a reference dataset (Figure 4; Supplementary Table 3). A description of the quality assessments for major clades and of particular cases with low quality results are provided in the Supplementary material. We make all these results freely available on the OMArk webserver, where users can easily compare the quality assessment of their query proteome to closely related species through the “Overview” button on the result page.

**Figure 4.**
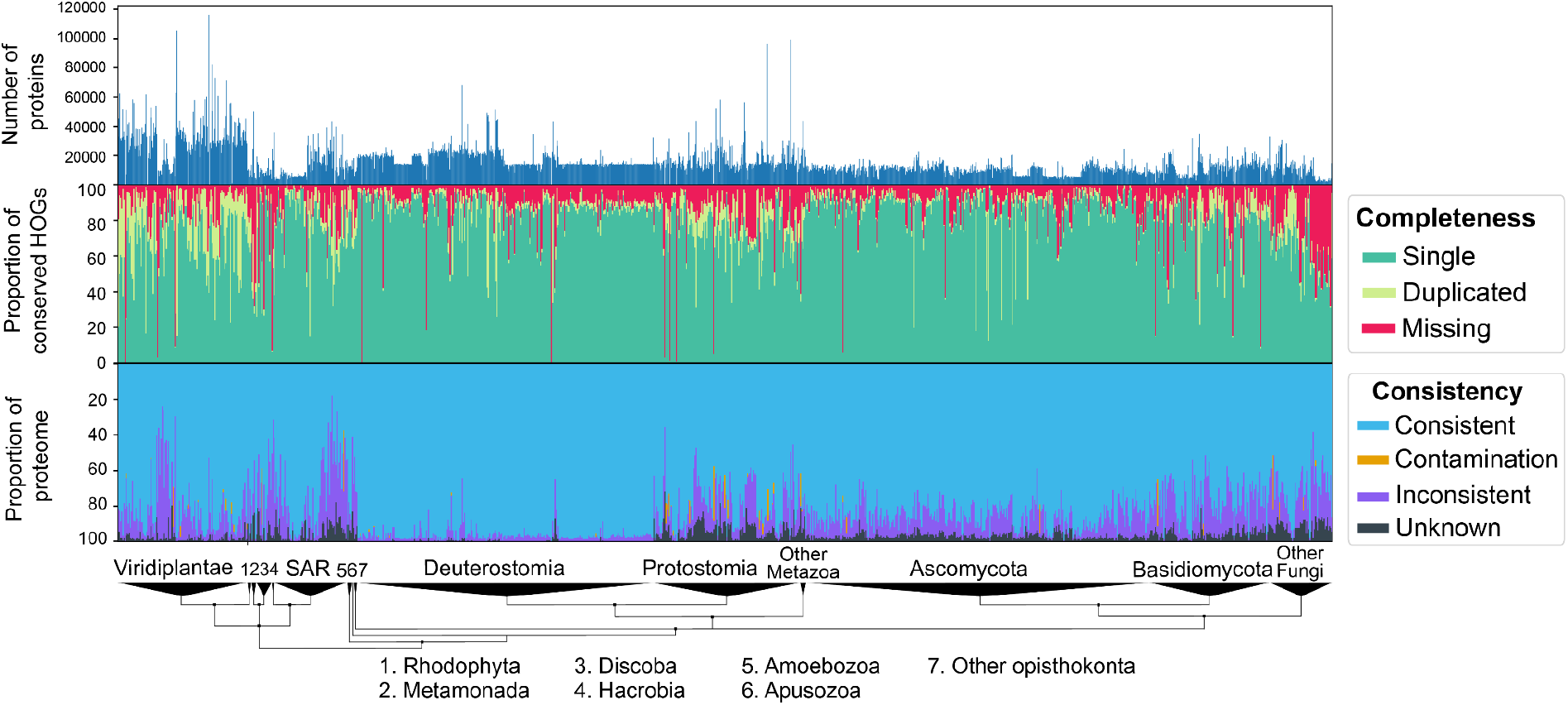
OMArk results on 1805 Eukaryotic proteomes from UniProt. Top - bar graphs of the number of canonical proteins in each proteome. Middle - Completeness assessment: Proportion of conserved genes (green) in the proteome with a breakdown between Single Copy (light green), Duplicated (dark green), and Missing (red) in all proteomes. Bottom - Consistency assessment: Proportion of accurately mapped proteins (Consistent; blue), incorrectly placed proteins (Inconsistent; purple), contaminant proteins (orange) and proteins with no mapping (Unknown; black). Genomes are ranked by taxonomy, with major eukaryotic taxa shown on a taxonomic tree at the bottom.

### Contamination in public databases

Out of the 1805 UniProt Eukaryotic genomes, OMArk detects 115 contamination events in 72 proteomes, with several proteomes contaminated by more than one species (list in Supplementary Table 4). Two of them, *Ricinus communis* and *Lupinus albus* were found to be contaminated by eight species each (mostly Bacteria and Fungi) indicating that extreme cases of contamination persist in public databases. For all contamination cases, we performed independent validation through BLAST queries and public results from BlobToolKit Viewer (Supplementary Table 4). We confirmed 100 of the detected contamination events in 59 species. An additional four corresponded to likely ancient HGT events or chloroplastic acquisition that could not be detected as such due to incomplete sampling in our database.

Finally, two of the 11 false positives were due to contamination in the proteomes included in our source database: contamination from *Saccharomyces cerevisiae* in the proteome of *Drosophila biarmipes* and from a fungal species in the proteome of *Picea glauca*. This illustrates the risk of exploiting contaminated proteomes for comparative analysis. These proteomes will be removed in the next iteration of the OMA database.

### Detection of error propagation in avian proteomes

We detected widespread presence of fragmented genes in the 234 avian species from the UniProt eukaryotic dataset (median proportion of Taxonomically Consistent Fragments: 16.9%, standard deviation: 4.29). However, this was not the case for the most studied bird species, such as chicken [*Gallus gallus*] (proportion of Taxonomically Consistent Fragments: 2.4%) (Supplementary Figure 7). The proportions of fragments depended mainly on the source of the proteome. Most of the highly fragmented proteomes originated from the same source, the Bird 10k consortium annotation pipeline, and tended to have fragments in the same gene families, suggesting systematic bias in the annotation process (Supplementary Figures 8 and 9, detailed analysis in Supplementary Material). Annotations of protein-coding genes for these genomes were partly done using homology from the Ensembl 85^15^ annotation of zebra finch (*Taeniopygia guttata*). Using OMArk, we also detected a high proportion of fragments in this version of the zebra finch proteome (proportion of Taxonomically Consistent Fragments: 12.47%). Furthermore, on average, more than half of the genes fragmented in the Bird 10k proteomes were also detected as fragments in this specific zebra finch proteome (Supplementary Figure 10). These results suggest fragments in these proteomes likely result from error propagation.

### Use case: selection of high-quality proteomes among closely related species

OMArk’s quality assessment is dependent on the selected ancestral lineage used to define the conserved gene repertoire. Thus, a best practice for OMArk is to compare the results to species sharing the same reference lineage. We illustrate this application by comparing OMArk results of a model species–*Mus musculus*–to its closely related species in the UniProt dataset, Myomorpha, a clade of mouse-like rodents (Figure 5a). As expected, the well curated model species *Mus musculus* and *Rattus norvegicus* are the best scoring in this taxonomic division both in completeness and consistency. Several other species in the clade exhibit noticeable quality issues, even when they are represented in the OMA database and thus contributed to the ancestral reference HOGs (e.g *Cricetulus griseus*). Results for other model organisms are comparable, with the model species consistently ranking as the better quality proteomes in their respective clade (detailed description in Supplementary Material, Supplementary Figures 11-19).

**Figure 5.**
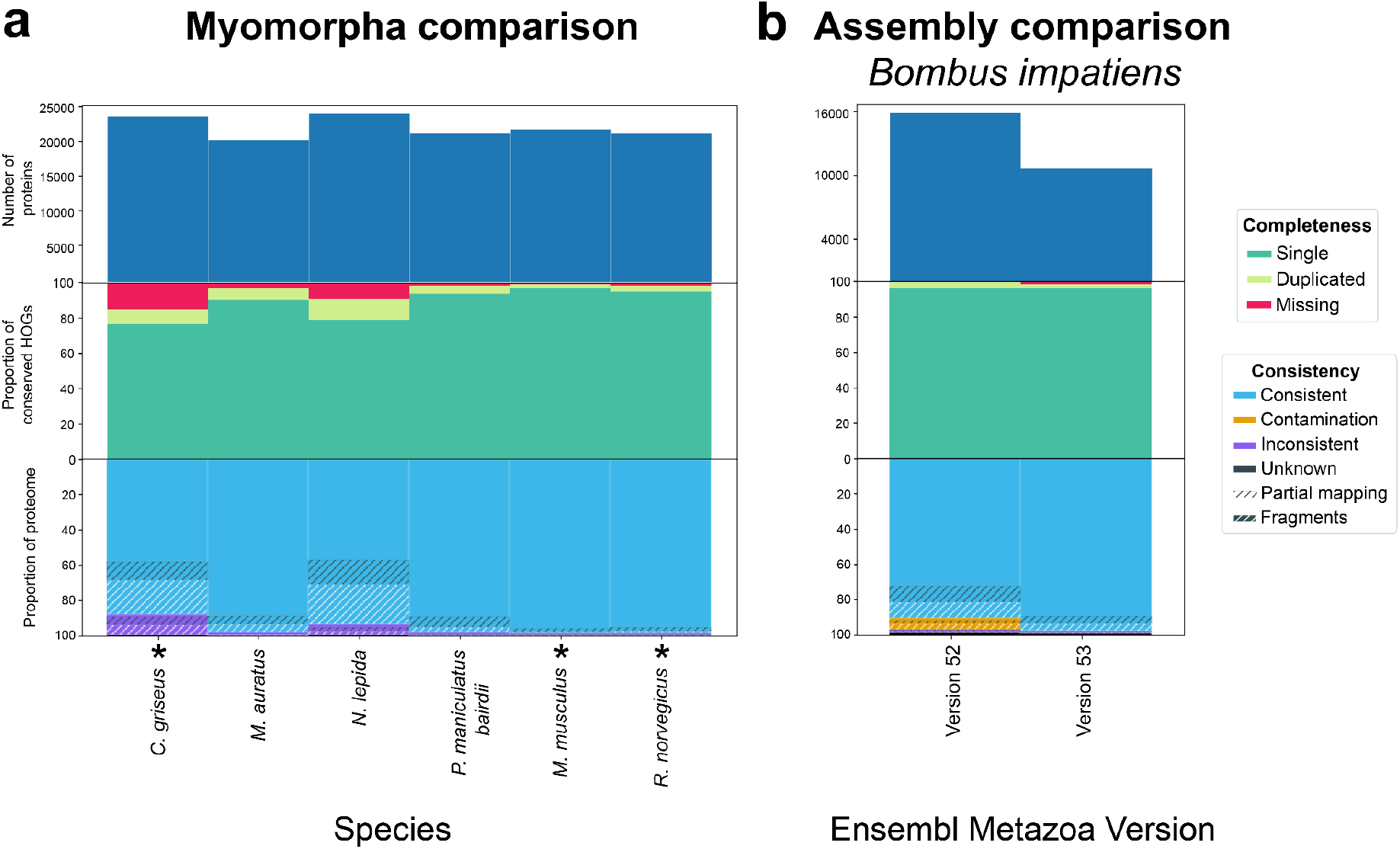
OMArk comparisons between closely related species within a clade or assembly versions. **a**. OMArk results for different Myomorpha proteomes. The species currently in OMA which contributed to the ancestral reference lineage gene families are shown with an asterisk. Mouse (*Mus musculus*) and the Norwegian rat (*Rattus norvegicus*) stand out as high quality annotations. **b**. OMArk results for *Bombus impatiens*, comparing two different versions of the assembly. The newest assembly (version 53) shows an improvement in gene set and contamination removal, with a slight decrease in completeness.

This shows how OMArk can be used to identify the best quality protein dataset in any taxon of interest and thus could be used to select a representative genome for this clade. Furthermore, it may help guide efforts in generating better quality genomic data for non-model species.

### Use case: assembly and annotation version comparisons

Another use case for OMArk is the comparison of gene repertoires inferred from the same species, either from different assemblies or from different annotation methods. This may be used to assess improvement in gene repertoire completeness or in consistency between dataset versions for the same species. It may also be used to benchmark annotation methods for a genome of interest. We perform an example comparison by running OMArk on newer vs. older assemblies or annotations for species with documented changes in the last two releases of Ensembl Metazoa (53-54)^16^. This corresponds to 11 protostome species with different assembly versions and seven nematode species with changes in the protein-coding gene annotation (Supplementary Table 5).

When comparing OMArk results across different annotation versions of the same assembly, the results only show minor changes. This is expected since these changes were likely iterative and did not concern many genes. However, we still detect a general trend of removal of Duplicated conserved genes and an increase in the proportion of Taxonomically and Structurally Consistent genes (details in Supplementary Material).

Between different assemblies, OMArk detects improvement in completeness and/or in structurally and taxonomically consistent genes for all but one species (detailed results in Supplementary Material). Interestingly, improvements were not always observed in both categories. In two species (*Bombus impatiens* (Figure 5b) and *Acyrthosiphon pisum*), we detected a small decrease in completeness, but a large improvement in terms of taxonomic and structurally-consistent proteins. In another species, *Crassostrae gigas*, we see the opposite, demonstrating the value of considering both measures at the same time. Finally, OMArk allowed us to detect the removal or decrease in contamination for four species (*Bombus impatiens, Schistosoma mansoni, Acyrthosiphon pisum* and *Glossina fuscipes*), as well as one species where contaminants were introduced in the newest assembly (*Teleopsis dalmanni*).

Together, these results show that OMArk can provide a quantitative characterization of proteome quality which would help scientists choose the best annotation method for the same species. It also provides a more exhaustive measure of change between assembly and annotation versions than completeness alone.

## Discussion

We introduced OMArk, a method that exploits the OMA database and *k-mer*-based fast gene family placement to assess the quality of a protein-coding gene annotation. Our results on simulated incomplete genomes and on real proteomes showed that the completeness measured by OMArk is comparable to BUSCO, another method often used for this purpose. This is not surprising, as the core principle behind both methods is the assessment of the presence or absence of near universally conserved genes in a lineage. However, there are several key differences. OMArk does not only focus on single-copy conserved genes, but also includes gene families that underwent duplication in the lineage leading to the query species. Second, BUSCO uses Hidden Markov model profiles to map query genes to their conserved gene families, a method more accurate but slower than the *k-mer* mapping exploited by OMArk. Finally, OMArk does not rely on a prespecified dataset of conserved genes and is able to automatically choose a set of conserved genes depending on the query species’ taxonomic lineage.

A major novelty of OMArk is its global assessment of proteomes based on a broader selection of orthologous groups than the universally conserved ones. While these cannot be used for estimating completeness because they are not expected to be conserved, we can still use them to make estimates about the quality of gene models reported in a protein-coding gene set. Proteins placed into gene families that are taxonomically consistent with the species of interest can be more confidently considered as *true* coding genes. Moreover, we can give an estimate of the quality of their gene structure by comparing to known sequences in the same family. However, there are a few caveats when interpreting OMArk results: consistency with genes from the same lineage is expected only in species where the main mode of gene family inheritance is vertical and if the chosen family is well sampled and of good quality in our reference database. Our reference database, OMA, is actively maintained and has a 6-month release cycle where we seek to improve coverage for underrepresented species and include only high-quality data.

Finally, OMArk reports possible contamination based on the taxonomic distribution of taxonomically inconsistent gene families. The contamination indicator provided by OMArk should help genome annotators to flag these cases before publication and potentially reassess their genomic data. OMArk can not discriminate between contamination and recent Horizontal Gene Transfer events. However, using the list of potential contaminants, annotators could identify the corresponding contigs in the genome assembly and either validate them or not as a likely part of the genomic sequence. Nevertheless, in the case of contamination detection by OMArk, we recommend users to use assembly-level dedicated methods^17^, such as BlobToolKit^18^, to perform an in-depth analysis and correction of the genome assembly.

By quickly providing a more comprehensive assessment of proteome quality than existing methods, OMArk will help annotators to improve gene annotation and users to select species for their investigations, where the quality of the gene repertoire is often critical. We plan to report the proteomes of lesser quality identified by our methods to UniProt and work with public databases to help improve existing gene sets.

## Methods

### OMAmer placement

The OMAmer database used in this study and provided with the paper was generated from the December 2021 release of the OMA database^12^. Root HOGs with five or less proteins and a species coverage (the proportion of species with a gene) lower than 0.5 were excluded, as they are most likely spurious.

### Overview of the OMArk algorithm

All analyses shown here were performed with OMArk version 0.2.1. The OMArk software takes the following as minimum input: 1) the output of the OMAmer placement for a whole proteome, whereby proteins of these proteomes are placed in HOGs, and 2) the path to the corresponding OMAmer database.

Optionally, OMArk can take the NCBI taxonomy ID of the proteome’s species which will be used to select its ancestral lineage, otherwise its taxa will be inferred automatically (see species identification below). The FASTA file of the query proteome is also an optional input, which may be used to generate output FASTA files for Inconsistent, Contaminant, and Unknown proteins. Finally, if the proteome contains multiple isoforms per gene, an additional option (-i) allows the user to provide a comma-separated file where all protein identifiers corresponding to a single gene are written on the same line. Only one isoform will be selected for completeness and consistency assessment, based on the OMAmer placement score as detailed in “Isoform selection” below.

#### Isoform selection

If the target proteome contains more than one protein by gene, and an isoform file was provided by the user, OMArk will automatically select the sequence with the best match in the OMA database, based on the hit’s “family score” (from OMAmer). This helps ensure that gene model comparison will happen between similar isoforms. To do this, OMArk will select the isoform *I* with the highest isoform score *I*s, calculated according to the formula:

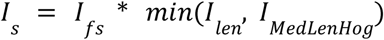

where *I*_*fs*_ is the family score of the isoform, *I*_*len*_, is its length in amino acids, and *I*_*MedLenHog*_ is the median length of the family it matches to.

#### Automatic species identification and contamination assessment

The taxonomic distribution of HOGs in the query proteome can be used to automatically detect the species from which they come from. OMArk does this by using the non-redundant list of HOGs in which proteins of the query proteomes were placed, and extracting the taxonomic level where each HOG is defined (i.e. the taxonomic node after emergence or duplication of the gene family). This is used to obtain the number of mappings to each clade in the tree of life which we call **clade occurrence *N***.

To reduce noise due to incorrect mapping, which is more common in broad clades with a large number of HOGs, we divide the clade occurrence by the number of total HOGs defined at each level to obtain a **normalized clade occurrence *N’***. In the presence of only one species and no noise, we assume the most likely placement would be the clade with the highest normalized clade occurrence, with all its parent clades having an equal or lower normalized occurrence count.

The OMArk algorithm uses this assumption and implements a few corrections to account for noise in HOG placement and allow for more than one species in the proteome, in case of contamination. First, all clades with an occurrence of more than 2 and that have a normalized occurrence of more than 0.001 are used to construct a simplified taxonomic tree containing only branches leading to these clades. The tree structure itself is derived from the OMA underlying taxonomy, which uses the NCBI taxonomy^19^.

The OMArk algorithm for species identification is a recursive postorder traversing function. At each leaf, it returns the leaf clade as likely placement, with occurrence *N*_*leaf*_ and normalized occurrence *N’*_*leaf*_ of the taxonomic level. At each node, it compares the occurrence scores of the current node to the most likely placements of its children. To be considered significant, a child’s placement has to satisfy:

- *N*_*child*_ > *N*_*node*_ * *D*_*node,child*_ where *D*_*x,y*_ represents the proportion of HOGs defined in clade *x* that have a child defined in clade *y*. A high value represents a high duplication rate in the branch leading from *x* to y. This condition controls for high duplication numbers in the branch leading to some lineages (e.g. ancestral whole genome duplication), which favor overspecific placement into those clades.
- *N*′_*child*_ > *N*′_*node*_ * |*S*_*child*_ |/|*S*_*node*_ where |*S*_*x*_| is the number of species in clade *x*. This condition controls for sampling imbalance in lineages which favor overspecific placement in larger clades.

If one child only is considered significant, it is returned as the most likely taxon. If more than one is considered significant, all are returned as likely taxa. If no child is considered significant, only the current node is returned as likely taxon. After traversal, this module outputs a list of independent clades which have more hits than expected by chance.

For each clade with more placements than expected, we select all proteins that can be unambiguously attributed to these clades: i.e. all proteins that map to a HOG defined at a node in the subtree leading to its first common ancestor with any other clade in the list. The clade with the most proteins is considered as the most likely main taxon, the other as contaminants. This possibly overestimates the proportion of contaminant sequences especially in presence of spurious hits (that would otherwise be in the Inconsistent category) but ensures most of the contaminants are included in the category.

#### Completeness assessment

The completeness assessment measures the proportion of HOGs that are expected to be conserved in the species’ lineage. This is done by first selecting the ancestral lineage of the species, defined as the most recent taxonomic level including the species and represented by at least 5 species in the OMA database. Then, OMArk defines the “conserved ancestral repertoire” of the query species: all the HOGs defined at this ancestral level which cover more than 80% of the species in the clade.

Since a HOG at the selected taxonomic level represents a single ancestral gene, conserved HOGs are classified as one of the following:

- **Single-copy** if one protein in the query proteome maps to it. To be robust to minor errors in phylogenetic placement, underspecific hits (placement in a parent HOG, Figure 2a) or a single overspecific hit (placement in a child HOG) are sufficient to consider a conserved HOG as single-copy.
- **Duplicated** if more than one query protein maps to it. A duplicated, conserved HOG is further classified as Unexpected if multiple proteins are all placed into the ancestral HOG itself (i.e., no evidence of such duplication exists in the OMA database) or as Expected if the proteins were placed into subfamilies of the HOG (i.e., the duplication event is documented in the database).
- **Missing** if no proteins in the query genome are placed into it.

#### Consistency assessment

The consistency assessment is used to evaluate the query proteome quality, again depending on the placement of its proteins into HOGs and the taxonomic level at which these HOGs are defined. Here, OMArk uses a “lineage repertoire” of the query species: all the HOGs from the conserved ancestral repertoire plus those that originated later on and are still present in at least one species of the lineage. It uses this lineage repertoire to classify proteins as:

- **Unknown** proteins are those that were not placed into existing HOGs. They correspond to either errors in the annotation or gene families with no detectable counterpart in OMA (due to falling in sparsely sampled clades or being a novel protein).
- **Consistent** proteins are those that were placed into a HOG consistent with the reference lineage: the HOG has a representative of at least one species from the lineage, whether it was present in the common ancestor of the reference lineage or emerged in its descendants.
- **Contaminant** proteins are those that map to a lineage consistent with the lineage of another species which has been detected as a likely contaminant by the contaminant detection module of OMArk (see “Automatic species identification and contamination assessment” in Methods).
- **Inconsistent** proteins are those that were placed into HOGs from other parts of the tree of life, and for which there is no evidence the gene families existed in the selected lineage or in any contaminant lineage. They are likely to be incorrect protein sequences or to correspond to gene families that were not observed in those species before.

For the proteins that map to existing HOGs, an additional characterization is provided:

- **Partial hits** are proteins from which less than 80% of the sequence overlaps with their target top-level HOG, that is, at least 20% of the sequence at the extremity of the protein has no k-mer in common with the proteins of the top-level HOG.
- **Fragments** are sequences which are not partial hits but whose length is <50% of the median length of sequences in the HOGs.

### Acquisition of proteome data

Reference proteomes were downloaded from UniProtKB^10^ on 10 February 2022 (version of 04/2021). Ensembl Metazoa proteomes were downloaded from their ftp website from version 52, 53 or 54 of the database (version number is reported in Supplementary Table 5).

### Generation of simulated proteomes

Two datasets were used to assess the effect of introducing errors into proteomes on OMArk quality scores. We worked on two datasets of real proteomes as the basis of the simulation: model proteomes and taxonomically representative proteomes. Model proteomes correspond to model eukaryotic species whose proteomes are assumed to be of high quality. Representative proteomes were selected under several criteria: they represent the major eukaryotic taxonomic divisions (2 of each when possible), they must not be present nor have species of the genus represented in the OMA database (to avoid circularity), and they must have the best score possible for the aspect of OMArk quality measures (mainly, few missing genes and a higher proportion of consistency from other species of their division). Both lists of species are available in Supplementary Table 1.

These source proteomes were manipulated in three ways, each simulating a case of artifactual annotation:

- *Missing genes*. For each proteome, only a fraction of the proteins were kept at random. This was repeated independently ten times with different proportions of the proteome kept, from 10% to 90% by increments of 10%.
- *Erroneous sequences*. Errors in genomic annotation were simulated from randomly generated nucleic sequences, from an equiprobable distribution of each base (25% of chance to draw A, T, G and C). The sequences were generated by increments of 3, representing codons in the Open Reading Frame, until a stop codon appeared. The resulting sequences were then translated into proteins and kept if their length was more than 20 amino acids. These sequences, independently generated in each simulation, were then added to each proteome proportionally to the original number of proteins in the original genomes, from 10% to 90% by increments of 10%.
- *Contamination*. A list of eukaryotic and bacterial proteomes, either from common contaminants in genomic data or microscopic species from a variety of clades were selected as contaminant proteomes. Then, a fixed number of proteins (10, 20, 50, 100, 200, 500 and 1000) were drawn randomly without replacement independently from the contaminant proteomes and added to the complete source proteomes.

### BUSCO comparisons

BUSCO^6^ v.5.2.2 was run on UniProt Reference Proteomes and simulated data, using the odb10 version of the BUSCO dataset of the most specific lineage possible covering the target proteome and with default options. The corresponding dataset is available in Supplementary Table 3. The summarized result for the BUSCO run was then extracted from the summary file.

### Avian proteome fragment analysis

To compare fragmented sets between all avian UniProt Reference Proteomes, we first queried the OMArk results for the proteins classified as lineage consistent fragments, then used the OMAmer placement file to obtain the gene families to which they were associated. To avoid biasing the comparisons in the presence of duplication, we associated each gene name to their whole gene family identifier (root HOG), rather than to their subfamilies. The overlap between fragmented gene set between two species was computed by directly comparing sets of their associated gene families using the following formula, for two sets A and B: 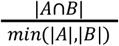

The denominator was chosen to be the cardinality of the smallest set in order to not underestimate overlap in the smallest sets.

The zebra finch proteome was downloaded from the Ensembl archive for version 85 of the database. Overlap with the zebra finch taxonomically Consistent Fragment set was done as above, but using the cardinality of the target proteome’s fragmented gene family set as the denominator.

### Additional analysis

The additional analyses were done in Python (v. 3.9.5) in a Jupyter Notebook. Plots were made using the matplotlib library (version 3.4.2)^20^.

## Data availability

OMArk is available on GitHub (https://github.com/DessimozLab/OMArk) and as a python package on PyPI. Precomputed results are made available through the OMArk webserver (https://omark.omabrowser.org). All datasets described here, Jupyter Notebooks used for the analysis, and Supplementary Table files are made available through Zenodo (doi:10.5281/zenodo.7359861).

## Acknowledgments

We thank Robert Waterhouse and Mark Blaxter for their helpful feedback and comments during the development process.

