## Supplementary Material for "Multifaceted quality assessment of gene repertoire annotation with OMArk"

**Supplementary Tables** 2

**Supplementary Results** 2

Simulation results 2

Incompleteness simulation 3

Erroneous sequences simulation 6

Contamination simulation 8

Results on UniProt Reference Proteomes 8

Global results 8

Contamination detection and validation 10

BLAST cross-validation 10

BlobToolkit cross-validation 10

Comparison with BUSCO on UniProt Reference Proteomes 11

Case studies 12

Analysis of avian proteomes 14

Comparison of proteomes from closely related species 17

Human and Hominidae proteomes 17

Mouse and Myomorpha 18

Gallus gallus and Galloanserea 19

Xenopus and Amphibia 20

Zebrafish and Otophysi 21

Drosophila melanogaster and melanogaster subdivision 22

Caenorhabditis elegans and the caenorhabditis genus 23

Saccharomyces cerevisiae and saccharomycetacea 24

Arabidopsis thaliana and Brassicaceae 25

Assembly and annotation comparison 26

Assembly comparisons 26

Annotation comparisons 39

**Supplementary references** 47

### Supplementary Tables

The Supplementary Tables are available at Zenodo (doi:10.5281/zenodo.7359861)

**Supplementary Table 1**. **Source proteomes used for simulations**. List of the proteomes used as source proteome in the simulation, divided by dataset. The Model dataset corresponds to common model organisms. The Representative dataset corresponds to species from across the eukaryotic taxonomy, with two species per each major lineage.

**Supplementary Table 2**. **Contamination detection by OMArk on simulated contaminated proteomes**. The main sheet, "Summary," indicates the percentage of proteomes in which contamination was detected for each simulated dataset, by contaminant species. Each other sheet, one for each degree of contamination, details for each simulated proteome what contaminant was detected by OMArk, and recapitulates the proportion of the dataset where the detection is accurate - as in the summary sheet.

**Supplementary Table 3. Global OMArk and BUSCO results on the UniProt Reference Proteome data**. Each row represents a proteome, its OMArk statistics, and BUSCO scores. All of OMArk and BUSCO statistics are given in percentage.

**Supplementary Table 4**. **Detected contamination events and validation**. List of all contaminations detected in the UniProt Reference proteomes: the proteome and species in which it was detected, the contamination source clade, and the corresponding number of unambiguously associated proteins. Also included are the results of cross-validation of detected contaminants with BLAST and BlobToolKit.

**Supplementary Table 5**. **OMArk results for Ensembl Metazoa proteomes with recent changes in assembly or annotation**. Each row represents a proteome, its version number and its OMArk statistics. All of OMArk statistics are given in percentage.

### Supplementary Results

#### Simulation results

In order to evaluate the ability of OMArk to provide accurate quality assessment, we simulated cases of genome incompleteness, presence of erroneous sequences, and cross-species contamination on two datasets of eukaryotic proteomes. The first dataset is composed of 9 model species' proteomes expected to be of high quality due to extensive curation (**model dataset**). The second dataset is composed of 16 proteomes representing the diversity of Eukaryotes with no presence in the original OMA database, which mirrors the expected use case (**representative dataset**).

##### Incompleteness simulation

For our sets of 25 complete proteomes, we removed proteins at random from each proteome to simulate from 10% to 90% completeness, by increments of 10%. Complete results for all simulated proteomes are shown in Supplementary Figure 1 (model dataset) and Supplementary Figure 2 (representative dataset). Given that source proteomes, especially from the representative dataset, do not all have 100% completeness to start with, we consider the expected completeness as the measured completeness of the source proteome multiplied by the simulated completeness. For instance, a source proteome that is 90% complete would have an expected completeness of 9% for the 10% simulation.

For the model dataset, both OMArk and BUSCO (Manni et al. 2021) were able to accurately assess completeness, although with a slight overestimation (+2.6% on average and +2.1% on average, respectively; Supplementary Figure 1). Generally, OMArk reports more duplicated genes than BUSCO. This is expected, as OMArk uses a set of conserved genes expected to be in single-copy in the ancestral repertoire, but not necessarily in single-copy in most species, contrary to BUSCO. There were a few differences between methods, depending on the species. For example, BUSCO gives, on average, a close estimate of completeness for *Caenorhabditis elegans* (+0.5%), while OMArk overestimates it by a higher margin (+3.48%). The opposite relationship was observed for *Danio rerio* (+2.8% for OMArk, +4.6% for BUSCO).

For the representative dataset, the overall accuracy was worse for both methods (overestimation of +8.8% for OMArk, +6.1% for BUSCO; Supplementary Figure 2), although BUSCO overestimated completeness by a smaller margin than OMArk. For all but 4 proteomes, even though slightly overestimated, the detected completeness scales in a linear manner with the actual completeness. However, for *Fistulifera solaris*, *Panicum miliaceum*, *Stentor coeruleus* and *Hibiscus syriacus,* the detected completeness is always noticeably overestimated and grows logarithmically with the actual completeness. This pattern may be explained by the high degree of duplication in these species due to polyploidy (Maeda et al. 2021; Hunt et al. 2014; Slabodnick et al. 2017; Kim et al. 2017). Thus, two or more genes would need to be lost for an ortholog to be considered as missing. Although OMArk is especially sensitive to this effect, BUSCO is also affected to some extent. These results point to a limitation of using "core gene sets"-based methods like OMArk and BUSCO for completeness evaluation in polyploid species. When excluding these outlier species, the average overestimate is +5% for OMArk and +3.14% for BUSCO for the representative dataset.

Overall, OMArk can reliably evaluate proteome completeness on simulated datasets, in most cases providing a realistic estimate. For these simulations, the estimates are close to, but slightly worse than, the state-of-the-art BUSCO. In particular, OMArk is more sensitive to overestimation in the presence of many duplications in the source genome.

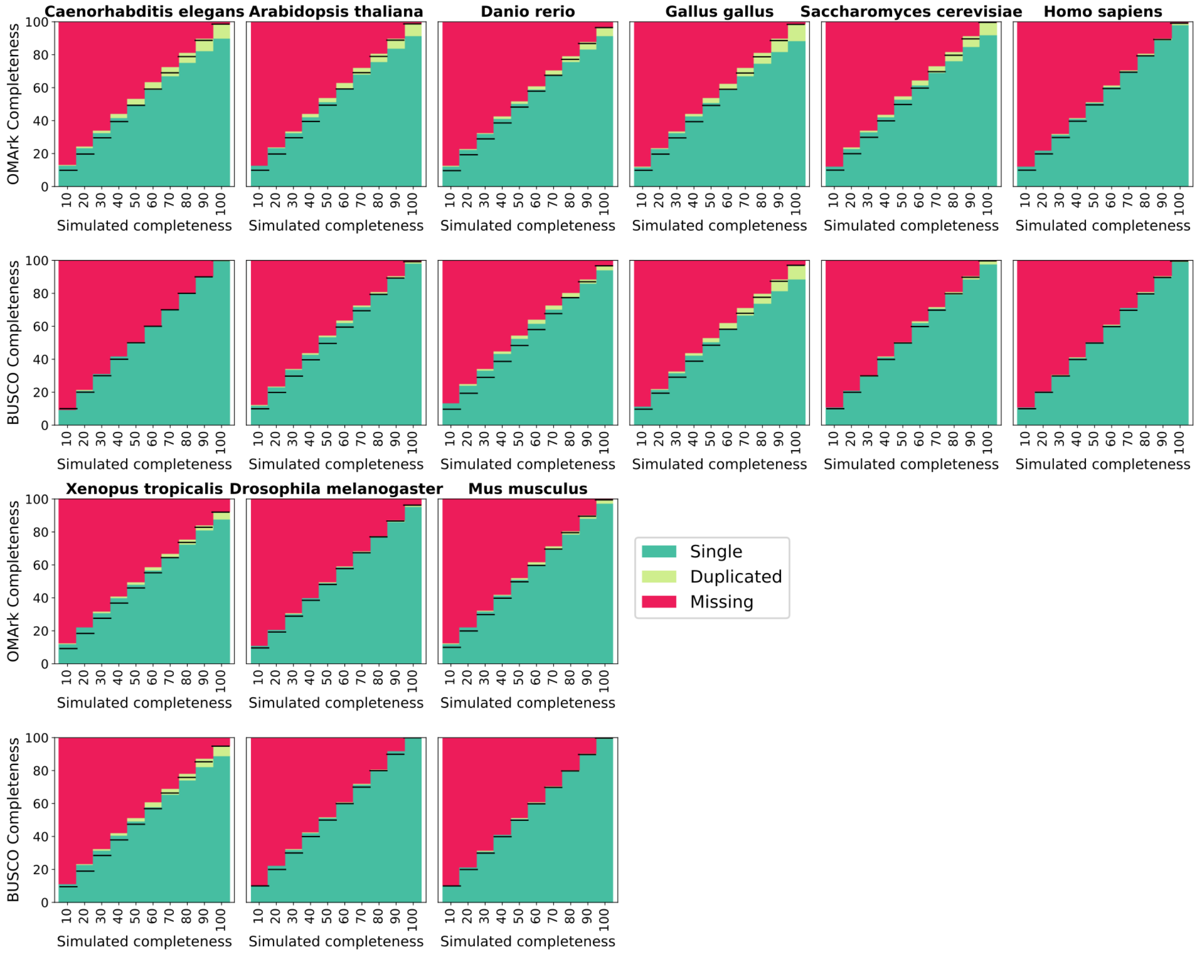


**Supplementary Figure 1**. **OMArk and BUSCO results for incompleteness simulations on the "model dataset" proteomes**. Pairs of vertical squares show the completeness statistics (y-axis) for OMArk (top) and BUSCO (bottom). Gene families (HOGs) present in single-copy (dark green), duplicated (light green), and missing (red) are shown for different levels of simulated completeness (x-axis). The dark horizontal lines show the expected completeness measurement (divide between the green and red section), considering the estimated completeness of the source proteome. The species name corresponding to the source proteome is indicated on top of each pair of squares. The results from running BUSCO and OMArk on the model dataset show that they accurately measure completeness, but tend to slightly overestimate it.

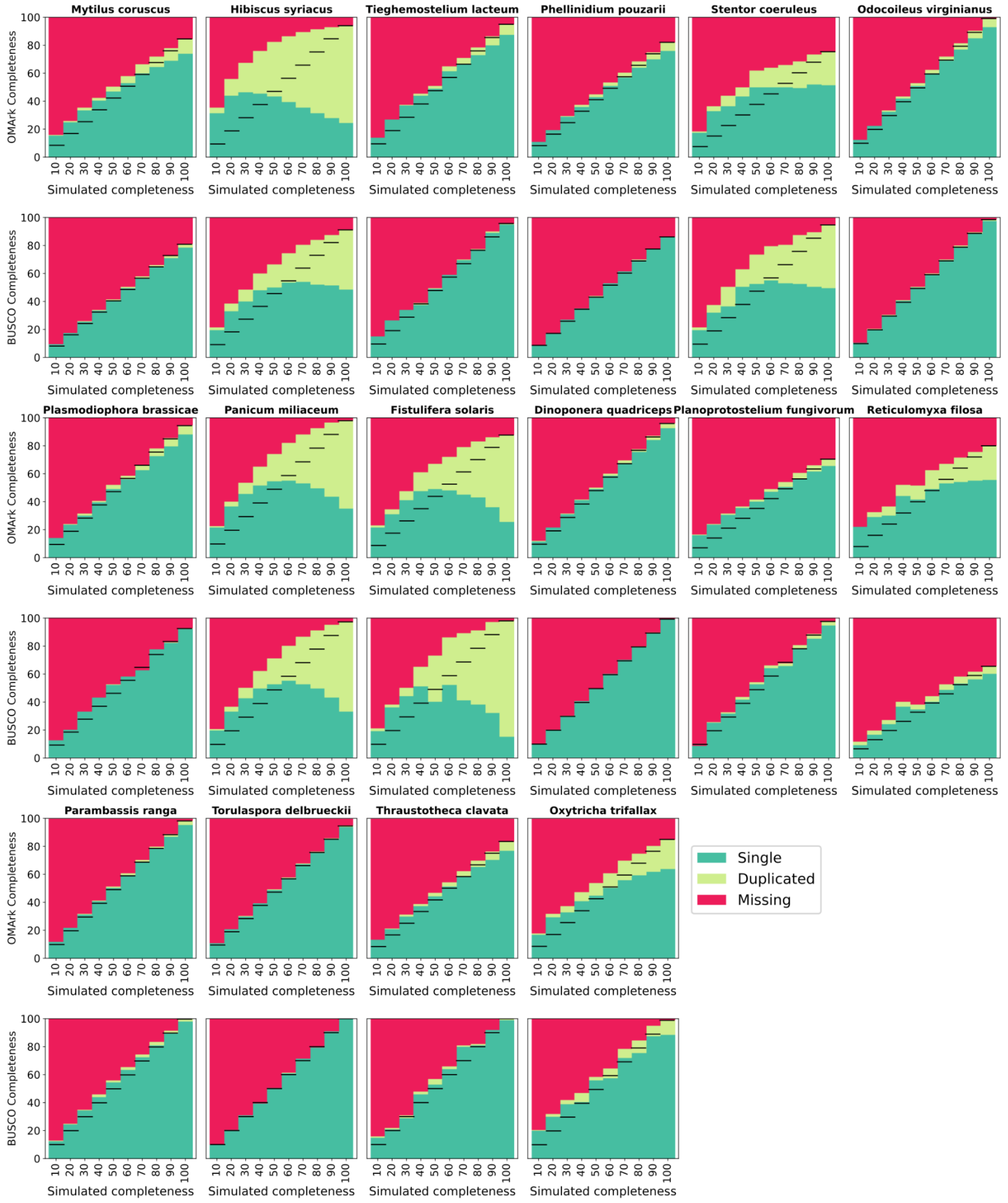


**Supplementary Figure 2. OMArk and BUSCO results for incompleteness simulation on the "representative dataset" proteomes**. Pairs of vertical plots show the completeness statistics (y-axis) for OMArk (top) and BUSCO (bottom). Conserved gene families (HOGs) present in single-copy (dark green), duplicated (light green), and missing (red) are shown for different levels of simulated completeness (x-axis). The dark horizontal lines show the expected completeness measurement (divide between the green and red section), considering the estimated completeness of the source proteome. The species name corresponding to the source proteome is indicated on top of each pair. BUSCO and OMArk accurately report completeness level in most cases, but overestimate it greatly in case of highly duplicated proteomes (polyploids).

#### Erroneous sequences simulation

For each proteome, we added randomly generated sequences to simulate proteins translated from randomly occurring open reading frames in the genomic sequence. The number of random sequences was proportional to the number of proteins in the source genome, from 10% to 90% by increments of 10%. All results are shown in Supplementary Figures 3 (model dataset) and 4 (representative dataset).

The proportion of Taxonomically Inconsistent proteins increases noticeably with the proportion of appended erroneous protein sequences (on average, +35.15% of Inconsistent genes in the 90% appended error simulation in the model dataset, +31,47% in the reference dataset). For both Taxonomically Consistently or Inconsistent, the proportion of Partial mapping and Fragments increase with the proportion of appended proteins (on average, +46.25% of total Fragment and/or Partial mapping in the 90% appended error simulation in the model dataset, +48.01% in the reference dataset). On the other hand, the proportion of proteins in the Unknown category decreases when adding random sequences. This is most common in the representative dataset where the proportion of Unknown proteins in the source proteome are high, as seen for *Mytilus coruscus* and *Reticulomyxa filosa* (on average, -0.11% of Unknown in the 90% appended error simulation in the model dataset, -3.95% in the reference dataset). Overall, this indicates that most of the randomly generated proteins are placed in gene families, but both taxonomical and structural consistency measures set them apart from real proteins. It is not clear why random sequences, which we expect to share characteristics with erroneously annotated Open Reading Frames (ORFs), are placed by OMAmer. This may be due to their overall small size (40.3 amino acids long on average), which means most of the *k-mers* in the sequence will be found in other gene families by chance.

As a result, we hypothesize that most of the protein families with no known homologs (Unknown) are longer proteins with an unusual *k-mer* content, more likely to be from lowly sampled but real gene families by virtue of their length— a long uninterrupted ORF being unlikely to occur by chance. This is reinforced by the fact that the species exhibiting the highest proportion of unmapped genes come from the lowest sampled clades in OMA, whereas mammalian species have few of these.

The number of taxonomically and structurally consistent proteins (non-hashed blue in Supplementary Figure 3 and 4) stays the same, regardless of the proportion of appended error (expected level in all simulation is shown as a red bar). Importantly, this effect is observed in both model and representative species.

As shown in this simulation, the combination of simple metrics provided by OMArk regarding the protein-coding gene set allows one to discriminate noise from high confidence protein-coding genes.

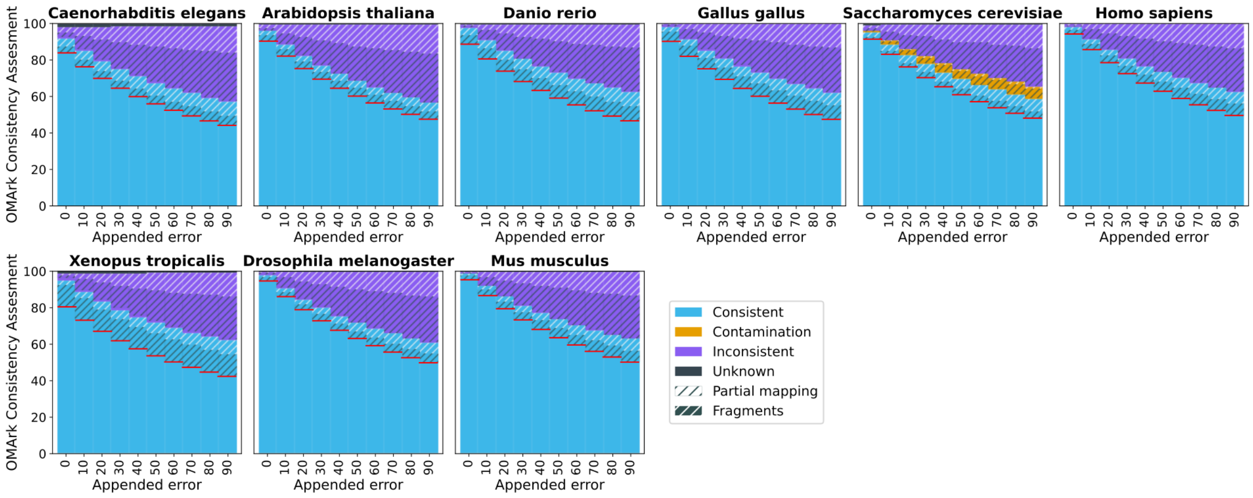


**Supplementary Figure 3**. **OMArk results for erroneous sequence simulation on "model" proteomes**. Squares represent OMArk statistics for consistency assessment using different proportions of randomly generated proteins in the source proteome, as a proportion of its original size. The species name of the source proteome is indicated on top of each square pair. The red lines show the expected proportion of taxonomically and structurally consistent genes in the source proteomes of the dataset. The proportion of taxonomically and structurally consistent genes (non-hashed blue) stays identical, regardless of the amount of noise added.

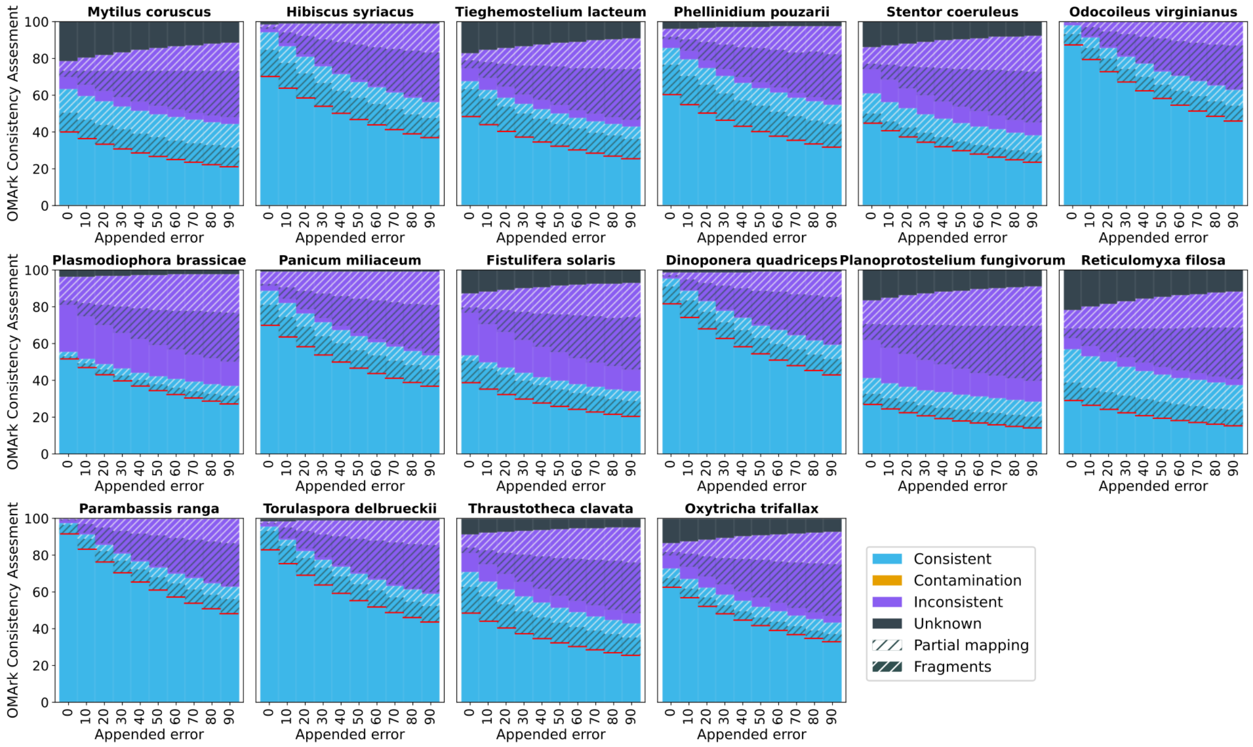


**Supplementary Figure 4**. **OMArk results for erroneous sequence simulation on "representative" proteomes**. Squares represent OMArk statistics for consistency assessment using different proportions of randomly generated proteins in the source proteome, as a proportion of its original size. The species name of the source proteome is indicated on top of each square pair. The red line shows the expected proportion of taxonomically and structurally consistent genes in the source proteomes of the dataset. The proportion of taxonomically and structurally consistent genes (non-hashed blue) stays identical, regardless of the amount of noise added.

#### Contamination simulation

We simulated different degrees of contamination with exogenous sequences by introducing, independently, proteins from a selection of species in each proteome of the model and representative datasets. Choice of the contaminant species was based on their perceived likeliness to contaminate sequence data: human, yeast, a diversity of bacteria often detected as contaminants in sequencing data, and unicellular eukaryotes were added as contaminants. We introduced an increasing number of proteins: 10, 20, 50, 100, 200, 500 and 1000. We then assessed whether the correct species or a taxonomically adjacent species was detected as the contaminant (Supplementary Table 3).

The effectiveness of contamination detection, expectedly, increases with the number of introduced proteins. Traces of contamination are detected in at least one of the simulations starting from 10 introduced proteins when the contaminant is a fungi or a bacteria, and at 50 for all but 2 species (see below). Detection of contamination appeared to be mainly dependent on the contaminant species and its taxonomic origin. Contamination was reliably detected (>50% of the samples) at 20 contaminant proteins from bacteria, 50 from fungi, and at a variable proportion for all other species, generally between 50 and 200 contaminant proteins.

With our method, contamination is harder to detect between relatively close species, for example, contamination between fungi *Torulaspora delbrueckii* and *Saccharomyces cerevisiae*, even at the highest degree of contamination. Similarly, contamination from *Homo sapiens* in other Vertebrates could not be detected unless the number of introduced proteins was high (1000 proteins).

The importance of clade sampling for accurate contamination detection is well illustrated by two Eukaryotic species from lowly sampled lineages in OMA: the amoeba *Planoprotstelium fungivorum* and the alveolata *Stentor coeruleus*. Contamination from either of the species could not be detected in any species below 1000 contaminant proteins. This is likely because the low sampling of the clade in the OMA database does not provide many specific placements.

Conversely, when the same species were used as the source genome, strong contamination often led OMArk to mistake the contaminant as the most likely main lineage for the proteome.
Nevertheless, the results we obtain show that, for contaminant species from well sampled clades and especially if the contaminant species is distant from the contaminated one, our software is able to detect even small traces of contamination in the dataset.

### Results on UniProt Reference Proteomes

We demonstrate the utility of OMArk on real data by quantifying the quality of 1805 publicly available UniProt Reference Proteomes (UniProt Consortium 2021) for eukaryotic species.

#### Global results

Figure 2 of the main manuscript (reproduced below) summarizes the results of quality assessment over the whole dataset. At first glance, it appears the overall completeness of the Reference Proteome data is high, with 1472 (81.5%) proteomes being detected with more than 80% of completeness and 64 proteomes (3.6%) missing more than 50% conserved genes. The results of the consistency assessment are more contrasted. Only 1246 proteomes (69.0%) have at least 80% of its proteins categorized as taxonomically consistent, and 459 proteomes (25.4%)

are more than 80% taxonomically *and* structurally consistent (excluding Partial mapping and Fragment). Surprisingly, 339 proteomes (18.8%) have less than half their genes that are consistent both taxonomically and structurally (neither *Partial mapping* or *Fragment*) consistent.

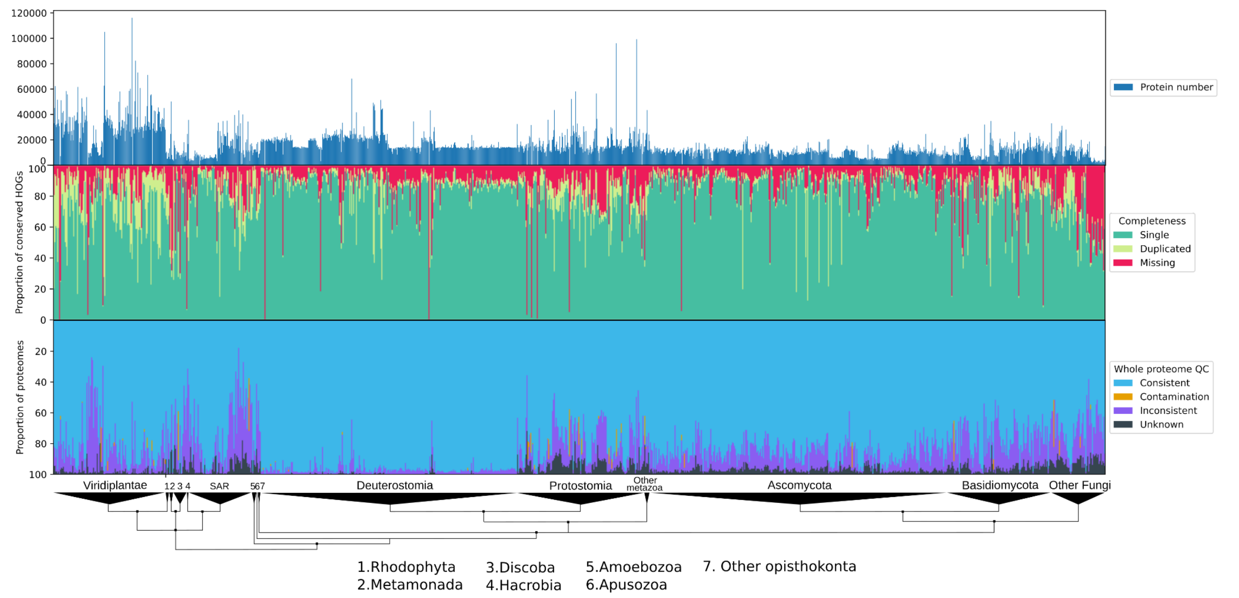


**Figure 2 (from main text). OMArk results on 1805 Eukaryotic proteomes from UniProt.** Middle - Completeness assessment: Proportion of conserved genes found (green) with a breakdown between single copy (light green) and duplicated (dark green) and missing conserved HOGs (red), in all proteomes ranked by proportion of found genes. Bottom - Whole proteome characterization: Proportion of accurately mapped proteins (blue), incorrectly placed proteins (purple), contaminant proteins (orange) and proteins with no mapping (black). Genomes are ranked by taxonomy with major eukaryotic taxa shown on a taxonomic tree at the bottom.

In terms of taxonomic specificities, a few particularities stand out. Deuterostomes have the highest proportion of taxonomically Consistent genes, however, they do not stand out in terms of completeness. A subset of the plants in the dataset has a high proportion of duplication, as expected in species often subject to Whole Genome Duplication. Finally, a few fungi, corresponding to the Microsporidian clade, appear to miss a high proportion of their genomes. This clade is represented by less than 4 species in OMA, preventing Microsporidia from being used as the ancestral reference lineage. The low completeness value is a reflection of the many losses known to have taken place in this particular clade (Nakjang et al. 2013).

We used the results of OMArk over these 1805 proteomes to assess the accuracy of OMArk's species identification. The taxon identified by our method corresponded exactly to the most specific taxon containing the species and represented in the OMA database for 838 (46.4%) proteomes. It is possible, especially when close species to the query are present in the underlying OMA database, that OMArk identifies as the main species a close relative or a sister clade to the accurate one. When using, for comparison, the ancestral lineage used by OMArk after species identification, the choice of lineage based on species identification was accurate for 1540 proteomes (85.3%). Finally, the majority of the remaining 265 species misidentification was either to an ancestral clade from the reference one (204; 11.3% of the total). Selecting such clades for the quality assessment step would lead to less specific but still accurate results.

Thus, in total, the automatic species identification of OMArk was able to select a satisfactory result in 96.6% of the cases.

#### Contamination detection and validation

Over the whole Reference Proteome dataset, OMArk detected contamination in 72 proteomes, sometimes from multiple species, for a total of 115 contamination events detected. The breakdown of detected contaminants indicates that most detected contamination comes from Bacterial species (97 out of 115; Supplementary Table 4).

In order to validate the contamination calls made by OMArk, we performed two supplementary analysis: one based on a BLAST search of a sample of the contaminant sequences in the non-redundant database and a second based on BlobToolkits results on the assembly corresponding to the proteome of interest.

##### BLAST cross-validation

For each contamination event detected by OMArk, we selected the first 10 sequences reported as contaminants and performed a search against the non-redundant database. Then for each result, we selected the best hit that was not from the query species itself and asserted whether the target species for this hit was closer to the query species or from the OMArk-determined contaminant species. Results for all of the BLAST searches were reported as closer from the potential contaminants for 73 events. For 19 other events, more than 80% of the searches were closer from the contaminant.

For those 24 samples with less than 80% successful hits, we manually evaluated the BLAST output to assess a possible contamination. Based on the distribution of best hits, we could confirm at least some of the sequences were likely exogenous and closest to the reported contaminant for an additional 9 OMArk-detected contamination events.

We could not confirm whether the 15 others were actual cases of contamination and labeled them as false positives, for a true positive rate of 87% overall. Two of the false positives (contamination from *Drosophila biarmipes* in *Saccharomyces cerevisiae* and contamination from *Picea glauca* in the fungi *Mytilinidion resinicola*) originate from contamination in our reference database. We will remove those contaminated proteomes in the next release of the OMA database. From the BLAST results, it appears that three others false positives (contamination from *cyanobacteria* in the algae *Nannochloropsis gaditana*, *Chloropicon primus* and *Auxenochlorella protothecoides*) are in fact due to detectioning chloroplastic genes as exogenous. This is likely due to sparse sampling of close species with an homologous chloroplast in the OMA database. FInally, a last event, contamination from *Lachnospiraceae* in the chytrid *Piromyces finnis* is likely another example of Horizontal Gene Transfer in an ancestral chytridiomycota erroneously detected as contamination. These false positives will likely be solved with higher sampling for those clades in our reference database.

##### BlobToolkit cross-validation

For each detected contamination event, we obtained the contaminated proteome's identifier for the corresponding assembly from UniProt and used this to query the BlobToolkit webserver. Results from BlobToolkit were available for 81 of our detected contamination events. Of those, 69 were also detected as contaminants in the assembly by Blobtoolkit, corresponding to a true positive rate of 85%. Of the 12 contamination events not detected as such by BlobToolkit, 9 of them were also marked as false positives in our BLAST analyses. The three others were contamination from a *Moraxellaceae* bacteria in the *Rhizophagus irregularis* proteome, from *Plasmodiophora brassicae* in the *Beta vulgaris* proteome, and from *Staphylococcus* *saprophyticus* in the *Cimex lectularius* proteome. However, in the BLAST results, most contaminant sequences detected by OMArk had a high sequence similarity with the identified contaminant, suggesting they are *bona fide* contamination.

In conclusion, from both the BLAST and BlobToolkit analyses, we could validate 100 of the contamination events detected, corresponding to contamination in 59 species. This corresponds to a true positive rate of 86% on the Reference Proteome data. Half of the false positives could be directly linked to sparse sampling or hitherto undetected contamination in our reference databases. These issues will be fixed in the future release of OMA. Overall, OMArk provides a viable indicator of potential contamination events, and we recommend exercising caution and more detailed analysis using complementary tools when using the contaminated proteomes.

#### Comparison with BUSCO on UniProt Reference Proteomes

We compared OMark completeness to BUSCO's assessment (Supplementary Figure 5). The results show a high correlation (Pearson: 0.81, p-value: 0) between the results but with divergence for a few proteomes. This is likely affected by differences in the ancestral clade selection. For example, results of both methods using the *Aves* (Supplementary Figure 6) lineage as source, are clearly collinear (Pearson: 0.98, p-value: $2e^{-78}$). Another salient example is the Microsporidian lineage, where the results are visibly different: as mentioned before, this is because OMArk does not select this clade as reference under default parameters because it is represented by less than 5 species in the OMA database. This leads to an underestimation of completeness in a lineage with high number of gene losses.

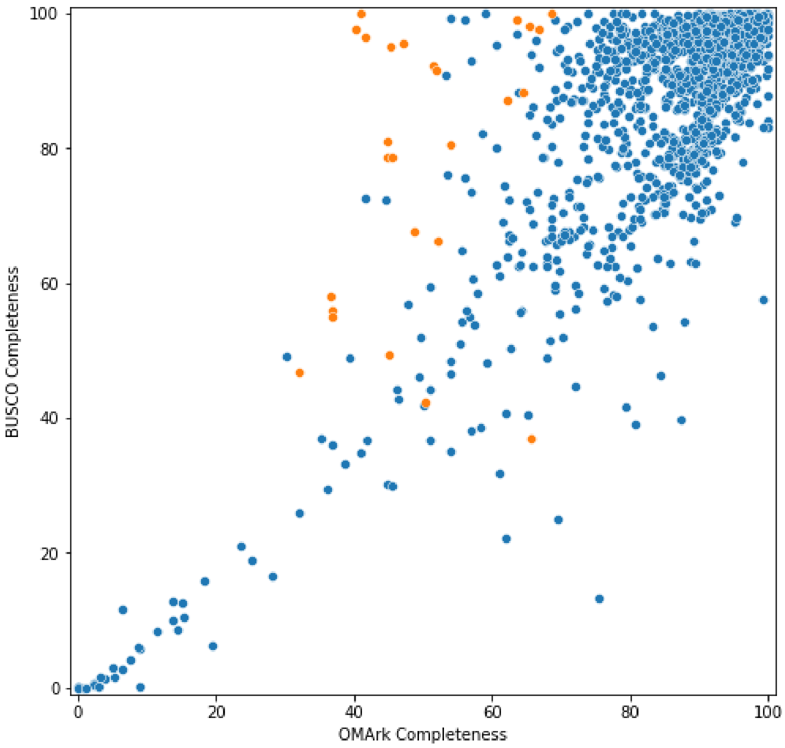


**Supplementary Figure 5**. **Comparison of BUSCO and OMArk completeness assessment of Eukaryotic proteomes**. The completeness assessment is highly correlated between methods, with some outliers. Microsporidian proteomes (orange) are an example, where OMArk underestimates completeness.

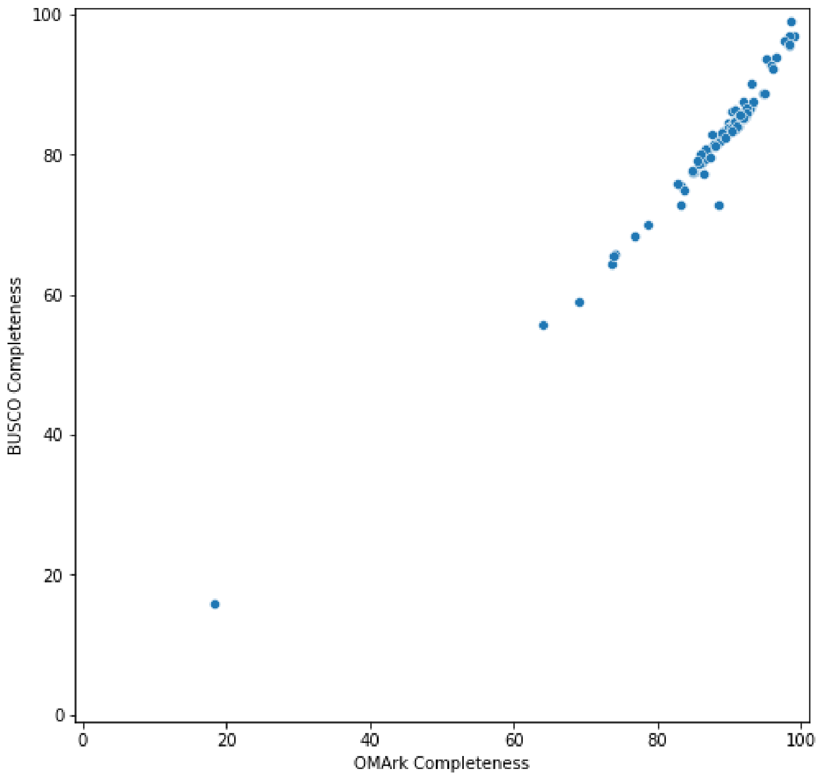


**Supplementary Figure 6. Comparison of BUSCO and OMArk completeness assessment on Aves proteomes.** The completeness assessment is nearly identical between methods.

#### Case studies

Finally, we performed targeted studies of the most extreme cases of low proteome quality according to OMArk, to confirm it was representative of *bona fide* artifacts in the source data.

The proteomes with the lowest completeness values, according to both OMArk and BUSCO, tend to be proteomes with a low protein count, with a median of 691 proteins for proteomes with <20% completeness (single + duplicated). This is expected and indicates that the completeness value is likely representative of the proteome as a whole. We focus here on a few exceptions: proteomes with low completeness statistics and high protein count such as the white spruce *Picea glauca* (6,155 proteins, OMArk completeness: 8.90%) and the Fungi *Auricularia subglabra* (strain TFB-10046 SS5) (5,290 proteins, OMArk completeness: 14.52%).

For the first example, *Picea glauca,* looking at the proteome characterization statistics gives an additional insight: 67.93% of the proteins that are placed in taxonomically inconsistent HOGs correspond to Partial mapping (36.85%) or Fragments (30.85%). Even the ones that are homologous to gene families consistent with its lineage (29.75%) have a higher proportion of divergent gene structures compared to the known members of their gene families and are classified as Partial mapping (8.92%) or Fragmented (19.22%). This indicates that the proteome for this species contains a high proportion of mis-annotated genes, and otherwise not well-defined gene models. Furthermore, the number of proteins in the proteome is uncharacteristic of land plant species and while no other Acrogymnosperm species is present in UniProt for comparison, the chinese yew *Taxus chinensis* possesses more than 44,000 coding-genes according the NCBI database. It is thus likely that this proteome is both incomplete and contains mostly erroneous sequences.

The picture is different for *Auricularia subglabra*, which, despite lacking a high proportion of the 'Conserved HOGs' (OMArk completeness: 14.52%), has 56.98% of genes that corresponds to expected gene families for its taxonomic division (*Consistent*: 40.13%). This indicates that at least half of the gene count consists of well-annotated genes. The low completeness level suggests that the complete genome is expected to have a much higher count of protein-coding genes than the 5,290 reported in the proteome. This is confirmed in the literature, where the number of protein coding genes for species in the genus is around 16,000 protein coding genes (Dai et al. 2019).

The proteomes with the highest degree of proteins placed into taxons inconsistent with the species' lineage tend to be from taxonomic groups underrepresented in the OMA database. Of the 10 proteomes with the highest "Inconsistent" score, we count 3 from the SAR division, 4 chlorophytes, 1 Rhodophyte, and 1 Apusozoa. In these cases, a high proportion of inconsistently placed genes may be due to uneven sampling of these taxonomic divisions in the OMA database and not only to dubious gene models. For example, the species with the highest Inconsistent percentage (83.93%) is *Cafeteria roenbergensis,* a species from the Stramenopiles clade. When performing a BLAST similarity search against the non-redundant NCBI database using sequences from the *Inconsistent* category, we consistently found high identity hits in multiple strains of the same *Cafeteria roenbergensis* species, with subsequent hits belonging to a variety of other species and with much lower identity, indicating a likely distant homology relationship.

High proportions of inconsistently placed proteins can also be due to faulty taxonomic labeling. For example, the *Paulinella micropora* proteome also displays a high proportion (77.15%) of Taxonomically Inconsistent proteins. From the known taxonomy of this species, OMArk used the SAR clade as the ancestral lineage. The *Paulinella micropora* proteome is detected as incomplete (only 7.4% conserved HOGs), yet most misplaced proteins are a perfect match. These misplacements appear to be due to exogenous proteins: OMArk's automatic species placement attributes the whole proteome to the *Synechococcaceae* family, a clade of cyanobacteria and, importantly, is not able to detect any other species from the protein placement. Since *Paulinella micropora* is a unicellular eukaryote known to have recently acquired (90-140 MYA) a plastid by primary endosymbiosis (Lhee et al. 2019), we can thus hypothesize that this proteome contains mainly genes derived from the genome of the plastids of this species.

In these two cases, the Inconsistent statistics are not necessarily an indicator of erroneous gene prediction, but indicate either species with no close relatives or inaccurate species labeling. In both cases, most of 'Inconsistent' genes are neither labeled as Fragmented nor Partial hits, as would be expected for erroneously predicted genes (like in our simulations). One can thus use this additional information to discriminate a situation where the Inconsistent proportion is indicative of OMArk misattributions rather than faulty gene predictions.

To exemplify this, the proteome of the chlorophyte *Haematococcus lacustris* also displays a high number of Inconsistent placement (68.85% of the proteome). In this case, however, a large fraction of it is detected as Partial mapping (11.07%) or Fragmented (43.70%), consistent with erroneous gene models. The fact that the protein count of this species (28,121) is 10,000 genes higher than any other chlorophytes in the dataset despite a likely incomplete proteome (72.01% OMArk completeness), makes the possibility of erroneous gene models more likely.

### Analysis of avian proteomes

In the precomputed UniProt Reference Proteomes, most of the 234 avian proteomes are characterized by an unusually high proportion of Fragmented genes (average Taxonomically Consistent Fragment proportion: 16.6%, standard deviation: 4.23) (Supplementary Figure 7). This proportion of fragments is not uniform across species (0.57% minimum to 29% maximum of Taxonomically Consistent Fragments). They vary depending on the source of the external annotation data used by the UniProt Reference proteome (Supplementary Figure 8). Proteomes from Ensembl (n=9, average Consistent Fragment proportion: 2.84%, standard deviation: 2.09) (Cunningham et al. 2022) and NCBI Refseq (n=4. average Taxonomically Consistent Fragment proportion: 2.41%, standard deviation: 1.84) (O'Leary et al. 2016) have much less Fragments than proteomes uploaded by the Bird 10k consortium (b10k) (Feng et al. 2020), which represent most of the dataset (n=212, average Taxonomically Consistent Fragment proportion: 17.58%, standard deviation: 2.26).

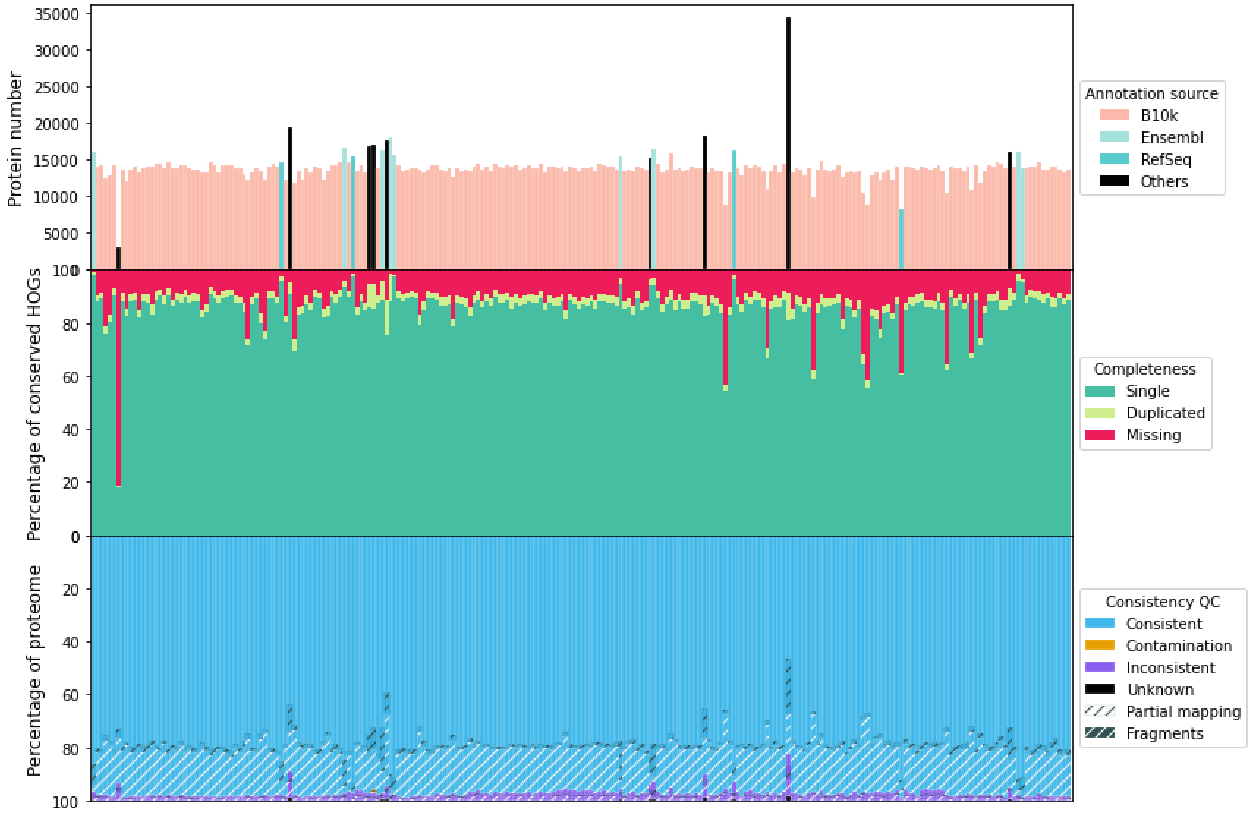


**Supplementary Fig 7. OMArk results for Avian proteomes**. Each column corresponds to an avian species in the UniProt Reference proteome, ordered taxonomically according to the NCBI taxonomy. Top subplot represents the number of protein-coding genes in each proteome, colored by annotation source. Middle subplot represents completeness, as the proportion of conserved genes present or missing. Lower subplot represents the consistency assessment. Proteomes with b10k annotation are characterized with high amounts of fragments and missing genes.

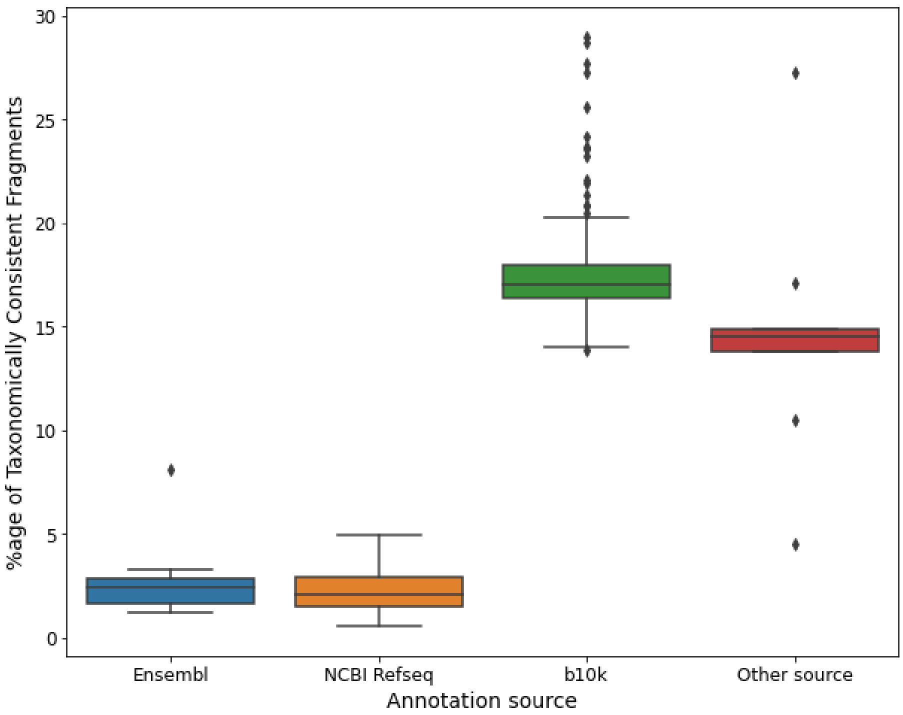


**Supplementary Fig 8. Relation between fragment proportion and annotation source**. Each category corresponds to a source of annotation. Proteomes with b10k annotation are characterized with high amounts of fragments while proteomes from Ensembl and Refseq have less of them..

The proportion of fragments being similarly high from all proteomes from the b10k consortium suggests that it is due to systematic bias in the data generation. In order to confirm this, we measured the overlap between the Fragmented gene set between each bird species by using the set of root HOGs into which these fragmented genes were placed by OMAmer. All pairwise overlap measures (calculated as the cardinality of the intersection of the sets divided by the cardinality of the smallest set) are represented in Supplementary Figure 9. The overlap between the b10k proteome fragment set is higher (average: 0.74) than with proteomes from different sources (average: 0.31). Hierarchical clustering based on overlap confirms that proteomes from the b10k annotation dataset form homogeneous clusters in terms of shared fragments.

Specifically, according to the clustering, the minimal partition of the data including all of them only include one proteome from another annotation source - *Oxylabes madagascariensis*. Notably, the annotation for this last species was made available from a previous phase of the Bird 10k project, suggesting this overlap is indeed due to systematic bias. The clustering reveals additional substratification of the dataset in terms of fragment overlap, suggesting another source-based systematic bias in part of the dataset, but we could not identify an obvious cause. Still, these results show high similarity in terms of fragmented genes for proteomes originating from this project. It could either be related to bias in the sequencing, assembly or annotation process to generate these data. Because fragmented proteins from this dataset tend to fall within the same gene families, it is most likely the latter.

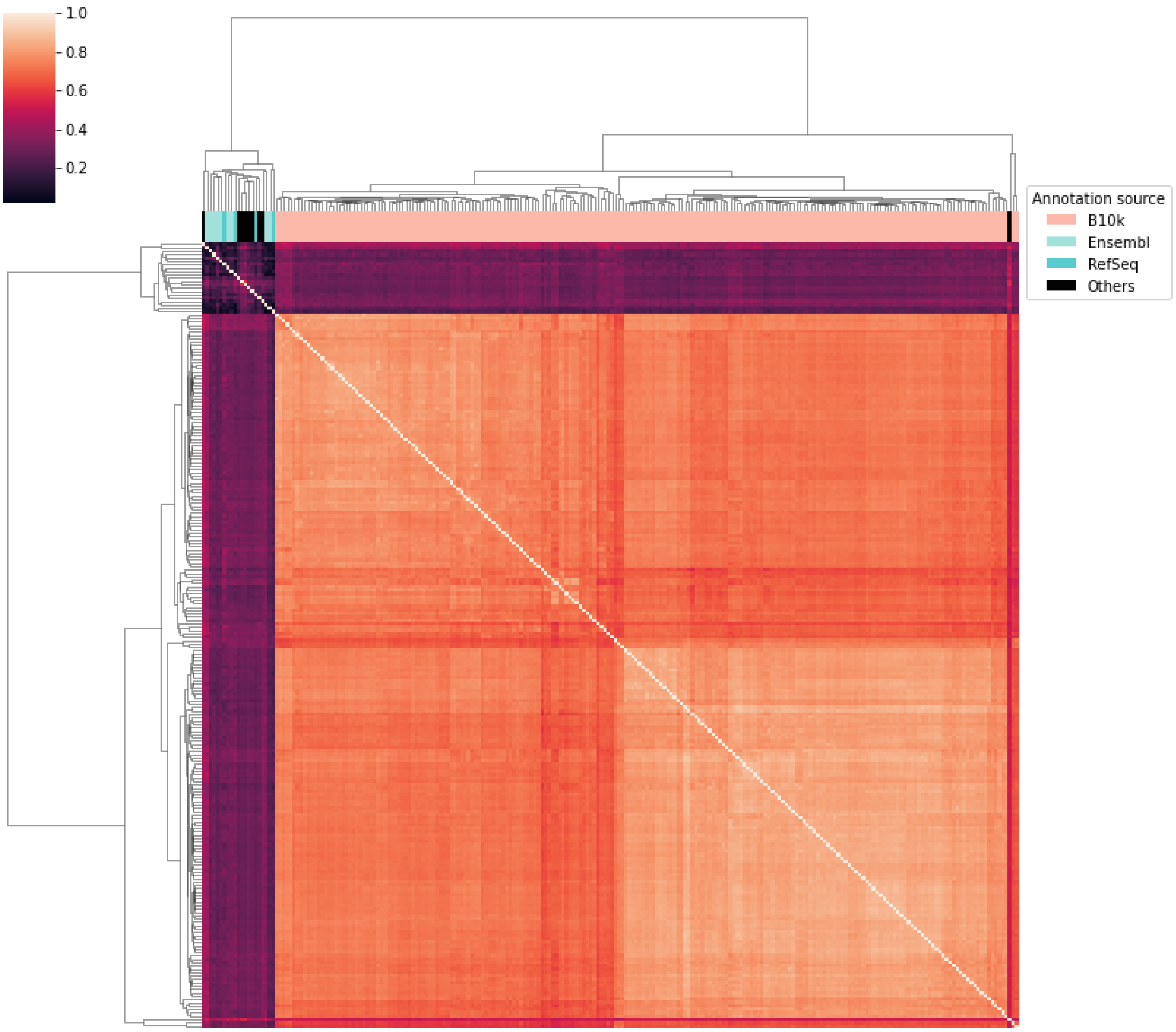


**Supplementary Figure 9. Clustered heatmap of overlap of sets of fragmented gene families between genomes.** Pairwise overlap is measured as the size of the set intersection over size of the smallest set and is represented by the color in the heatmap, as coded by the color scale. All proteomes from b10k cluster together and away from other sources of annotation, with high similarity between fragmented genes.

The annotation protocol for the b10k annotation includes, among other sources of evidence, homologous annotation from the Ensembl 85 (Cunningham et al. 2022) version of the zebra finch (*Taeniopygia guttata*) proteome. This proteome was annotated on the TaeGut3.2.4 version of the genome assembly. OMArk assessment for this version of the zebra finch proteome reports 12.47% lineage consistent fragments, much higher than for the current assembly for the same species included in our dataset (1.59%). This is consistent with the improvements of gene models in Zebra finch with the latest assembly noted in (Rhie et al. 2021). We hypothesize that part of the cause for the high proportion of fragments in the b10k annotation is due to error propagation from this fragmented proteome. Accordingly, comparing the overlap between the set of gene families with fragments from the previous version of zebra finch annotation (Supplementary Figure 10), we show that proteomes from the b10k annotation share, in average, more than half of their fragments with it (average percentage of fragments shared: 58.98%), more than double the proportion in proteomes from any other sources (average percentage of fragments shared: 26.61%).

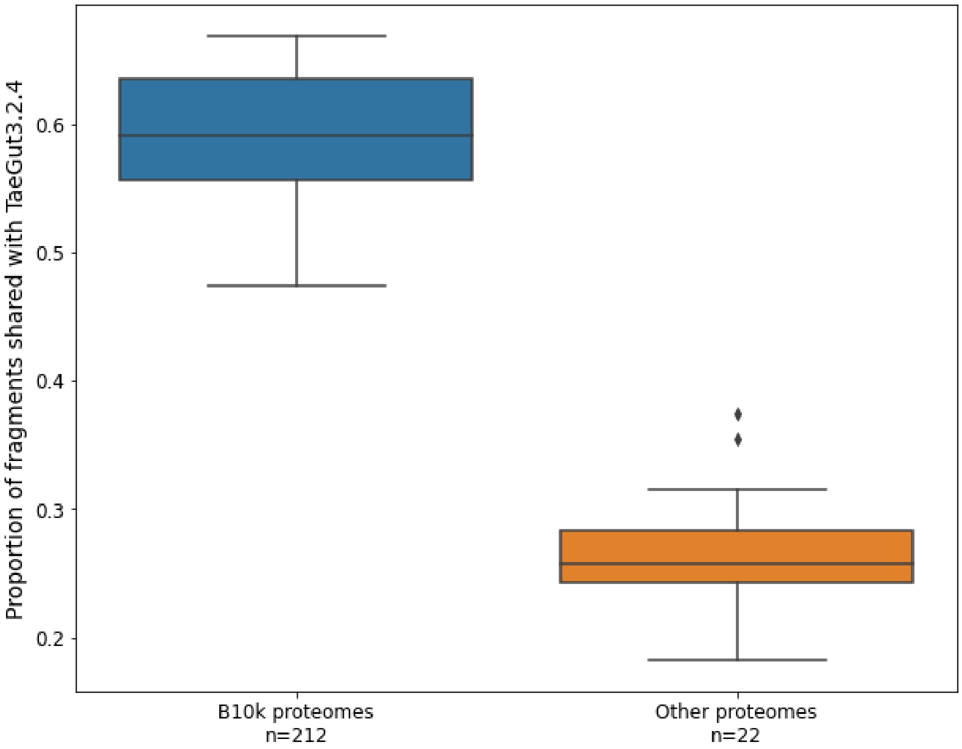


**Supplementary Figure 10. b10k annotated proteomes share many fragments with the previous version of the zebra finch proteome**. Proportion of fragmented genes that are also fragmented in the proteome for TaetGut3.2.4 assembly. All proteomes from b10k have more fragments in common with their genomes than any other proteome in the dataset.

Taken together, these results provide evidence that the high proportion of fragments in bird species in the UniProt Reference Proteome result, at least in part, from error propagation from the reference proteomes used to perform gene annotation.

### Comparison of proteomes from closely related species

Similar to BUSCO or other methods based on reference gene sets, OMArk is best used for comparing the proteomes of closely related species. In this context, it can be used to identify the best quality proteomes within a certain taxonomic division. We exemplify this by comparing the OMArk assessment of well-studied, model organisms to the closest related species in the UniProt Reference Proteome dataset.

#### Human and Hominidae proteomes

Proteomes from the Hominidae taxa are homogeneous in terms of OMArk statistics (Supplementary Figure 11), with high completeness and a high proportion of genes found to be Consistent with their taxonomic division, with few Fragments and Partial mapping. As expected, the human proteome has the lowest proportion of missing genes (0.87%), with very similar results as the chimpanzee (*Pan troglodytes*, 1.1%). The closely related *Pan paniscus* appears to be missing more genes than both (2.83%). In terms of consistency assessment, *Gorilla gorilla* and *Pongo abelii* have a higher proportion of divergent gene structure than the three others (91.68% and 90.76% of taxonomically and structurally Consistent genes, respectively, compared to 94.26%-94.52% for the rest). In addition, *Pongo abelii* also displays a higher proportion of Missing (6.54%) and Duplicated (8.60%) genes than all the other proteomes, likely indicating a slightly worse assembly or gene annotation.

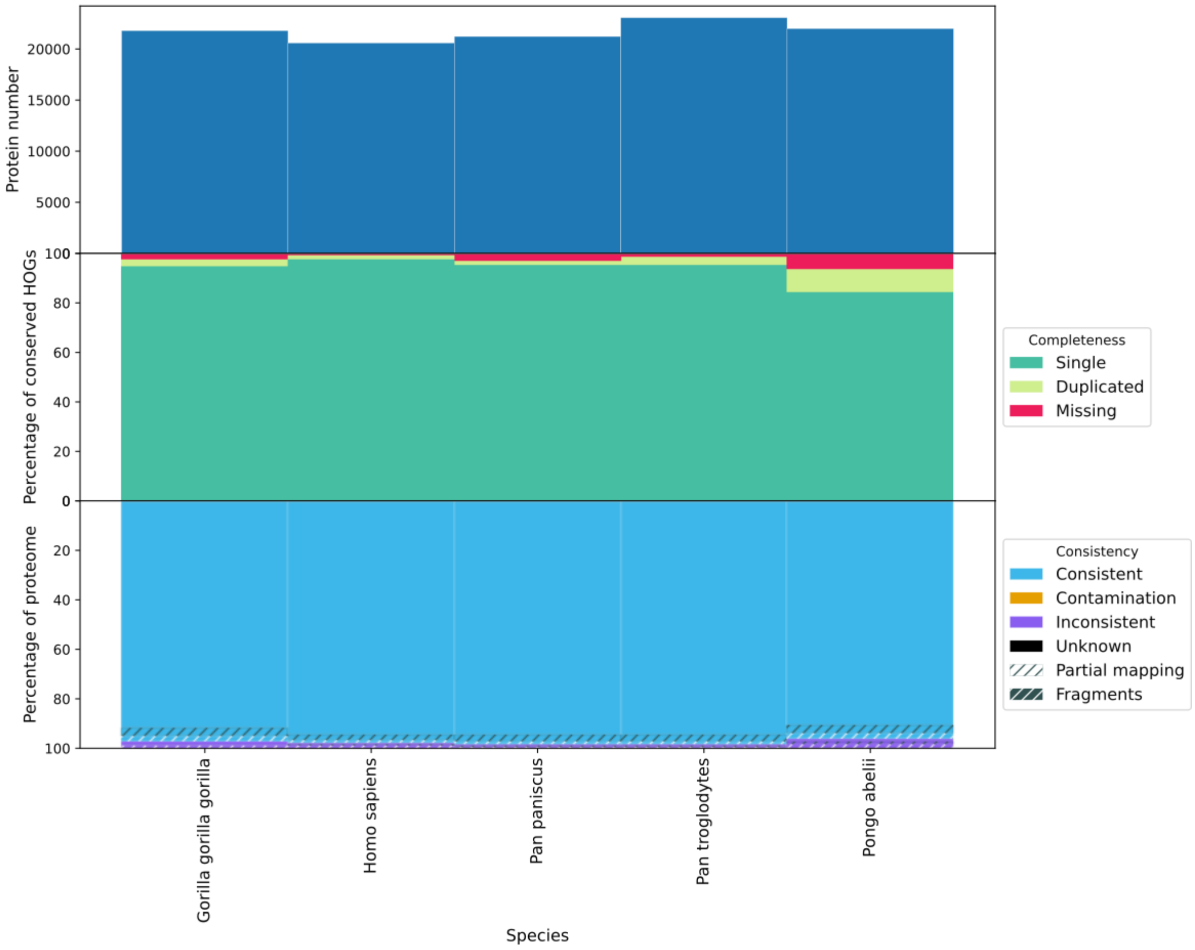


**Supplementary Figure 11**. **OMArk results for proteomes from the Hominidae clade.**

#### Mouse and Myomorpha

Proteomes from the Myomorpha clades are noticeably different, both in terms of completeness and of gene model quality (Supplementary Figure 12). As expected, the Mouse proteome is highly complete (99.47% completeness) and consistent (95.34% of taxonomically and structurally Consistent genes), as is the *Rattus norvegicus* proteome, another well studied rodent (98.15% completeness, 95.17% taxonomically and structurally Consistent genes). *Mesocricetus auratus* and *Peromyscus maniculatus* have a slightly higher proportion of genes with divergent gene models (88.33% and 89.17% taxonomically and structurally Consistent genes, respectively) and Duplicated genes (6.71% and 4.76%, respectively), indicating the proteome may be less reliable. Finally, *Neotoma lepida* and *Cricetulus griseus* display a comparatively high proportion of Missing (9.21% and 14.97%, respectively) and Duplicated genes (12.08% and 8.35%, respectively), and less than 60% of taxonomically and structurally Consistent gene models. This indicates the gene content is not only incomplete but of lesser quality overall. This is outlined by the number of reported coding genes being higher from the other species by a few thousand, while being less complete overall. This result is particularly notable given *Cricetulus* *griseus* is included in our reference database, showing our method is somewhat robust to circularity (flagging a proteome as high quality because it has an exact counterpart in the reference dataset). Accordingly, one should be cautious when using these in comparative studies.

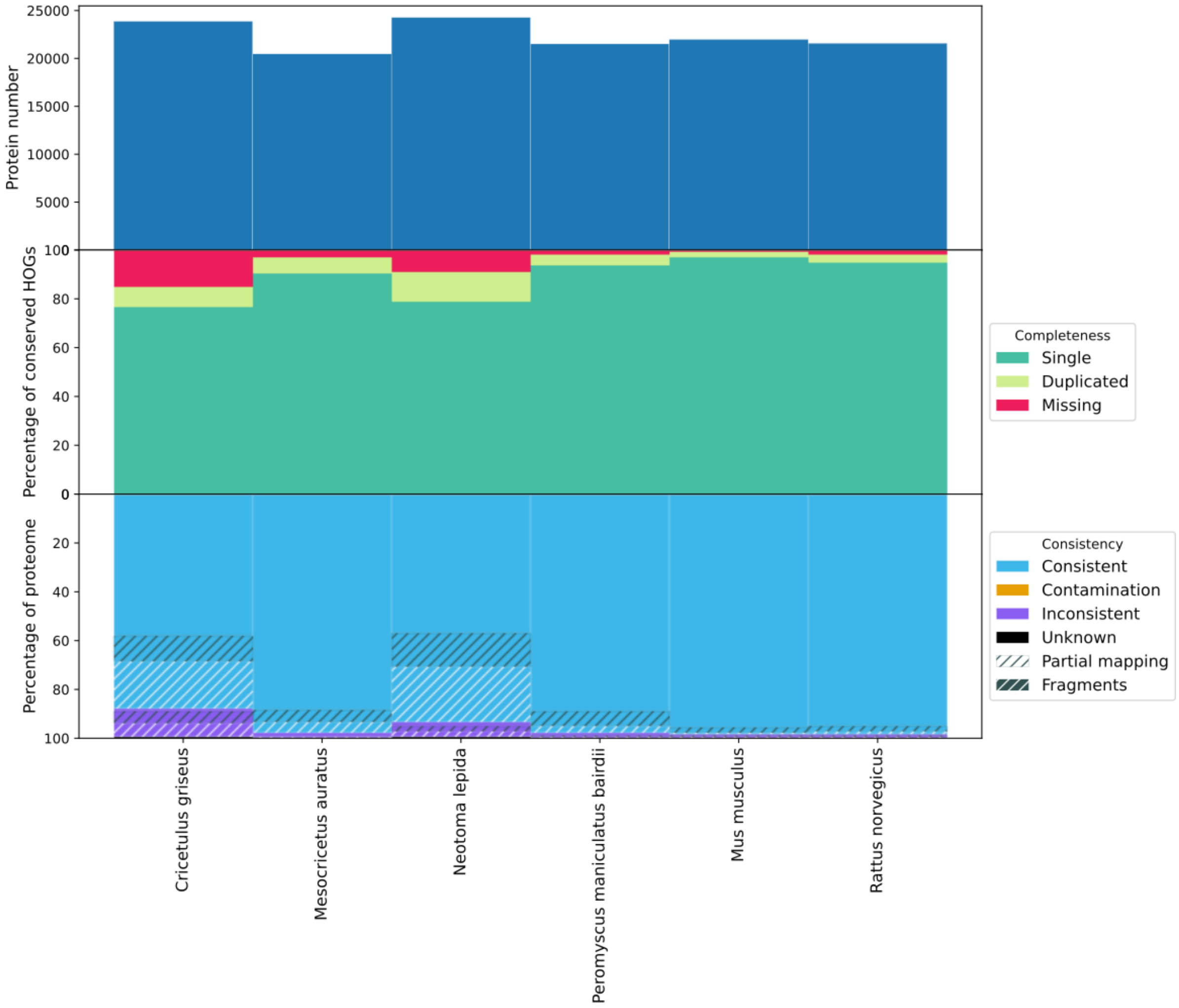


**Supplementary Figure 12**. OMArk results for proteomes from the Myomorpha clade.

#### *Gallus gallus* and Galloanserea

In the Galloanserae clade (Supplementary Figure 13), OMArk statistics are highly heterogeneous. All proteomes display some extent of Partial mapping (minimum 3.51% for *Aythya fuligula*). This is possibly due to the gene set included in the OMA database being imperfect, thus having an impact on the assessment. However, this bias is expected to have a similar impact over all proteomes. Both in terms of completeness and consistency, three proteomes from this clade stand out as more reliable: the chicken *Gallus gallus*, the pheasant *Phasianus colchicus* and the tufted duck *Aythya fuligula.* The chicken shows an uncommonly high number of Duplicated families (10.08%) compared to the two others (0.9% each), which is reflected by the number of proteins. Other proteomes in this clade are less complete and have a higher proportion of fragmented gene models. This is, in part, explained by the use of different annotation methods (See above section - Analysis of avian proteomes).

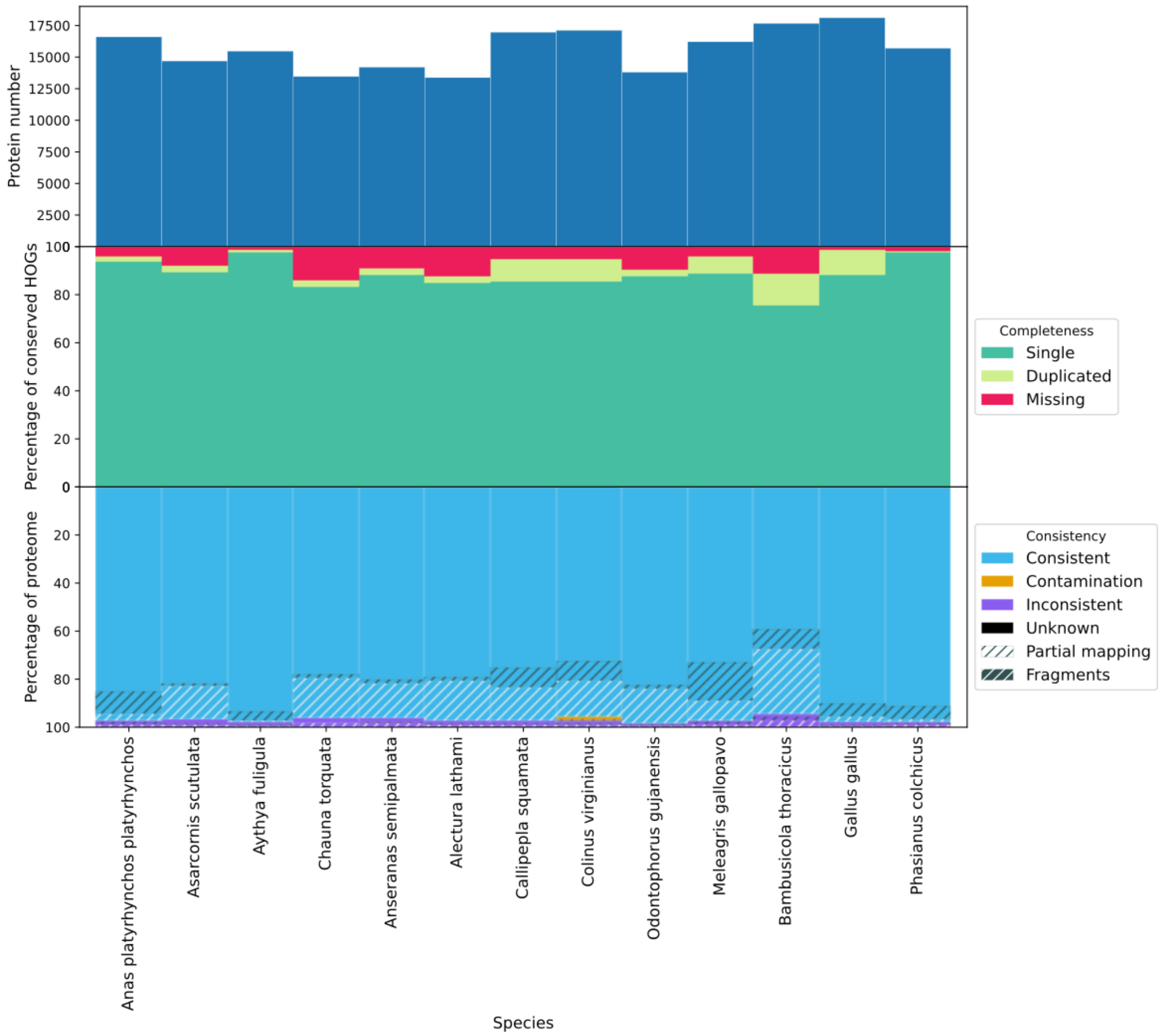


**Supplementary Figure 13. OMArk results for proteomes from the Galloanserae clade.**

#### Xenopus and Amphibia

Due to the low sampling of Amphibian genomes in the OMA database, OMArk data assessments for this clade are based on the Tetrapoda ancestral genomes (Supplementary Figure 14). It is thus expected to be less precise than other species where the ancestral lineage is more recent. The *Lithobates catesbeianus* proteome is missing most of its gene, however, likely due to an upload error of this proteome onto UniProt and it can be ignored for this analysis. In this example, *Xenopus tropicalis*, *Geotrypetes seraphini* and *Microcaecilia unicolor* have a close amount of taxonomically Consistent gene placements (94-96%) and with consistent gene structure (80-82%), and all have high completeness. However, *Xenopus tropicalis* is missing nearly twice as many genes as the others (7.96% versus 4.34% and 5.06%). This is unexpected as it is considered a model organism for this taxonomic range. Finally, *Xenopus laevis* has a high number of duplicated genes, as expected for a polyploid species, however the proportion of inconsistent genes and structurally divergent ones is also higher than all other representatives of the clade, suggesting issues with the annotation.

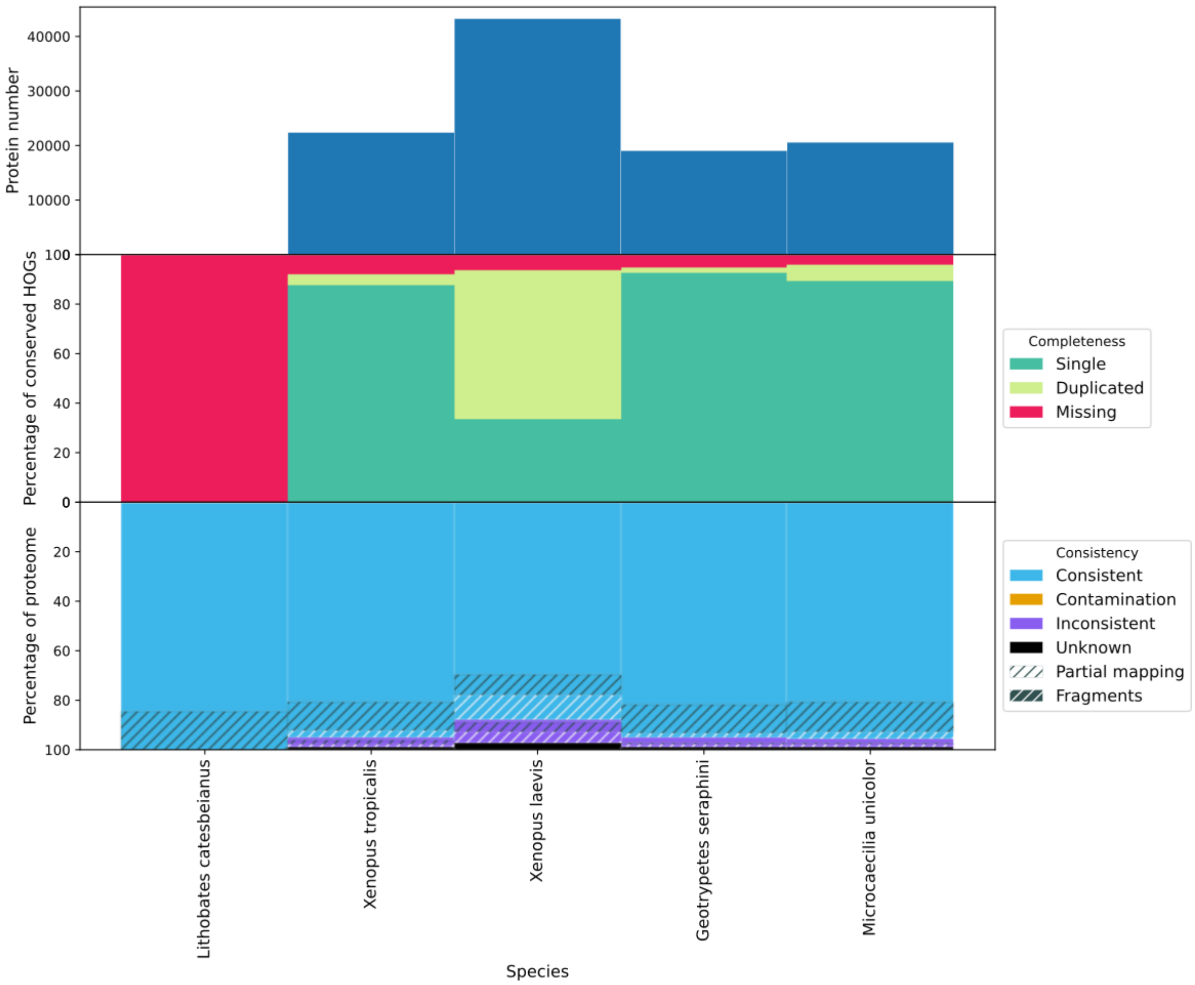


**Supplementary Figure 14**. **OMArk results for proteomes from the Amphibia clade.**

#### Zebrafish and Otophysi

OMArk results in Otophysi (Supplementary Figure 15) are heterogeneous, with all proteomes including the best annotated ones, having around 8% of genes detected as having a divergent gene structure. As all proteomes appear to be impacted, this may be due to a limitation in the reference gene set. Still, *Danio rerio* as well as *Electrophorus electricus* and *Sinocyclocheilus grahami* are detected as the most consistent to known gene families in terms of overall gene model accuracy (between 88% and 90% taxonomically and structurally consistent genes) while all having less than 5% detected missing genes. Species from the *Sinocyclocheilus* genus and *Carassius auraeus* all have the lowest proportion of missing genes and a high proportion of duplicated genes, likely due to a recent Whole Genome Duplication in the ancestor of these species (Xu et al. 2019). Finally, we can note that the Mexican tetra *Astyanax mexicanus* proteome is highly complete but appears to have issues in terms of gene models when compared to other species from the same clade (12.15% Partial mapping and 4.83% Fragmented consistent genes).

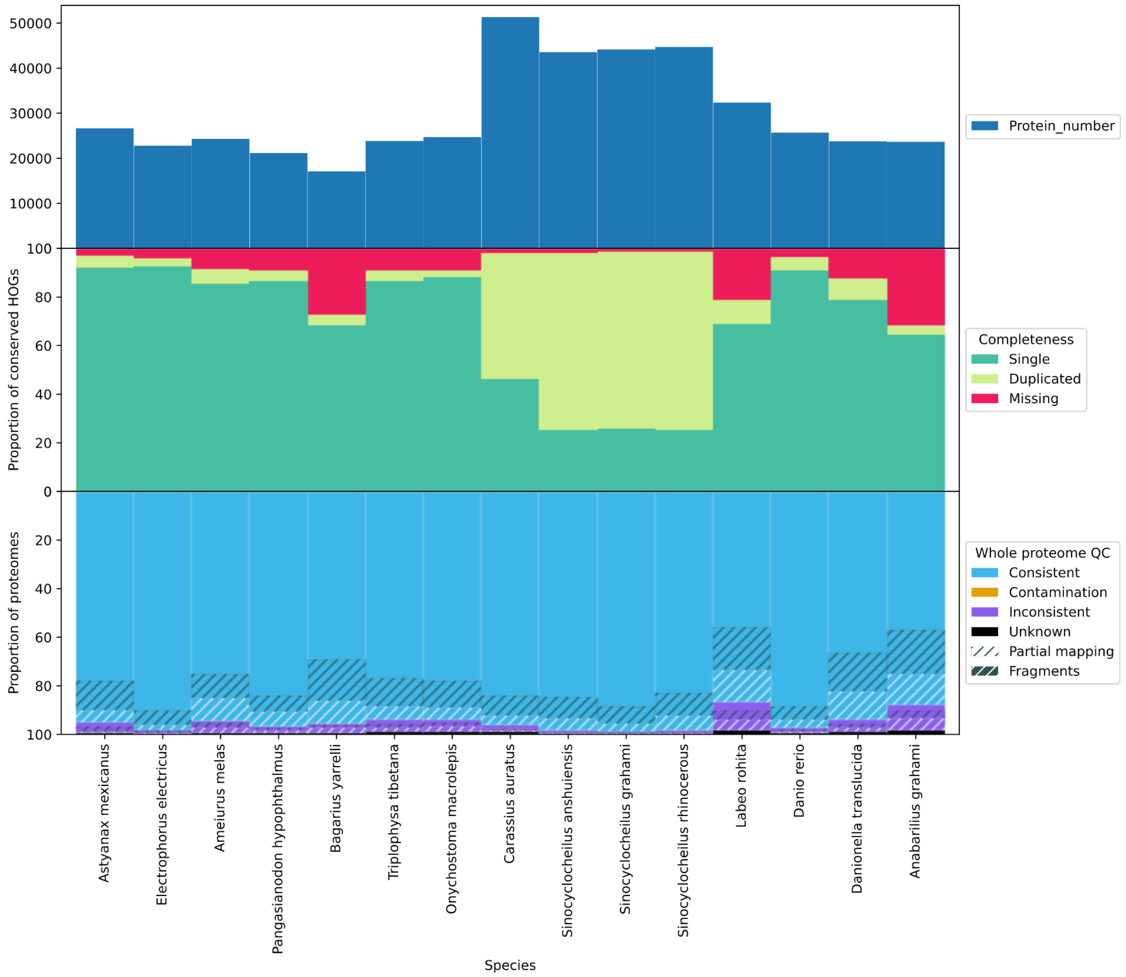


**Supplementary Figure 15. OMArk results for proteomes from the Otophysi clade.**

#### *Drosophila melanogaster* and melanogaster subdivision

Due to a high sampling in the *Drosophila* genus in the OMA database, the fruit fly *Drosophila melanogaster* is compared to the ancestral gene set of the *melanogaster* subdivision (Supplementary Figure 16). In terms of taxonomically and structurally consistent proteins, *D. melanogaster* obtains distinctly higher measures than the other two species (94.70% non-partial, non-fragment *C*onsistent genes against 83.84% and 81.32% for *D. sechellia* and *D. simulans,* respectively), despite all three species being present in our reference dataset. In comparison with *D. sechellia, D. melanogaster* has a higher proportion of Missing conserved genes (3.75% and 1.63%, respectively), which may be due to a higher number of gene losses as a domesticated species rather than incompleteness. Accordingly, when selecting the Diptera clade as a reference gene set for the same proteome, the proportion of core genes missing drops to 0.13%.

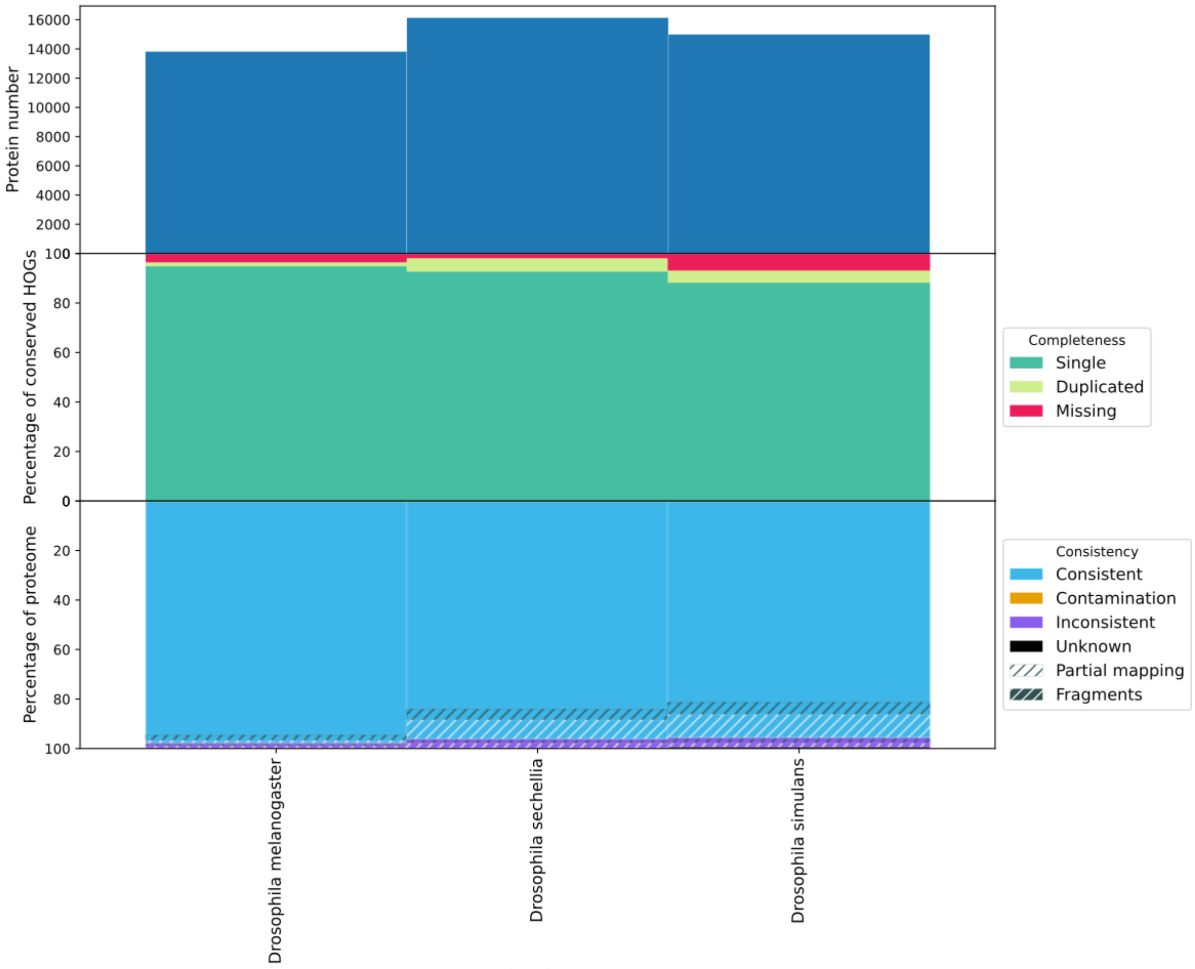


**Supplementary Figure 16. OMArk results for proteomes from the “melanogaster subdivision” clade.**

#### *Caenorhabditis elegans* and the *caenorhabditis* genus

The *caenorhabditis* genus (Supplementary Figure 17) is another example of high heterogeneity in terms of OMArk results but also in term proteome content, with a variation of gene number from 19,812 for *Caenorhabditis elegans* to 28,999 in *Caenorhabditis brenneri*. *Caenorhabditis elegans*, the most studied and curated species in this dataset, has both the lowest number of missing genes (1.43%) and the highest proportion of taxonomically consistent genes (91.78%). This can not be explained by being present in our reference database since *briggsae*, *brenneri* and *japonica* are as well but are marked as less consistent overall. Interestingly, the two other most complete proteomes, *Caenorhabditis briggsae* and *Caenorhabditis nigoni,* have a similar amount of *M*issing genes (2.63% and 2.70%), however *briggsae* appear to have more taxonomically and structurally *C*onsistent genes (75.66% Taxonomically and Structurally Consistent compared to 60.47% for *C. nigoni*), indicating this proteome might be more viable overall than the other non-*C. elegans* species.

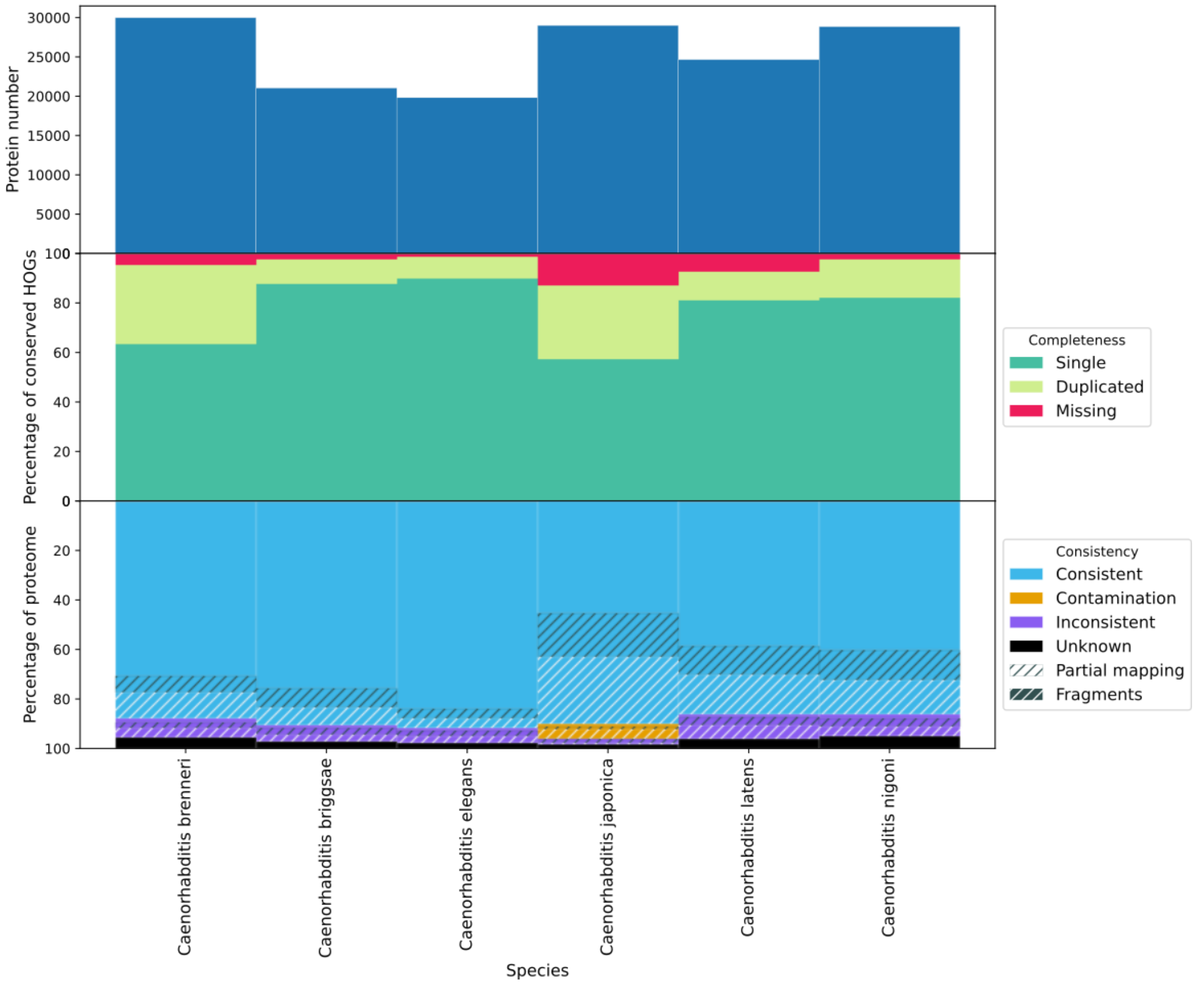


**Supplementary Figure 17. OMArk results for proteomes from the Caenorhabditis clade.**

#### *Saccharomyces cerevisiae* and saccharomycetacea

Many species in the *Saccharomycetaceae* family (Supplementary Figure 18) exhibit high completeness (less than 1% missing genes) and high consistency (95%) in most cases. However, some species, disseminated across the taxonomy, have around 5% of missing genes and 10% partial mapping genes. It is unclear if this is an annotation quality issue or a result of gene repertoire plasticity in fungi species, since these proteomes are the ones not represented in the OMA database. The two other representatives of the *Saccharomyces* genus, *Saccharomyces arboricola* and *Saccharomyces kudriavzevii* can be marked as incomplete in relation to close species, with 26.28% and 28.91% of missing conserved genes, consistent with their low number of coding genes.

In *Saccharomyces cerevisiae* s288c, OMArk detects contamination from *Drosophila biarmipes.* This is unexpected in a model species and is actually an artifact resulting from contamination in the proteome of *D. biarmipes* by yeast sequences in our reference database.

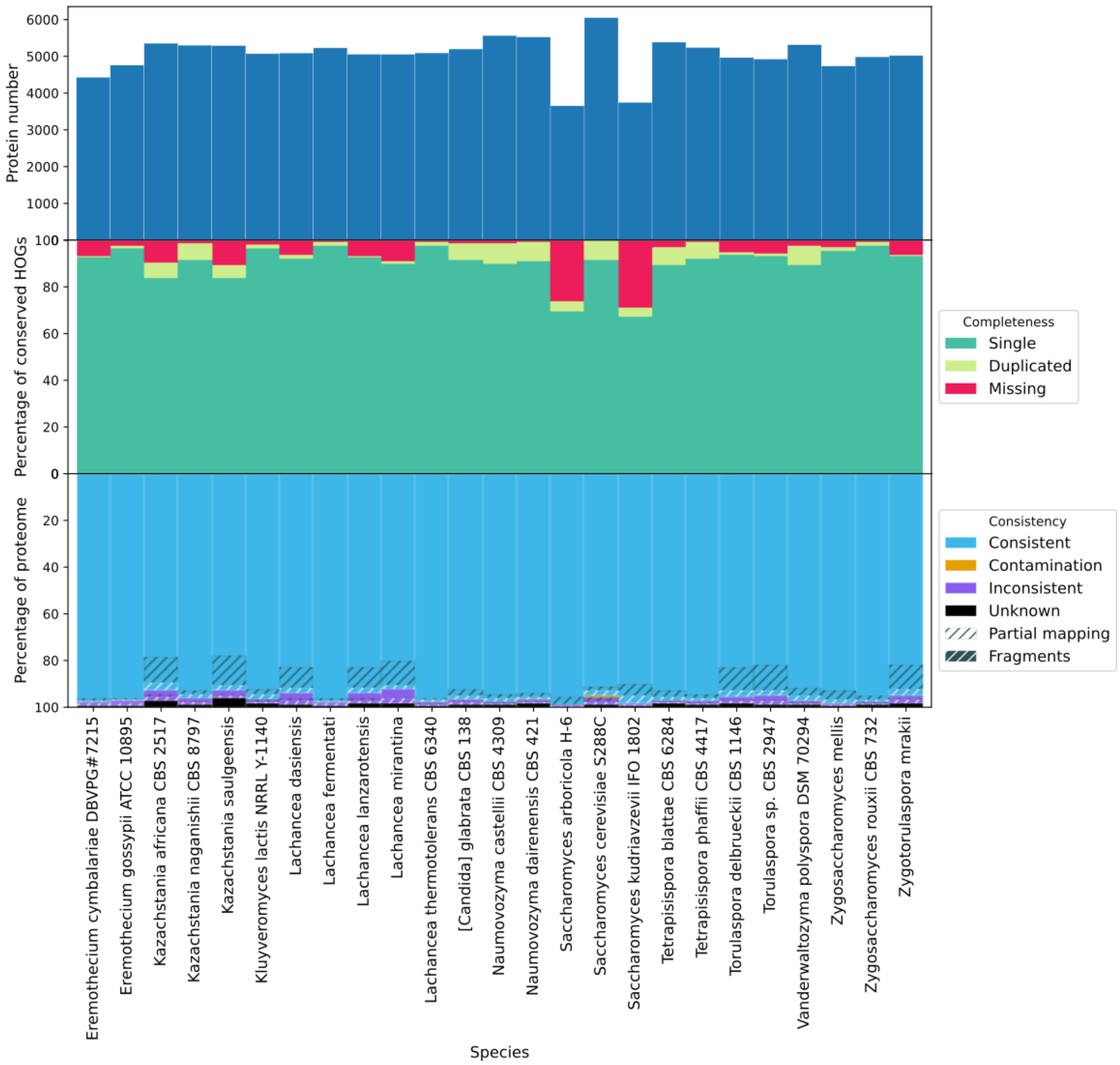


**Supplementary Figure 18. OMArk results for proteomes from the Saccharomyces genus.**

#### *Arabidopsis thaliana* and Brassicaceae

We compare *Arabidopsis thaliana* to other species in the Brassicaceae clade (Supplementary Figure 19). As expected, its proteome ranks among the most complete in the dataset according to OMArk (98.71% completeness), along with both subspecies of *Brassica rapa* (98.6% and 98.26%). The same three proteomes are also among the best in terms of taxonomically and structurally Consistent genes (90.32% for *A. thaliana,* 88.14% for *B. rapa*, 92.41% for *B. rapa subspecies pekinensis*. Every species in this clade displays some amount of duplicated genes, especially high among species from the *Brassica* genus, which is likely due to high ploidy levels in those species (Yang et al. 2016). Species from the same genus display heterogeneous levels of completeness and/or different consistency. As an example, even though *Arabidopsis lyrata* is fairly complete (97.95%), its proportion of taxonomically and structurally Consistent genes is

80.91%, nearly 10% lower than its sister species.

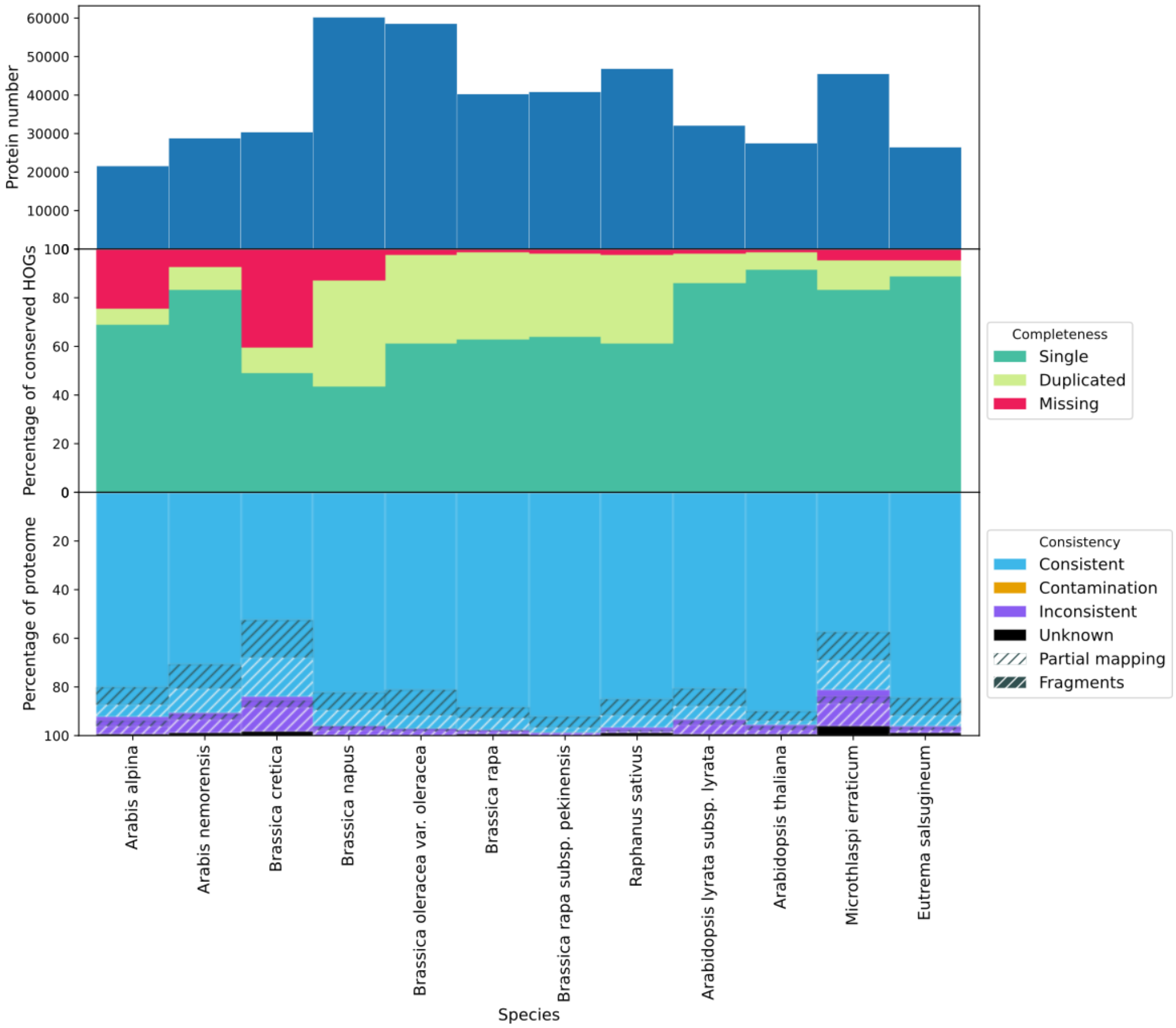


**Supplementary Figure 19. OMArk results for proteomes from the Brassicaceae clade.**

### Assembly and annotation comparison

We obtained the list of species with either annotation or assembly change for the two previous releases of Ensembl metazoa (53 and 54) (Yates et al. 2022), and computed OMArk results for the previous and newly released version of each one. This corresponds to 18 pairs of proteomes from protostomian species, 11 resulting from an assembly change and 7 (only nematode species) from an annotation change. All the results are available in Supplementary Table 5 and shown in Supplementary Figure 19-36.

#### Assembly comparisons

Changes in proteomes resulting from a different assembly manifest as important differences in the number of reported protein coding genes, with all but one leading to a change in number of more than 500 genes. In OMArk, this is reflected as a minor decrease in completeness (less than 2%) for 4 species, a minor increase for 2 species, and an increase of several percent for the

5 others, up to 22.94% in the most extreme case (*Teleopsis dalmanni*) (Supplementary Table 5, Supplementary Figures 20-30).

Interestingly, increase in completeness was not always associated with an increase in the number of proteins, with *Glossina fuscipes* gaining 5% of completeness despite having 7,730 fewer protein coding genes and *Bombyx mori* gaining 3.78% completeness with 742 fewer genes. Another notable case is the latest *Acyrthosiphon pisum* assembly is only 0.64 less complete with 17,918 less genes.

The newest assembly also led to a decrease in proportion of *Duplicate* conserved genes in all but three cases, likely corresponding to a decrease in the proportion of fragmented genes or improved phasing with better genome coverage. For the other three species, the increase in *D*uplicated genes is slight for *Pristionchus pacificus* (1.51%) and *Schistosoma mansoni* (0.54%) but significant for *Teleopsis dalmanni* (+11.18%). As the selected conserved genes are not supposed to be conserved in a single copy, this is not enough to conclude in introduced errors in the novel assembly, but the later increase being so drastic is likely indicative of fragments or spurious duplicates.

In terms of consistency assessment, the new assembly leads to a higher proportion of Taxonomically Consistent genes in 6 cases out of 11. The highest increases are detected in species for which the number of proteins decrease the most, namely: *Acyrthosiphon pisum* (-17,918 proteins, +8.43% Taxonomically Consistent genes), *Glossina fuscipes* (-7,730 proteins, +15.57% Taxonomically Consistent genes), *Bombus impatiens* (-5,264 proteins, +7.30% Taxonomically Consistent genes) and *Danaus plexippus plexippus* (-2,023 proteins, +4.32% Taxonomically Consistent genes). Together with the reported increase or low decrease in completeness, this indicates that the newest assembly leads to a better quality gene set overall.

In contrast, the updated assembly of *Crassostrea gigas* leads to a decrease in the proportion of Taxonomically Consistent of genes, together with an increase in number of genes and completeness (+4,239 proteins, -3.71% Taxonomically Consistent genes) meaning that as a whole, the added genes show lower agreement with known Lophotrochozoan gene families and contain likely artifactual sequences.

The signal of gene set quality improvement on most assemblies is even stronger when considering the proportion of taxonomically and structurally consistent genes which show an increase in 9 out of 11 cases and increases more in value than the taxonomically consistent genes alone in all but one case (*Crassostrea gigas*), showing that new assemblies often result in more structurally accurate gene models (Supplementary Table 7).

Finally, the contamination detected by OMArk in different assemblies of the same species varies between assemblies. For *Bombus impatiens* and *Schistosoma mansoni*, the contamination is no longer detected in the gene set of the updated assembly. In the cases of *Acyrthosiphon pisum* and *Glossina fuscipes,* the number of detected contaminants is reduced in the newer assembly, implying that either some contamination still exists in the current or there is a case of horizontal gene transfer in these genomes. We notice one case (*Teleopsis dalmanni*) where OMArk detects contamination from species of the Bacteriales taxa, only on the updated assembly, indicating contamination introduced during resequencing.

| Species | Completeness change (%) | Structurally consistent gene change (%) |
|---|---|---|
| *Solenopsis invicta* | -0.23 | 3.1 |
| *Bombus impatiens* | -1.23 | 16.73 |
| *Glossina fuscipes* | 5.02 | 29.16 |
| *Pristionchus pacificus* | 4.44 | 1.51 |
| *Sarcoptes scabiei* | -0.76 | -0.53 |
| *Bombyx mori* | 3.78 | 2.31 |
| *Danaus plexippus plexippus* | 0.40 | 13.00 |
| *Acyrthosiphon pisum* | -0.64 | 18.82 |
| *Schistosoma mansoni* | 0.33 | 2.06 |
| *Crassostrea gigas* | 4.67 | -4.25 |
| *Teleopsis dalmanni* | 22.94 | 8.90 |

**Supplementary Table 7. Summary table of OMArk quality assessment of percentage of change from previous to current assembly release of Ensembl Metazoa**. Most proteomes increased in completeness orconsistency in the newest release. Green color indicates an improvement, and red indicates a decrease in quality, based on OMArk.

By providing statistics about presence of conserved genes and consistency of the overall gene sets, OMArk allows detection of change along both axes. Over all proteomes, OMArk detects an increase in either completeness and proportion of taxonomically and structurally consistent genes in the new assembly in all but one case *(Sarcoptes scabiei*). Considering both axes can be particularly useful when they give contradicting information: for example, we notice that the updated proteome of *Bombus impatiens* is marked as slightly less complete than the previous version, but has higher consistency measures and is no longer contaminated. However, *Crassostrea gigas*' newer proteome is more complete but exhibits an overall lower consistency. This nuance would be lost when comparing completeness alone.

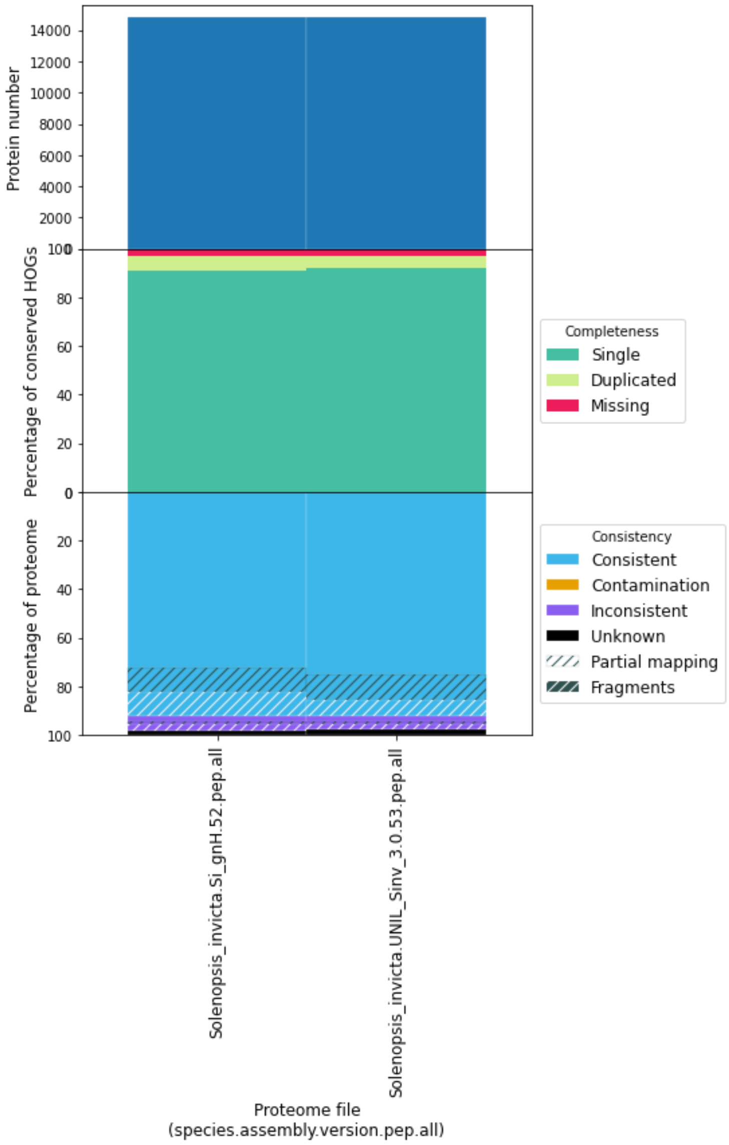


**Supplementary Figures 20. OMArk comparison of different versions in *Solenopsis invicta* assemblies.** Left bar plot corresponds to the proteome version in Ensembl Metazoa 52. Right bar plot is the proteome available in Ensembl Metazoa 53 and onward.

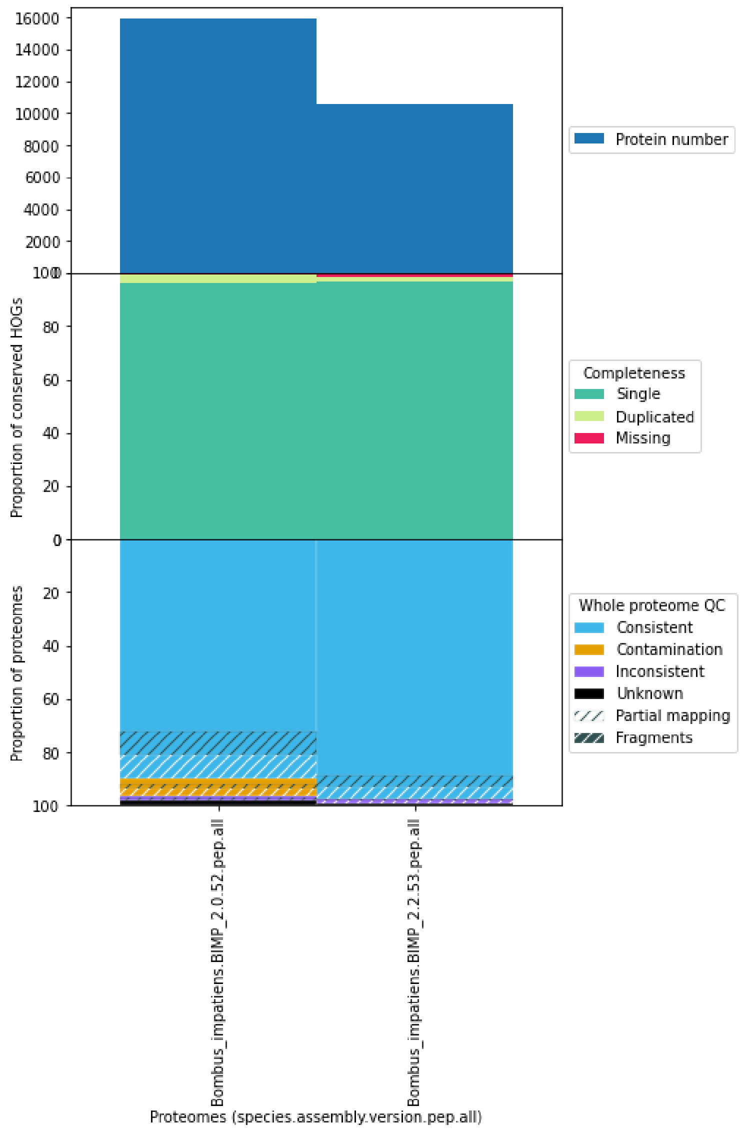


**Supplementary Figures 21. OMArk comparison of different versions in *Bombus impatiens* assemblies.** Left bar plot corresponds to the proteome version in Ensembl Metazoa 52. Right bar plot is the proteome available in Ensembl Metazoa 53 and onward

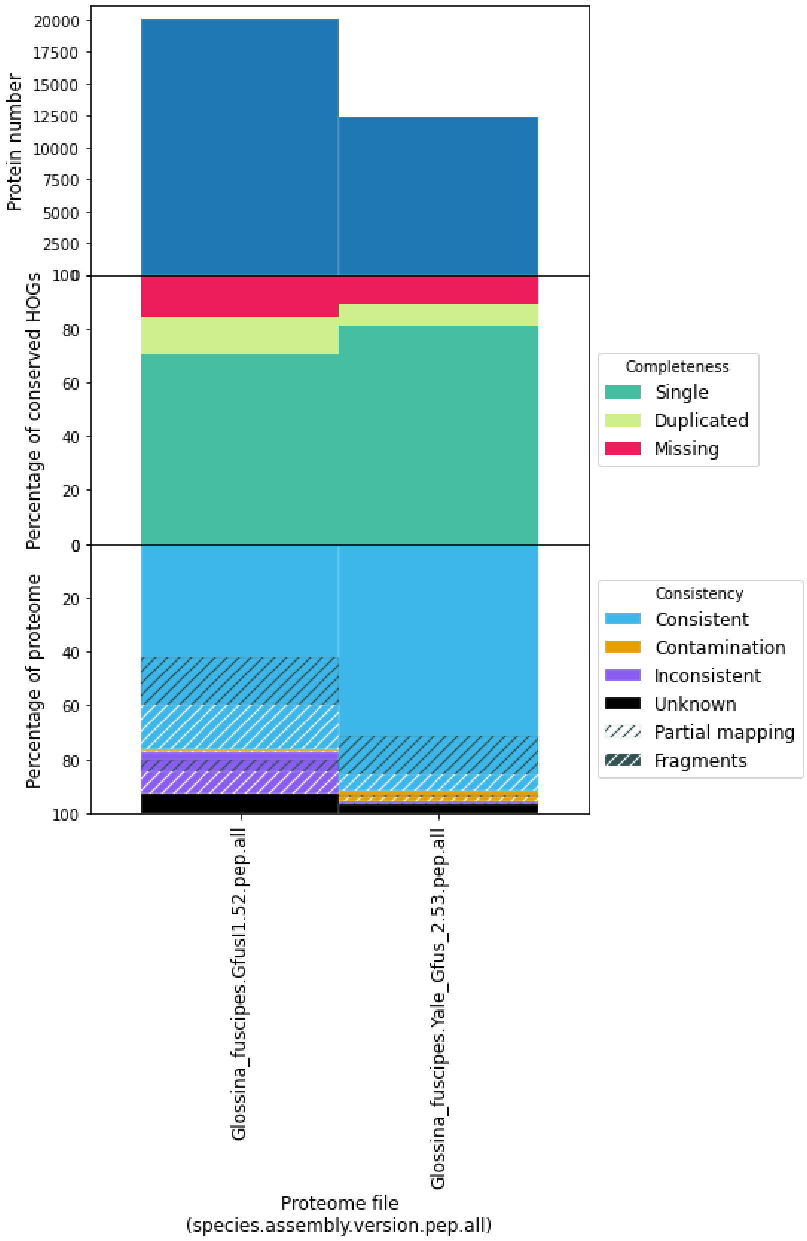


**Supplementary Figures 22. OMArk comparison of different versions in *Glossina fuscipes* assemblies.** Left bar plot corresponds to the proteome version in Ensembl Metazoa 52. Right bar plot is the proteome available in Ensembl Metazoa 53 and onward

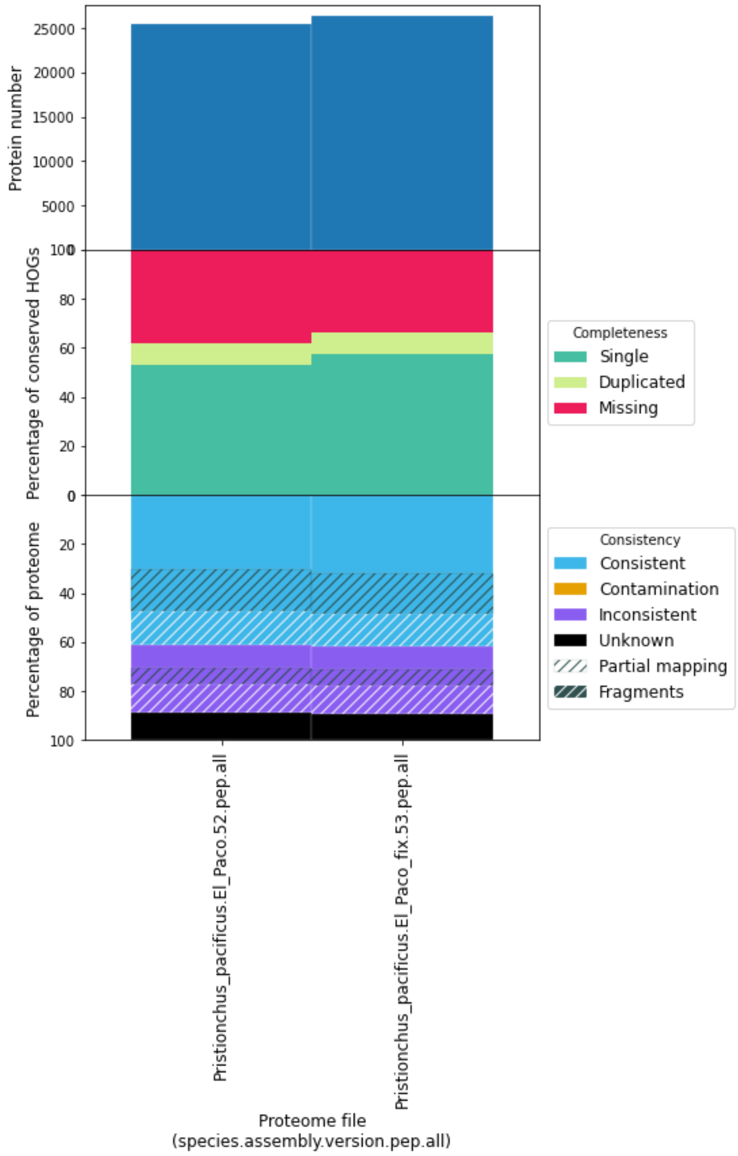


**Supplementary Figures 23. OMArk comparison of different versions in *Pristionchus pacificus* assemblies.** Left bar plot corresponds to the proteome version in Ensembl Metazoa 52. Right bar plot is the proteome available in Ensembl Metazoa 53 and onward

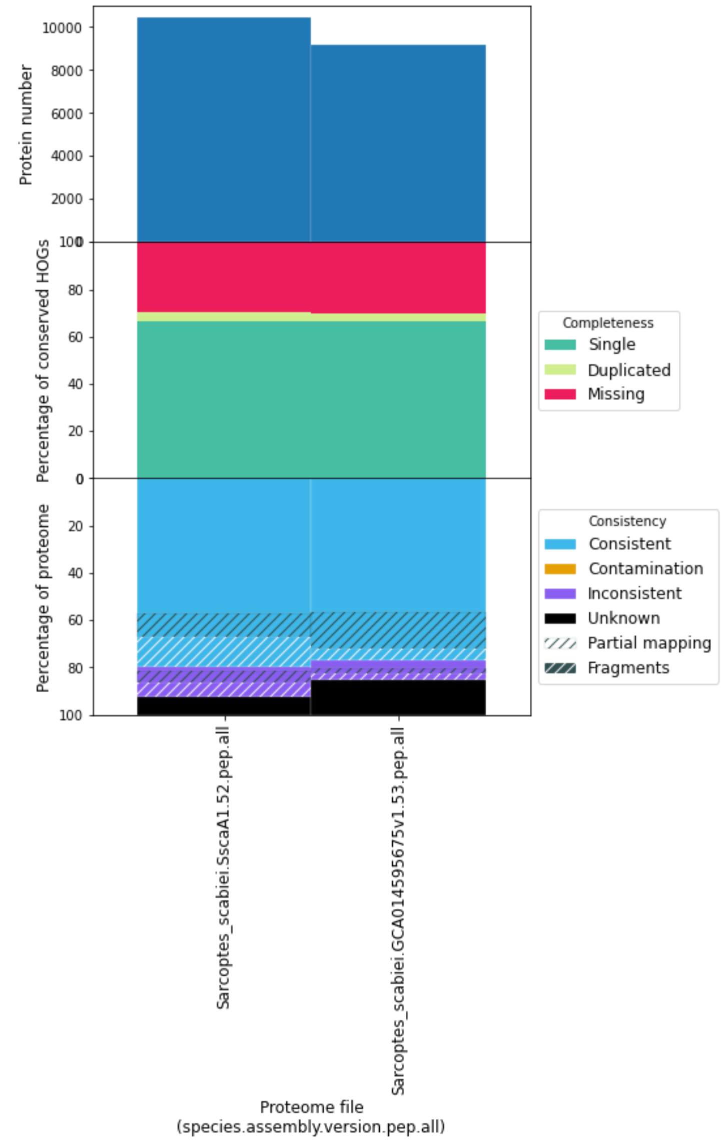


**Supplementary Figures 24. OMArk comparison of different versions in *Sarcoptes scabiei* assemblies.** Left bar plot corresponds to the proteome version in Ensembl Metazoa 52. Right bar plot is the proteome available in Ensembl Metazoa 53 and onward

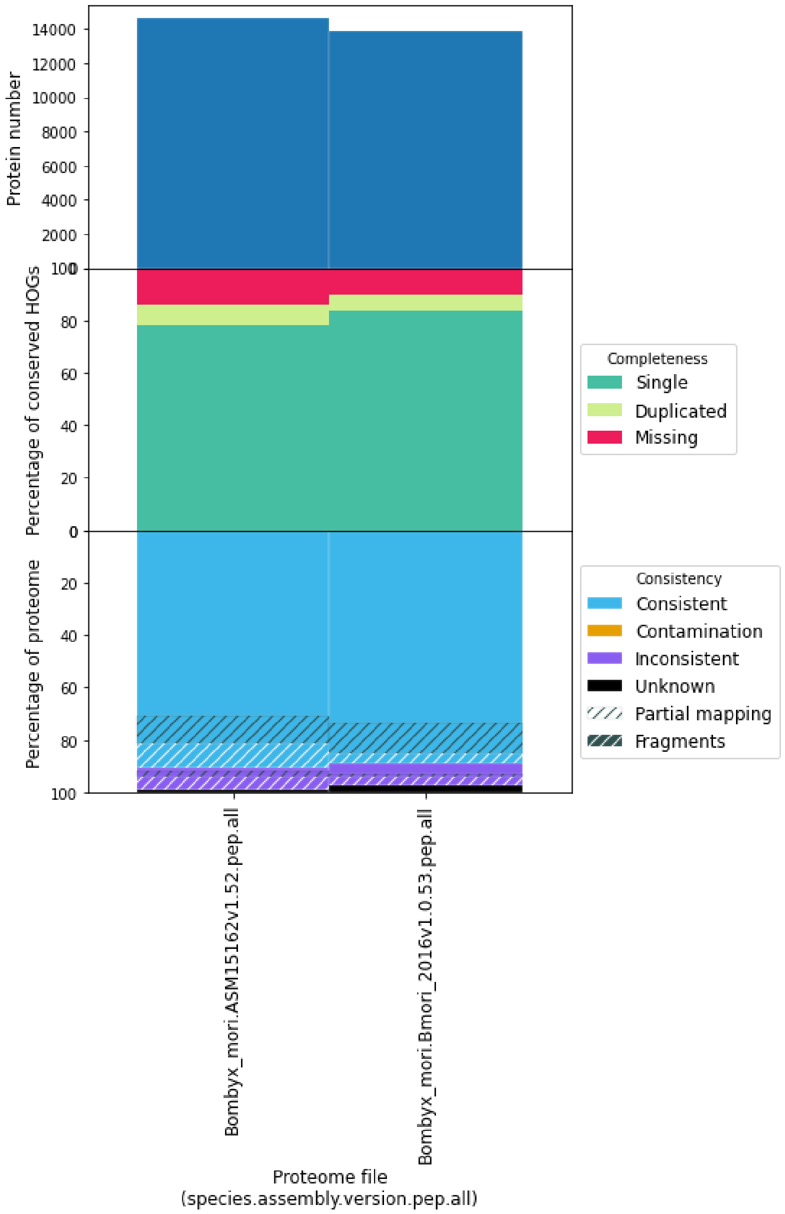


**Supplementary Figures 25. OMArk comparison of different versions in *Bombyx mori* assemblies.** Left bar plot corresponds to the proteome version in Ensembl Metazoa 52. Right bar plot is the proteome available in Ensembl Metazoa 53 and onward

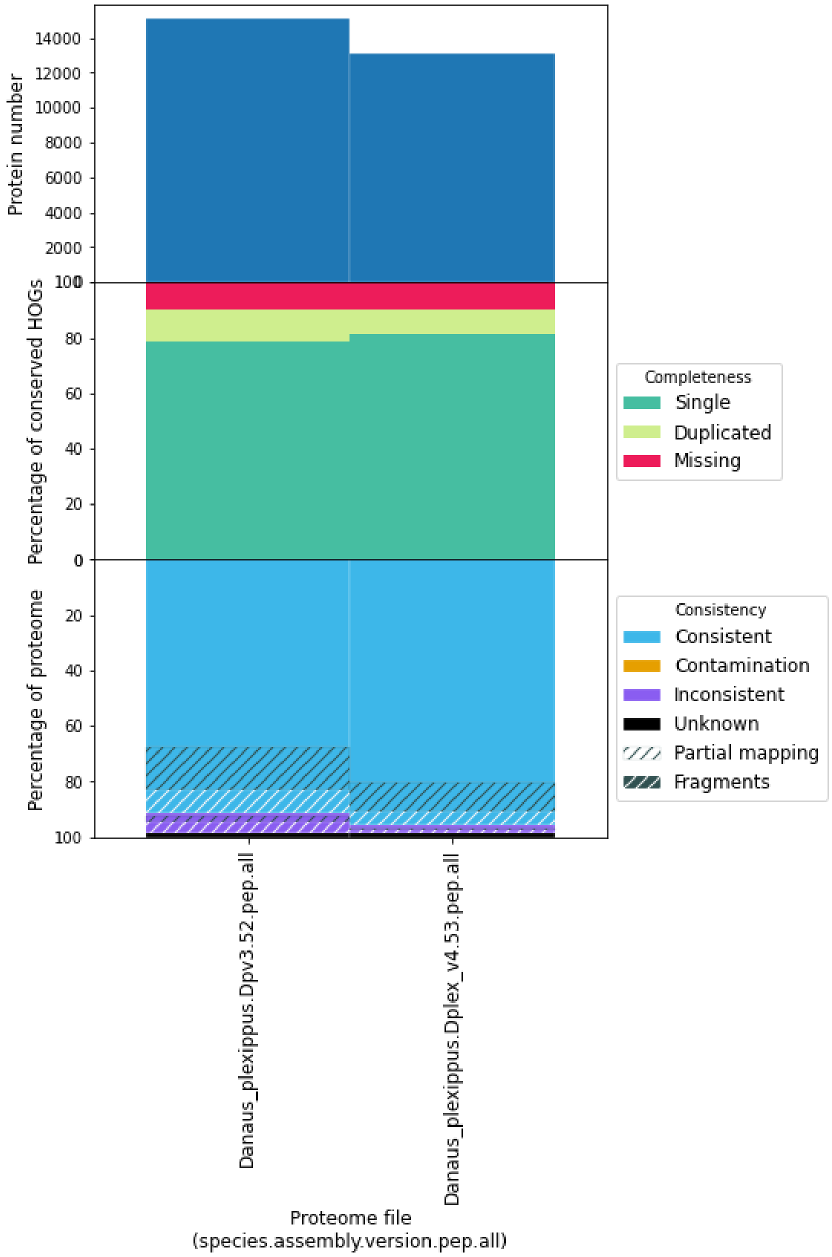


**Supplementary Figures 26. OMArk comparison of different versions in *Danaus plexippus* assemblies.** Left bar plot corresponds to the proteome version in Ensembl Metazoa 52. Right bar plot is the proteome available in Ensembl Metazoa 53 and onward

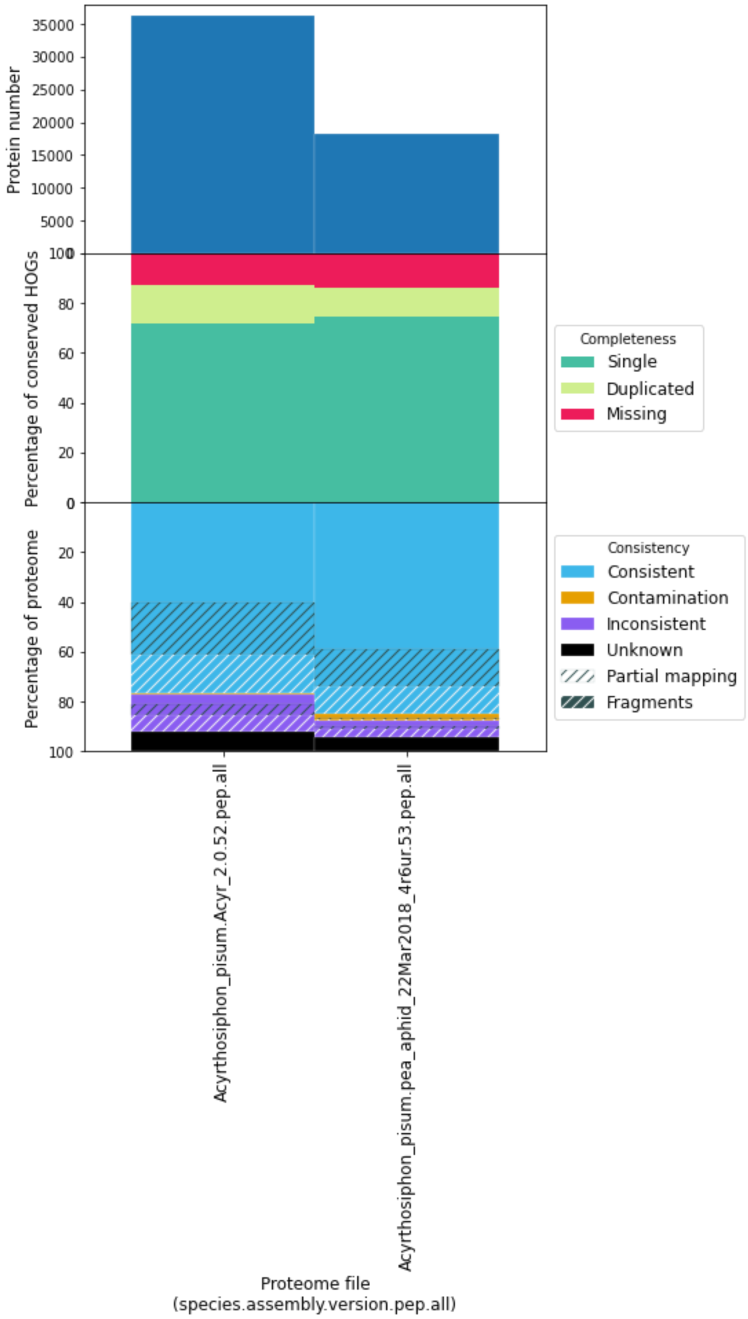


**Supplementary Figures 27. OMArk comparison of different versions in *Acyrthosiphon pisum* assemblies.** Left bar plot corresponds to the proteome version in Ensembl Metazoa 52. Right bar plot is the proteome available in Ensembl Metazoa 53 and onward

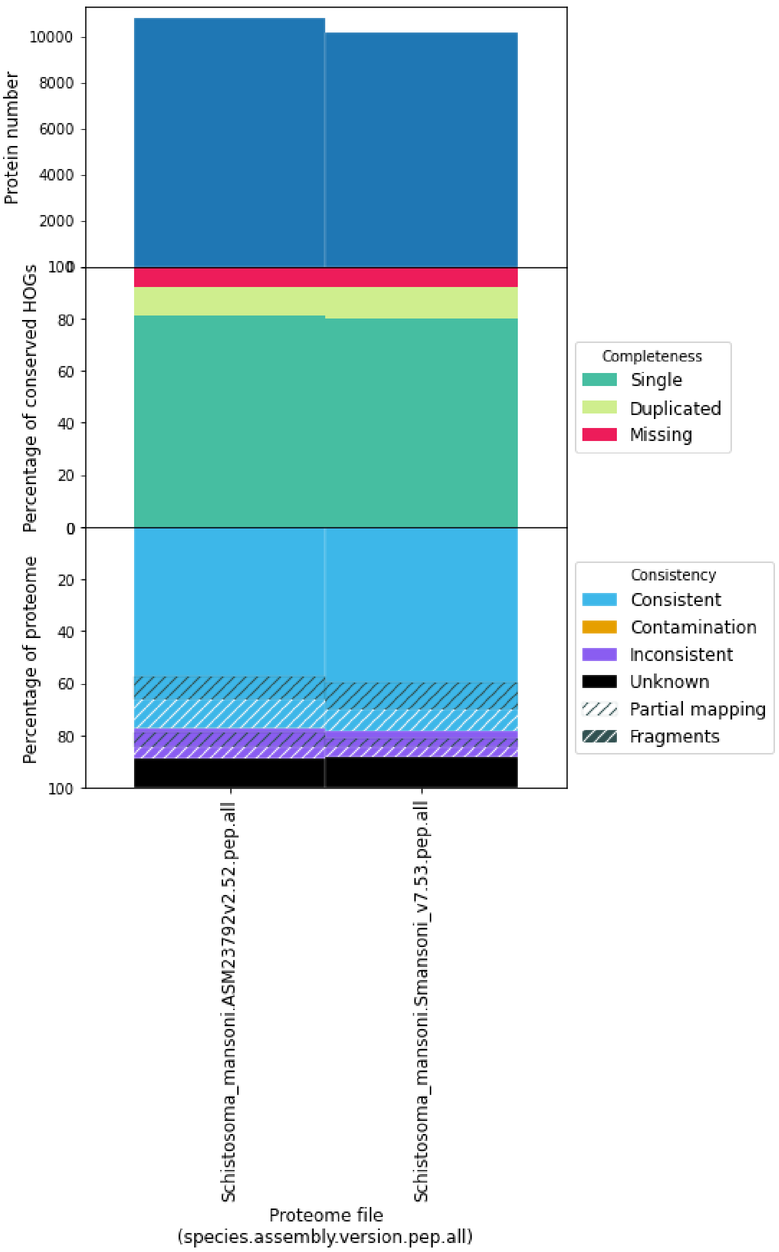


**Supplementary Figures 28. OMArk comparison of different versions in *Schistosoma mansoni* assemblies.** Left bar plot corresponds to the proteome version in Ensembl Metazoa 52. Right bar plot is the proteome available in Ensembl Metazoa 53 and onward

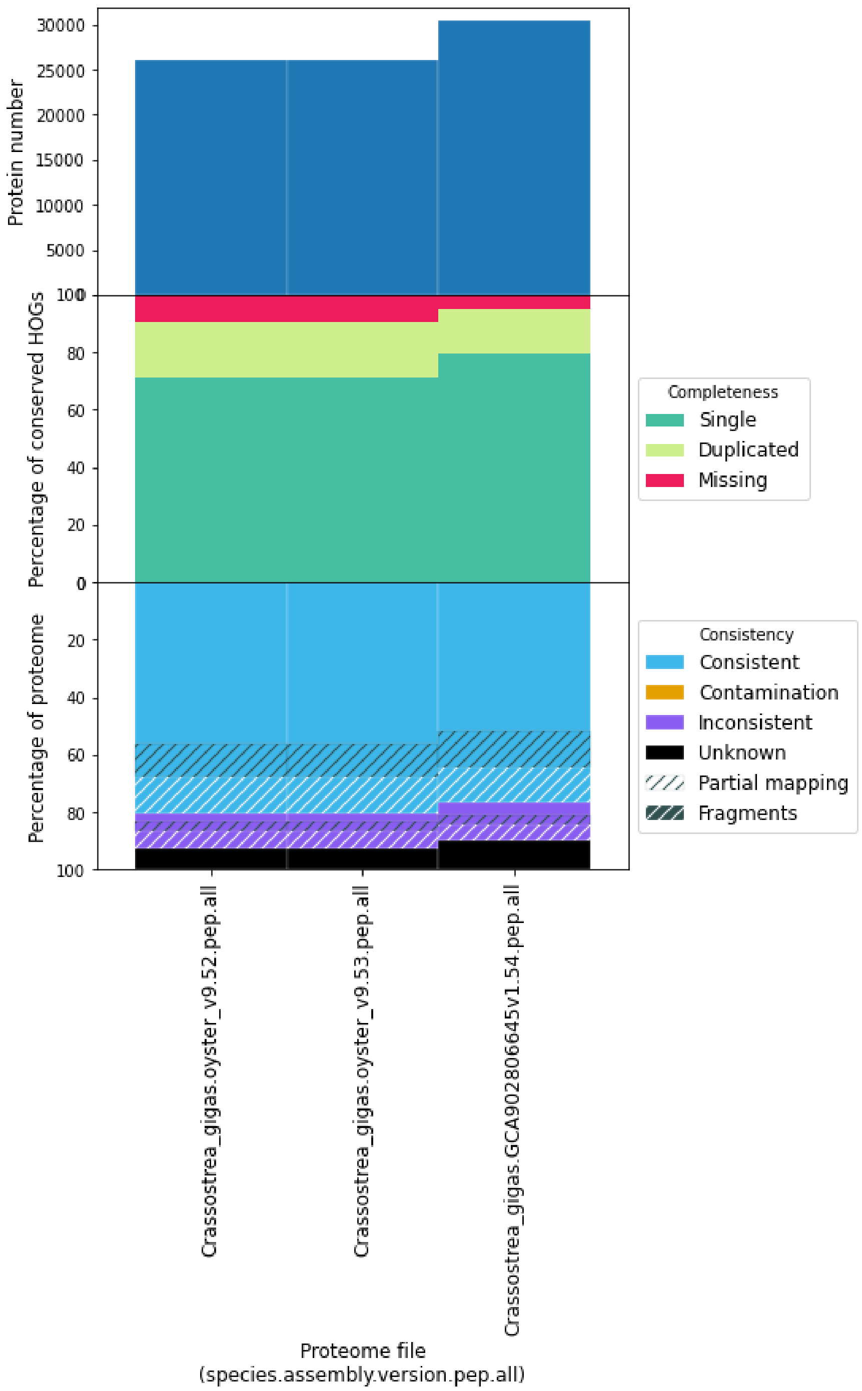


**Supplementary Figures 29. OMArk comparison of different versions in Crassostrae gigas assemblies.** Left bar plot corresponds to the proteome version in Ensembl Metazoa 52. Central bar plot is the proteome available in Ensembl Metazoa 53. Right bar plotis the newest assembly available in Ensembl Metazoa 54.

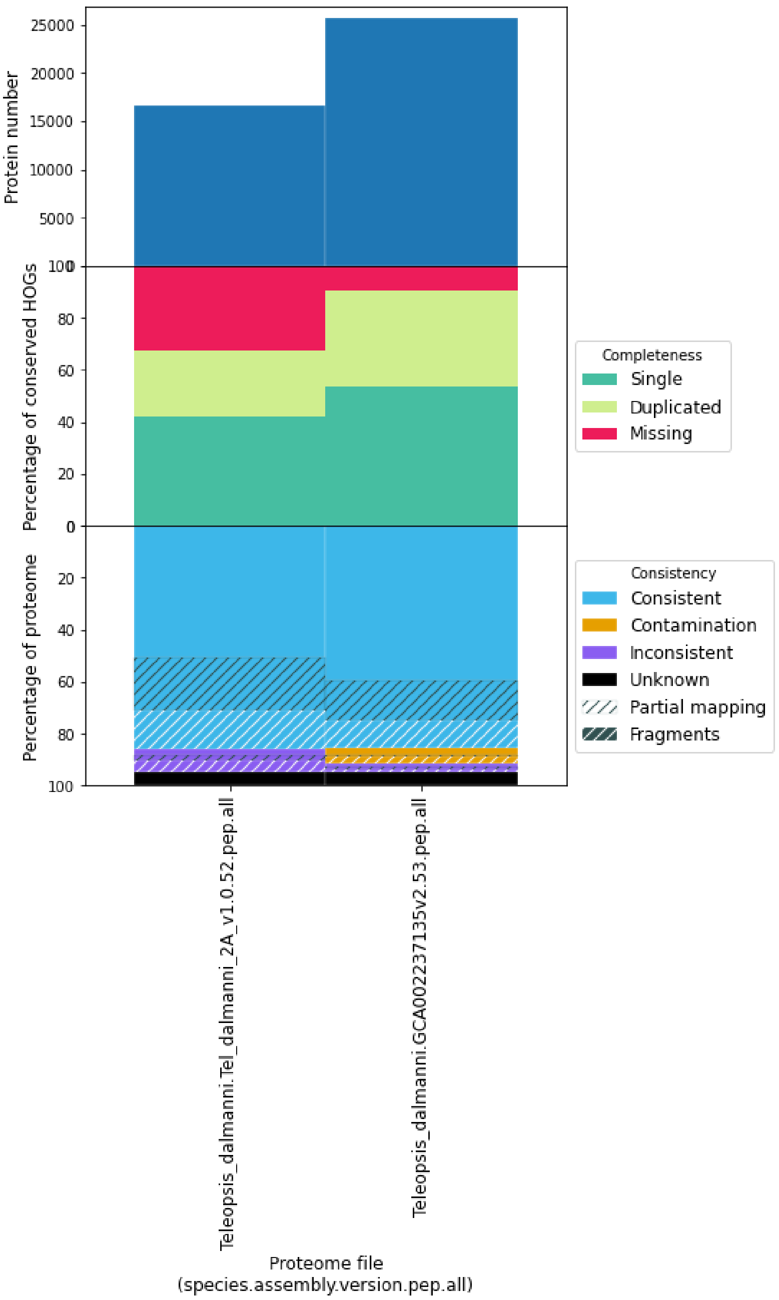


**Supplementary Figures 30. OMArk comparison of different versions in *Teleopsis dalmanni* assemblies.** Left bar plot corresponds to the proteome version in Ensembl Metazoa 52. Right bar plot is the proteome available in Ensembl Metazoa 53 and onward.

#### Annotation comparisons

The species with updated annotations in our dataset were all nematodes. The change in the gene set between annotations was minor according to OMArk results (few metrics with a change higher than 1%) and in terms of coding-gene number, likely reflective of iterative changes (Supplementary Figures 31-37). Absolute change in gene number was higher than 10 for 4 of the 7 proteomes, namely: *Caenorhabditis brenneri* (+30), *Brugia malayi* (-132), *Caenorhabditis briggsae* (-175) and *Caenorhabditis elegans* (-195).

Surprisingly, the change in gene content of the three latter species leads to a slightly lower completeness as detected by OMArk (respectively, -0.03%, -0.05% and -0.16%), meaning some of the removed genes correspond to ones that are presumably conserved within the lineage. However, the observed changes in the same species is more important in the Duplicated category (respectively, -0.10%, -0.08% and -0.18%), hinting that part of this change is due to removal of spuriously duplicated genes. In the opposite direction, the added genes in *Caenorhabditis brenneri* lead to slight increase in completeness (0.02%) and a more important increase in duplicated genes (0.06%). The change in annotation in other species has a comparatively lower impact on OMArk statistics.

The change in gene set has a small impact on Taxonomical consistency, but a noticeable one for *C. elegans* (+0.12%) and *Brugia malayii* (+0.42%) meaning it leads to better quality metrics overall. This is also the case for taxonomically and structurally consistent proteins, where it increases by 0.12% for *C. elegans*, by 0.4% for *C. briggsea* and by 1.21% for *Brugia malayi*.

As expected, given no changes were made to the assembly, we noticed no significant differences in terms of contaminant for these species, with *Caenorhabditis japonica* being contaminated by a *Paenibacillus* species and *Caenorhabditis remanei* being contaminated with *Acinetobacter* and *Enterobacteriacea* species in both annotation according to OMArk.

To conclude, we show that while iterative changes in the annotation do not lead to large changes in OMArk statistics, it still provides information about the nature of these changes when used in a one-to-one comparison.

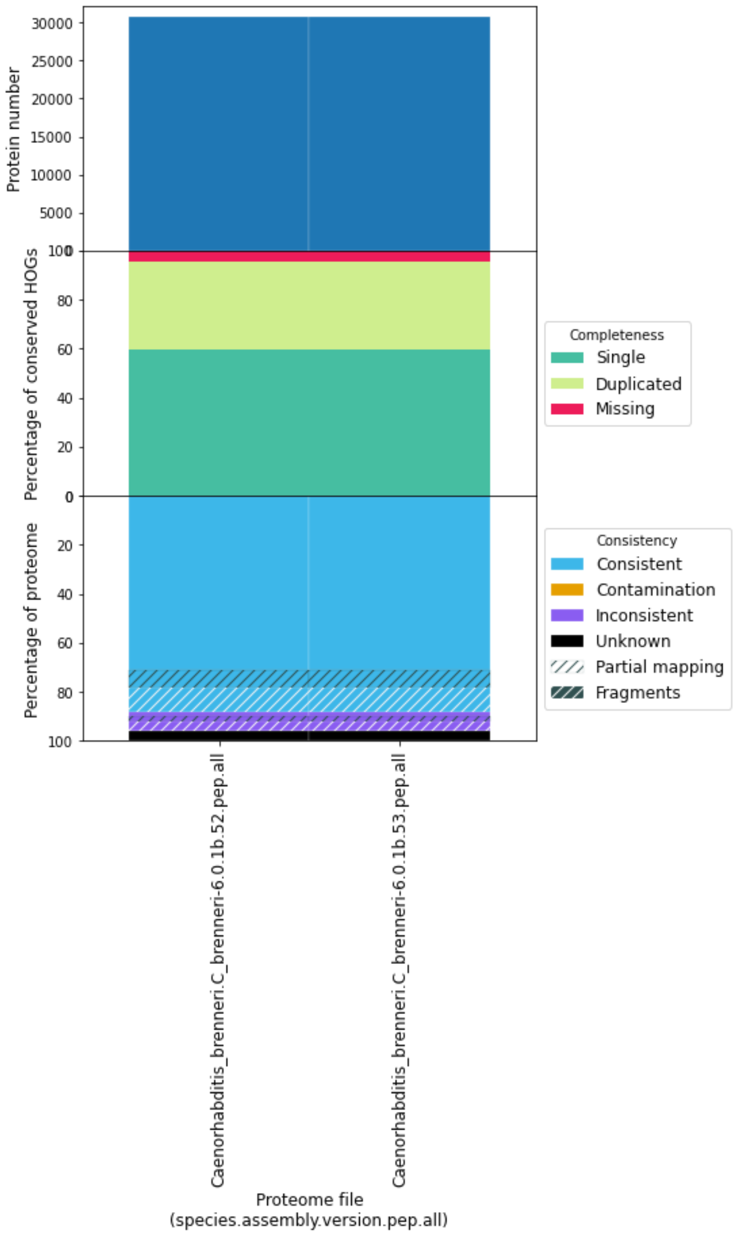


**Supplementary Figures 31. OMArk comparison of different versions in *Caenorhabditis brenneri* annotation.** Left bar plot corresponds to the proteome version in Ensembl Metazoa 52. Right bar plot is the proteome available in Ensembl Metazoa 53 and onward.

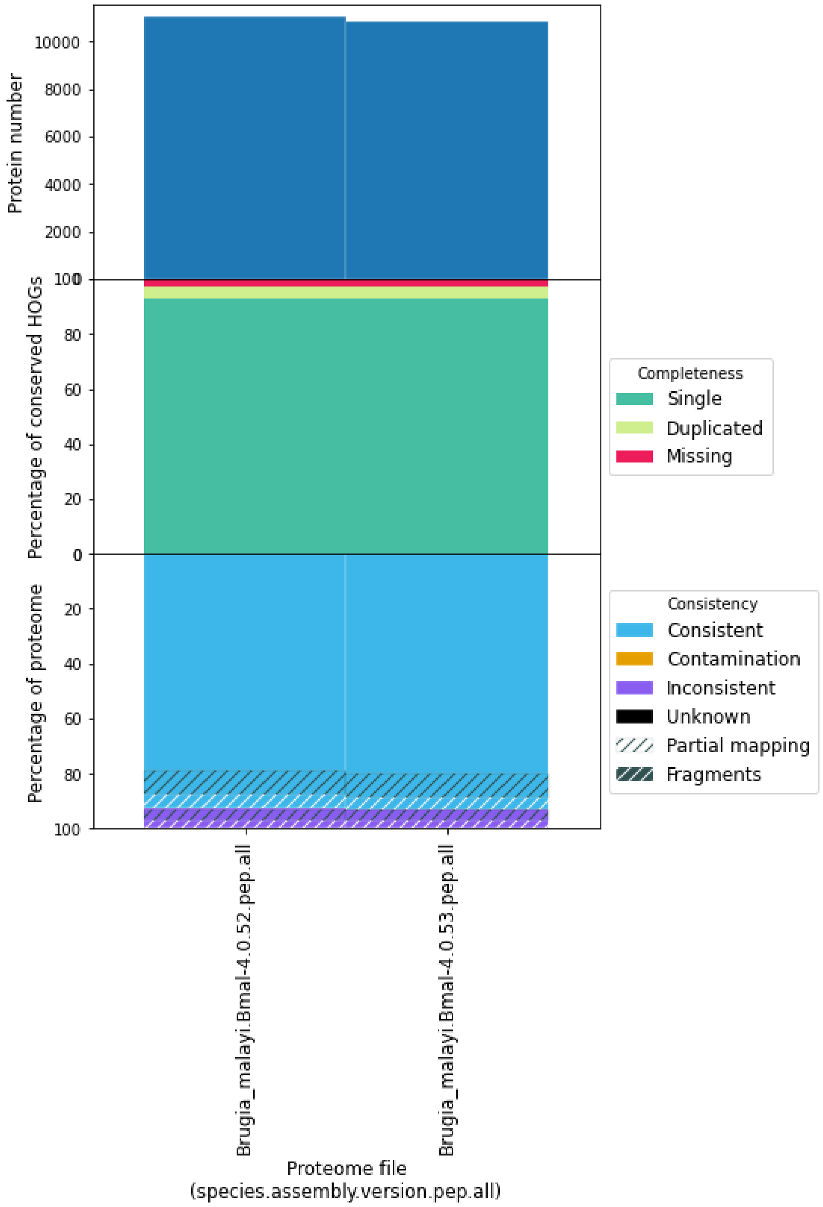


**Supplementary Figures 32. OMArk comparison of different versions in *Brugia malayi* annotation.** Left bar plot corresponds to the proteome version in Ensembl Metazoa 52. Right bar plot is the proteome available in Ensembl Metazoa 53 and onward.

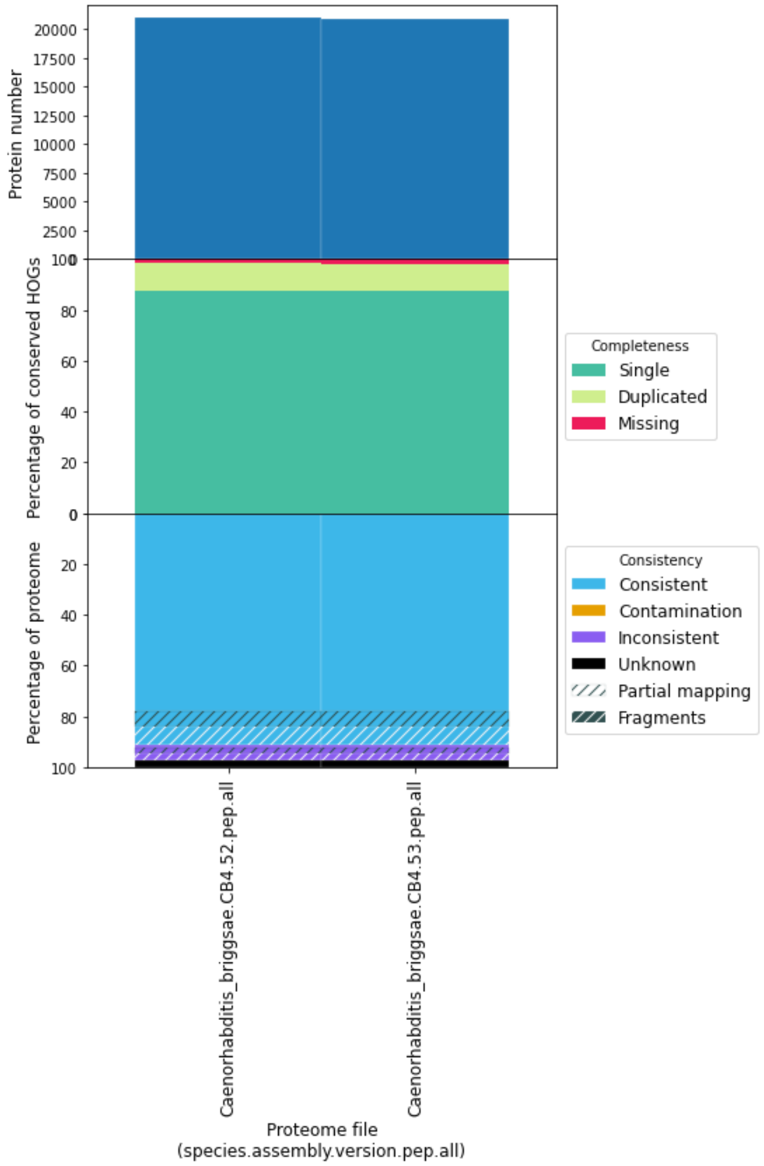


**Supplementary Figures 33. OMArk comparison of different versions in *Caenorhabditis briggsae* annotation.** Left bar plot corresponds to the proteome version in Ensembl Metazoa 52. Right bar plot is the proteome available in Ensembl Metazoa 53 and onward.

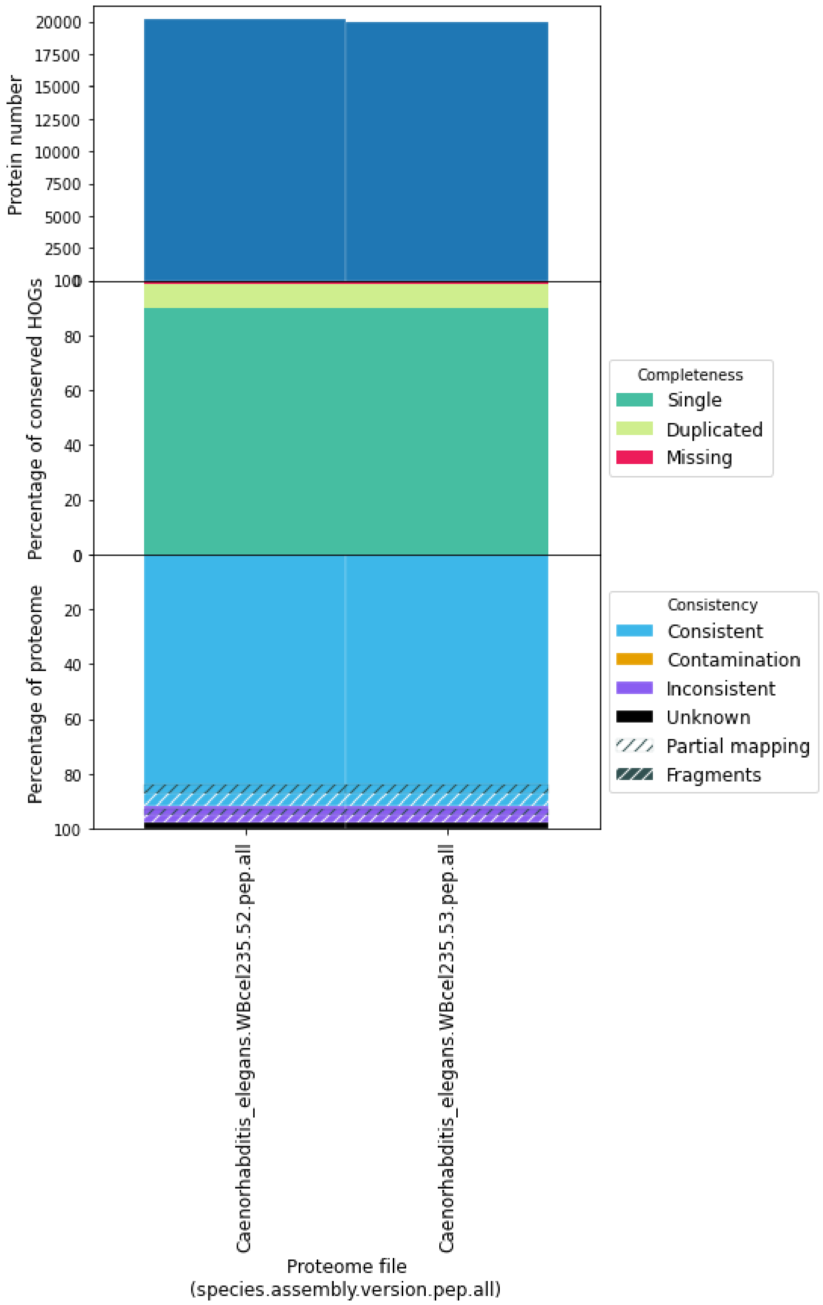


**Supplementary Figures 34. OMArk comparison of different versions in *Caenorhabditis elegans* annotation.** Left bar plot corresponds to the proteome version in Ensembl Metazoa 52. Right bar plot is the proteome available in Ensembl Metazoa 53 and onward.

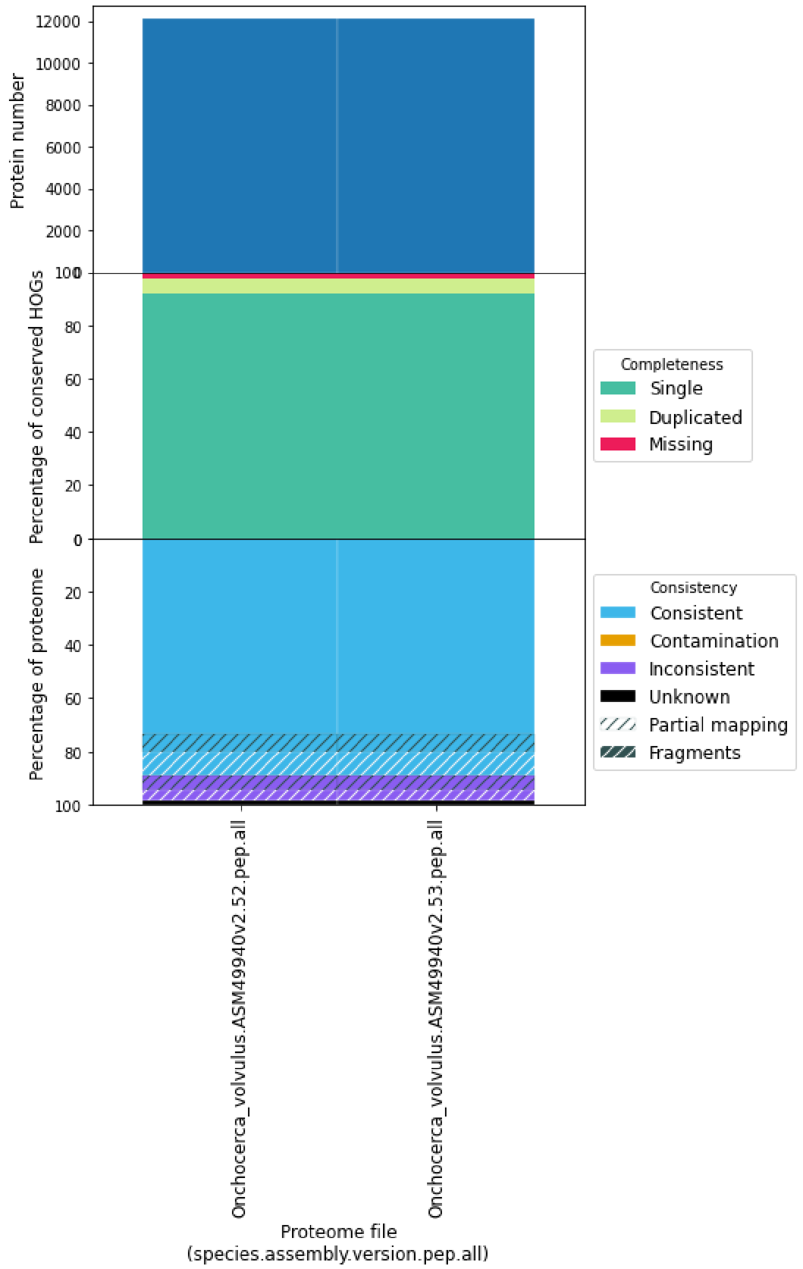


**Supplementary Figures 35. OMArk comparison of different versions in *Onchocerca volvulus* annotation.** Left bar plot corresponds to the proteome version in Ensembl Metazoa 52. Right bar plot is the proteome available in Ensembl Metazoa 53 and onward.

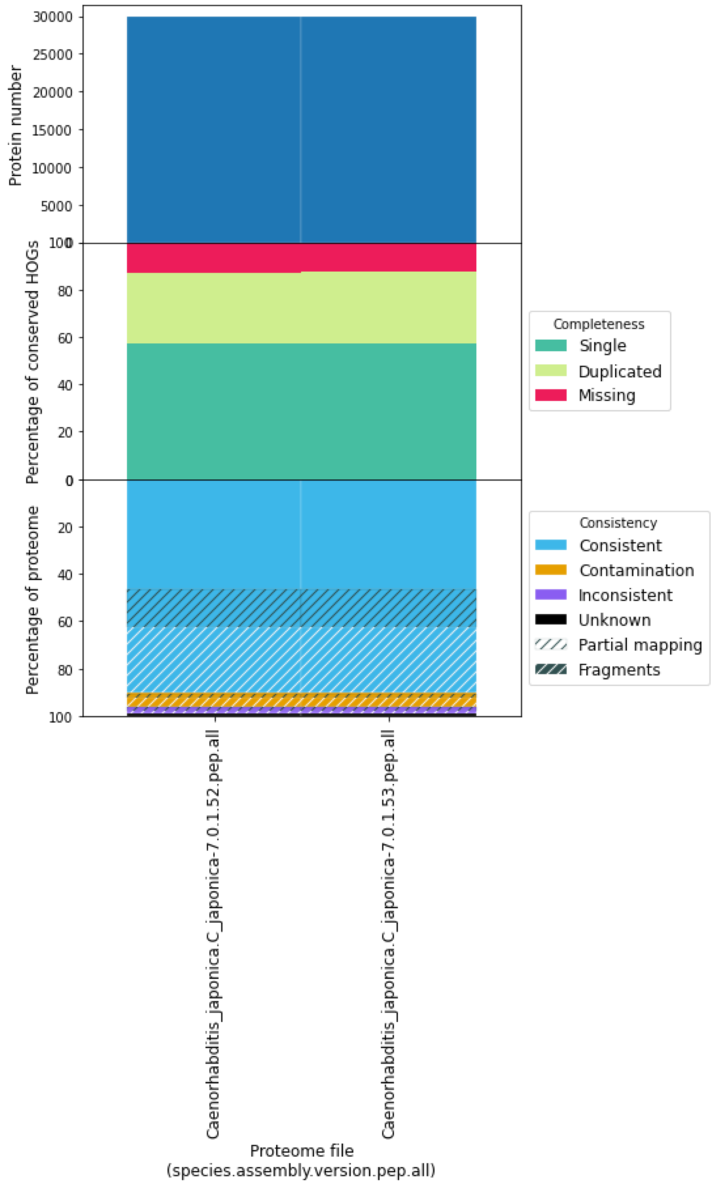


**Supplementary Figures 36. OMArk comparison of different versions in *Caenorhabditis japonica* annotation.** Left bar plot corresponds to the proteome version in Ensembl Metazoa 52. Right bar plot is the proteome available in Ensembl Metazoa 53 and onward.

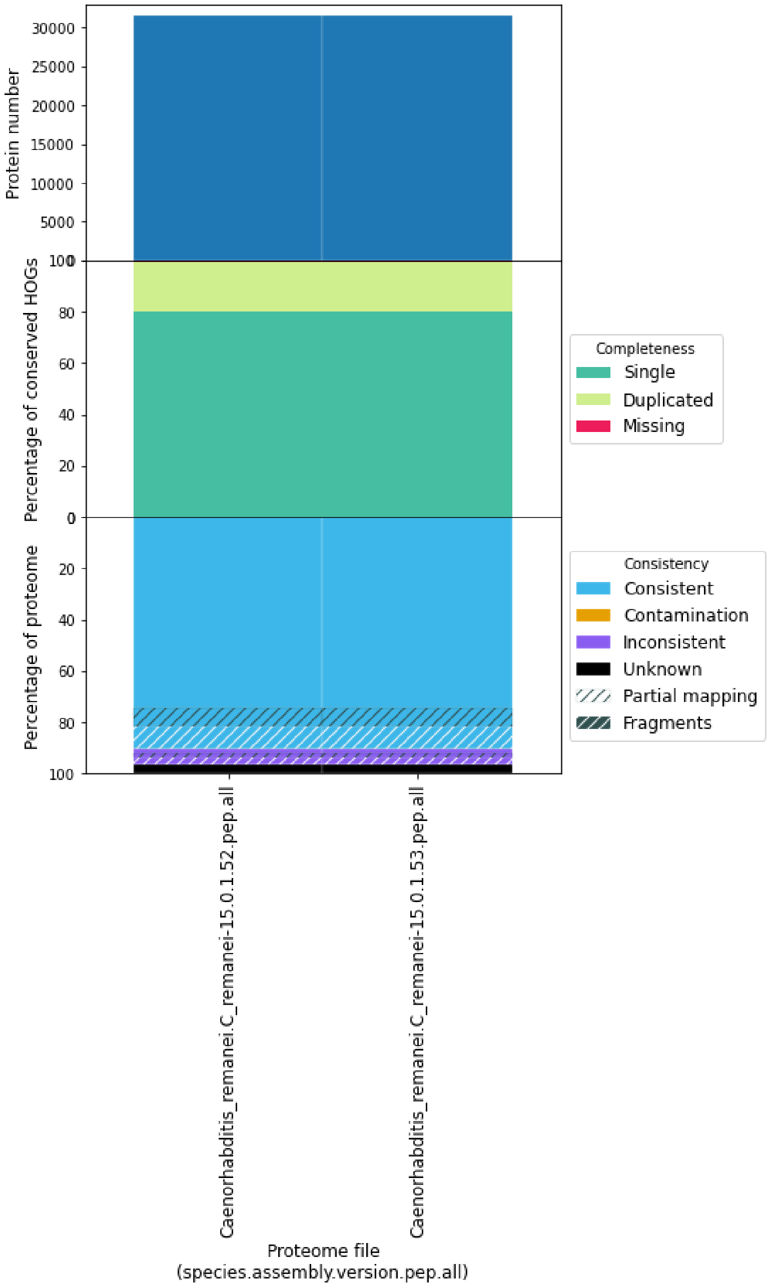


**Supplementary Figures 37. OMArk comparison of different versions in *Caenorhabditis remanei* annotation.** Left bar plot corresponds to the proteome version in Ensembl Metazoa 52. Right bar plot is the proteome available in Ensembl Metazoa 53 and onward.

Analysis of Hibiscus Syriacus Provides Insights of Polyploidization and Indeterminate Flowering in Woody Plants.” *DNA Research: An International Journal for Rapid Publication of Reports on Genes and Genomes* 24 (1): 71–80. https://doi.org/10.1093/dnares/dsw049.

Lhee, Duckhyun, Ji-San Ha, Sunju Kim, Myung Gil Park, Debashish Bhattacharya, and Hwan Su Yoon. 2019. “Evolutionary Dynamics of the Chromatophore Genome in Three Photosynthetic Paulinella Species.” *Scientific Reports* 9 (1): 2560. https://doi.org/10.1038/s41598-019-38621-8.

Maeda, Yoshiaki, Kahori Watanabe, Ryosuke Kobayashi, Tomoko Yoshino, Chris Bowler, Mitsufumi Matsumoto, and Tsuyoshi Tanaka. 2021. “Chromosome Scale Assembly of Allopolyploid Genome of the Diatom Fistulifera Solaris.” *bioRxiv*. https://doi.org/10.1101/2021.11.10.468027.

Manni, Mosè, Matthew R. Berkeley, Mathieu Seppey, Felipe A. Simão, and Evgeny M. Zdobnov. 2021. “BUSCO Update: Novel and Streamlined Workflows along with Broader and Deeper Phylogenetic Coverage for Scoring of Eukaryotic, Prokaryotic, and Viral Genomes.” *Molecular Biology and Evolution* 38 (10): 4647–54. https://doi.org/10.1093/molbev/msab199.

Nakjang, Sirintra, Tom A. Williams, Eva Heinz, Andrew K. Watson, Peter G. Foster, Kacper M. Sendra, Sarah E. Heaps, Robert P. Hirt, and T. Martin Embley. 2013. “Reduction and Expansion in Microsporidian Genome Evolution: New Insights from Comparative Genomics.” *Genome Biology and Evolution* 5 (12): 2285–2303. https://doi.org/10.1093/gbe/evt184.

O’Leary, Nuala A., Mathew W. Wright, J. Rodney Brister, Stacy Ciufo, Diana Haddad, Rich McVeigh, Bhanu Rajput, et al. 2016. “Reference Sequence (RefSeq) Database at NCBI: Current Status, Taxonomic Expansion, and Functional Annotation.” *Nucleic Acids Research* 44 (D1): D733–45. https://doi.org/10.1093/nar/gkv1189.

Rhie, Arang, Shane A. McCarthy, Olivier Fedrigo, Joana Damas, Giulio Formenti, Sergey Koren, Marcela Uliano-Silva, et al. 2021. “Towards Complete and Error-Free Genome Assemblies of All Vertebrate Species.” *Nature* 592 (7856): 737–46. https://doi.org/10.1038/s41586-021-03451-0.

Slabodnick, Mark M., J. Graham Ruby, Sarah B. Reiff, Estienne C. Swart, Sager Gosai, Sudhakaran Prabakaran, Ewa Witkowska, et al. 2017. “The Macronuclear Genome of Stentor Coeruleus Reveals Tiny Introns in a Giant Cell.” *Current Biology: CB* 27 (4): 569–75. https://doi.org/10.1016/j.cub.2016.12.057.

UniProt Consortium. 2021. “UniProt: The Universal Protein Knowledgebase in 2021.” *Nucleic Acids Research* 49 (D1): D480–89. https://doi.org/10.1093/nar/gkaa1100.

Xu, Peng, Jian Xu, Guangjian Liu, Lin Chen, Zhixiong Zhou, Wenzhu Peng, Yanliang Jiang, et al. 2019. “The Allotetraploid Origin and Asymmetrical Genome Evolution of the Common Carp Cyprinus Carpio.” *Nature Communications* 10 (1): 4625. https://doi.org/10.1038/s41467-019-12644-1.

Yang, Jinghua, Dongyuan Liu, Xiaowu Wang, Changmian Ji, Feng Cheng, Baoning Liu, Zhongyuan Hu, et al. 2016. “The Genome Sequence of Allopolyploid Brassica Juncea and Analysis of Differential Homoeolog Gene Expression Influencing Selection.” *Nature Genetics* 48 (10): 1225–32. https://doi.org/10.1038/ng.3657.

Yates, Andrew D., James Allen, Ridwan M. Amode, Andrey G. Azov, Matthieu Barba, Andrés Becerra, Jyothish Bhai, et al. 2022. “Ensembl Genomes 2022: An Expanding Genome Resource for Non-Vertebrates.” *Nucleic Acids Research* 50 (D1): D996–1003. https://doi.org/10.1093/nar/gkab1007.
